# Longitudinal Investigation of Neurobiological Changes Across Pregnancy

**DOI:** 10.1101/2024.03.08.584178

**Authors:** Yanbin Niu, Benjamin N. Conrad, M. Catalina Camacho, Sanjana Ravi, Hannah A. Piersiak, Lauren G. Bailes, Whitney Barnett, Mary Kate Manhard, David A. Cole, Ellen Wright Clayton, Sarah S. Osmundson, Seth A. Smith, Autumn Kujawa, Kathryn L. Humphreys

## Abstract

Pregnancy is a period of profound biological transformation. However, we know remarkably little about pregnancy-related brain changes. To address this gap, we chart longitudinal changes in brain structure during pregnancy and explore potential mechanisms driving these changes. Ten participants (Mean age = 28.97 years) are assessed 1–6 times (median = 3) during their pregnancy. Each visit includes anatomical and diffusion-weighted MRI, and assessments of waking salivary hormones, hair hormones, and inflammatory cytokines. Here we observe a reduction in gray matter volume gestational week, while neurite density index (NDI), a proxy of axon density, in white matter tracts increase across pregnancy. Progesterone levels are associated with reductions in brain volumetric measurements, and both progesterone and estradiol levels are linked to increases in NDI in white matter tracts. This study highlights the profound neurobiological changes experienced by pregnant individuals and provides insights into neuroplasticity in adulthood.

## Introduction

Pregnancy is a period characterized by profound changes, including hormonal surges, increased blood volume, heightened oxygen consumption, as well as morphological adjustments such as uterine expansion and relocation of organs^1–3^. These adaptations support fetal development and preparation for labor and delivery. The body’s response to pregnancy may also change the pregnant person’s brain, potentially as adaptive, nonadaptive, or merely evolutionary byproducts^4,5^. However, we know remarkably little about pregnancy-related brain changes. This knowledge gap regarding how, why, and to what extent the brain changes during pregnancy is, at least in part, attributable to historical reluctance of medical studies to include pregnant individuals due to concerns over potential risks and ethical complexities^6–9^. However, MRI has been used clinically among pregnant patients for the last four decades^10^, and numerous studies find no evidence that MRI is associated with increased risk for adverse effects on pregnant individuals or their offspring (e.g., congenital anomalies, neoplasm, or vision or hearing loss)^11–13^. Given that approximately 85% of US women give birth in their lifetimes^14^ and the existence of safe imaging procedures, it is critical to identify how a pregnant person’s brain changes across pregnancy.

Non-human animal studies of pregnancy report diminished hippocampal cell proliferation^15,16^ and reduced microglial cell density^17^, both of which may manifest as detectable reductions in gray matter volume. Research on structural brain changes associated with pregnancy in humans remains limited. While several studies suggest potential reductions in brain volume during pregnancy^18–22^, these findings, however, come with limitations. For example, only four participants were assessed repeatedly (3–7 times) during pregnancy in Oatridge et al.^21^, and anatomical scans were collected via a 1.0-T MRI system, which has relatively low signal-to-noise ratio (SNR) and spatial resolution. Further, neither Hoekzema et al.^18,19^, which compared women’s brains before conception to 2-3 months postpartum, nor Paternina-Die et al.^20^, which examined differences between pregnant individuals and nulliparous controls at late pregnancy and early postpartum, addressed changes occurring within pregnancy. More recently, Pritschet et al.^22^ leveraged precision imaging to map neuroanatomical changes in a single individual from preconception through two years postpartum, suggesting decreases in gray matter volume and cortical thickness across the brain during pregnancy. While this work offers intriguing evidence of remarkable neuroplasticity during pregnancy, the findings are based on a single subject, which limits the generalizability and scope of broader inferences. As such, the current literature regarding brain changes during pregnancy remains inconclusive. Within-subject longitudinal studies are essential for mapping brain dynamics throughout gestation, offering insights into pregnancy-related neuroplasticity.

In terms of potential pregnancy-related changes in white matter microstructure, non-human animal research has observed an increase in myelinated axons during pregnancy, potentially resulting from augmented proliferation of oligodendrocytes—a type of brain cell vital for axonal myelination^23–25^. In particular, Chan et al.^26^ reported a general increase in diffusivity across the whole brain during pregnancy, as well as regional increases in mean diffusivity (MD) and fractional anisotropy (FA) that were derived from Diffusion Tensor Imaging (DTI). The current human studies, however, have generated inconsistent findings. Hoekzema et al.^19^ found no significant changes in white matter diffusion metrics derived from DTI or in white matter volume when comparing pregnant women to nulliparous controls. In contrast, Pritschet et al.^22^ observed nonlinear increases in white matter quantitative anisotropy across the brain— indicating greater tract integrity—as gestational weeks progressed in a single subject.

In vivo longitudinal studies are crucial for detecting and understanding the dynamics of brain changes during pregnancy. Moreover, there are increasing calls for a more inclusive approach to biomedical research that recognizes and investigates the distinctive biological experiences of females^27–29^. Responding to this need, we initiated a pilot longitudinal study of pregnant people (*n* = 10, assessed 1–6 times [median = 3], spanning 12–39 gestational weeks). We used anatomical magnetic resonance imaging (MRI) to assess brain morphometry (e.g., gray matter volume and cortical thickness). White matter microstructure was evaluated with multi-shell diffusion-weighted MRI and advanced neurite orientation dispersion and density (NODDI)^30^ modeling, which provided metrics such as the neurite density index (NDI) and orientation dispersion index (ODI). The primary goals of the current study were twofold. The first objective was to chart the brain structural changes occurring during pregnancy. The second goal was to explore potential mechanisms driving these changes, particularly focusing on pregnancy hormones, inflammatory cytokines, and the intracranial cerebrospinal fluid (CSF) volume.

## Results

### Brain Changes during Pregnancy

#### Brain Morphometry Changes

Covarying for total intracranial volume (TIV) and age, total brain volume exhibited decreases across pregnancy (▱ = -0.105, 95% CI [-0.169, -0.040], *p* = .004). This translates to a 2.6% decline over the observed gestational range in this study (i.e., 12–39 weeks). We further conducted leave-one-out analyses, which resulted in estimated ▱ values that ranged from -0.184 to -0.090. Similarly, total gray matter volume exhibited decreases across pregnancy (▱ = -0.192, 95% CI [-0.301, -0.082], *p* = .002). This translates to a 3.9% decline over the observed gestational range in this study (see Fig. 1). With leave-one-out analyses, ▱ values ranged from -0.328 to -0.179. There were decreases in total cortical volume (▱ = -0.238, 95% CI [-0.374, -0.102], *p* = .002) and in subcortical volume (▱ = -0.094, 95% CI [-0.196, 0.008], *p* = .068), although the latter was not statistically significant. Additionally, there was a large decrease in cortical thickness (▱ = -0.516, 95% CI [-0.915, -0.116], *p* = .015), yet cortical surface area remained unchanged (▱ = -0.018, 95% CI [-0.052, 0.017], *p* = .297).

**Fig. 1.**
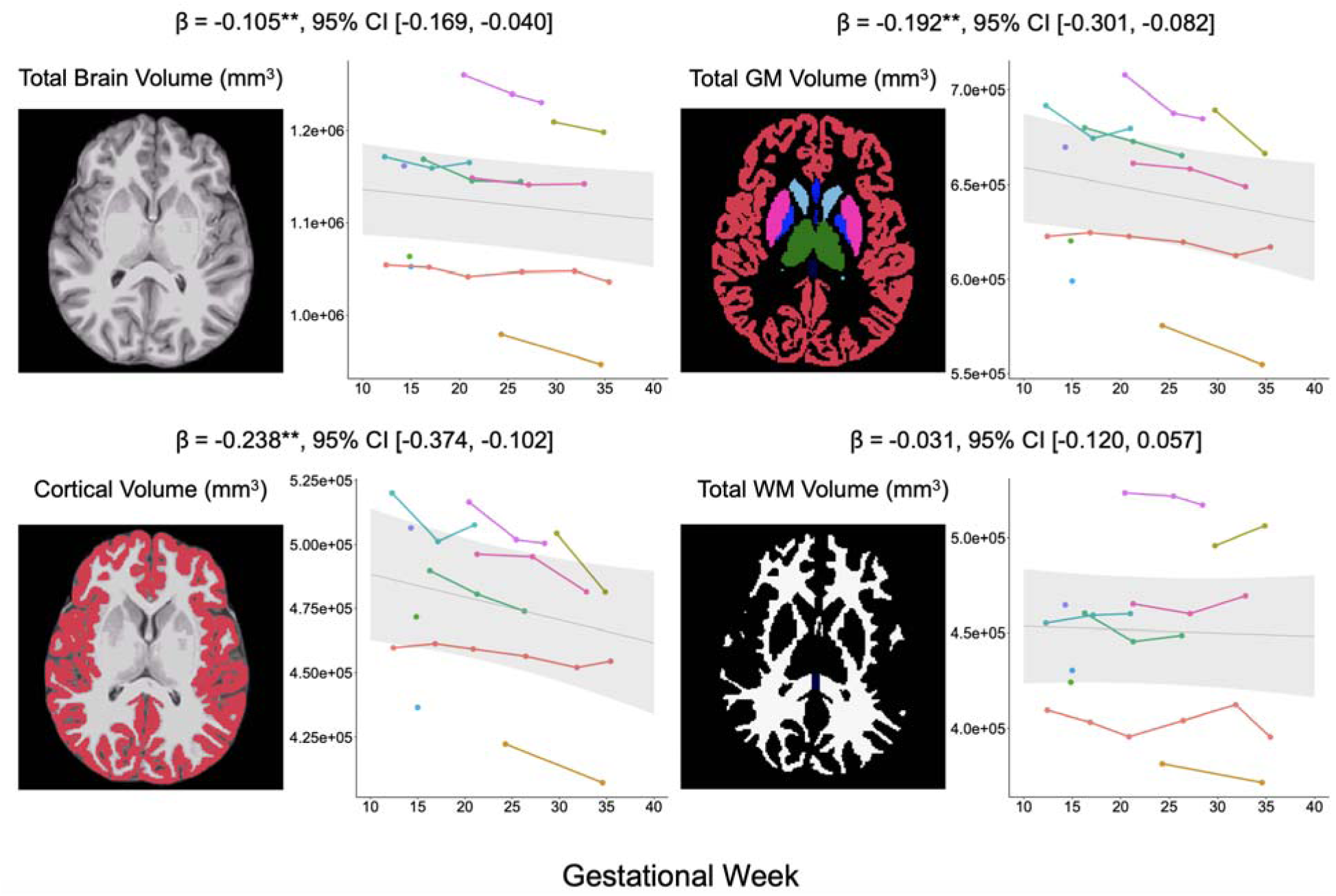
The observed decrease in brain volumes throughout the course of pregnancy. The plots are based on the linear mixed-effects model shown in Equation 1. Specifically, brain metrics were the dependent variable and gestational week was the predictor, covarying for age and total intracranial volume. GM: gray matter, WM: white matter. The shaded area represents the 95% confidence interval (95% CI) for the fitted line. **Note**. ** *p* < .01.

These results suggest that the observed reduction in total gray matter volume may primarily result from decreases in cortical volume, particularly cortical thickness (see Fig. S2 for detailed statistics). Lastly, total white matter volume did not exhibit a statistically significant change across pregnancy (▱ = -0.031, 95% CI [-0.120, 0.057], *p* = .460). All findings were consistent using leave-one-out analyses, as detailed in Table S2.

#### Changes in Cortical Regions

To further characterize the reduction in total cortical volume, we fitted linear mixed-effects (LME) models for all Desikan-Killiany atlas regions, using gestational week as the fixed effect and hemisphere, TIV, and age as covariates. A surface map with the standardized regression coefficients for all cortex regions is shown in Fig. 2a. Overall, we observed a consistent trend of decreasing cortical volume in most cortical regions, except for three regions in the occipital lobe (pericalcarine, lingual, and cuneus) and two regions in the medial temporal lobe (entorhinal and temporal pole). Statistically significant reductions in cortical volume were observed in prefrontal cortex regions, including the medial orbitofrontal, superior frontal, and lateral orbitofrontal cortex, as well as two regions surrounding the central sulcus: the precentral and postcentral cortex (Fig. 2b). Detailed statistical reports, including leave-one-out analyses consistent with the full sample results, are provided in Table S3.

**Fig. 2.**
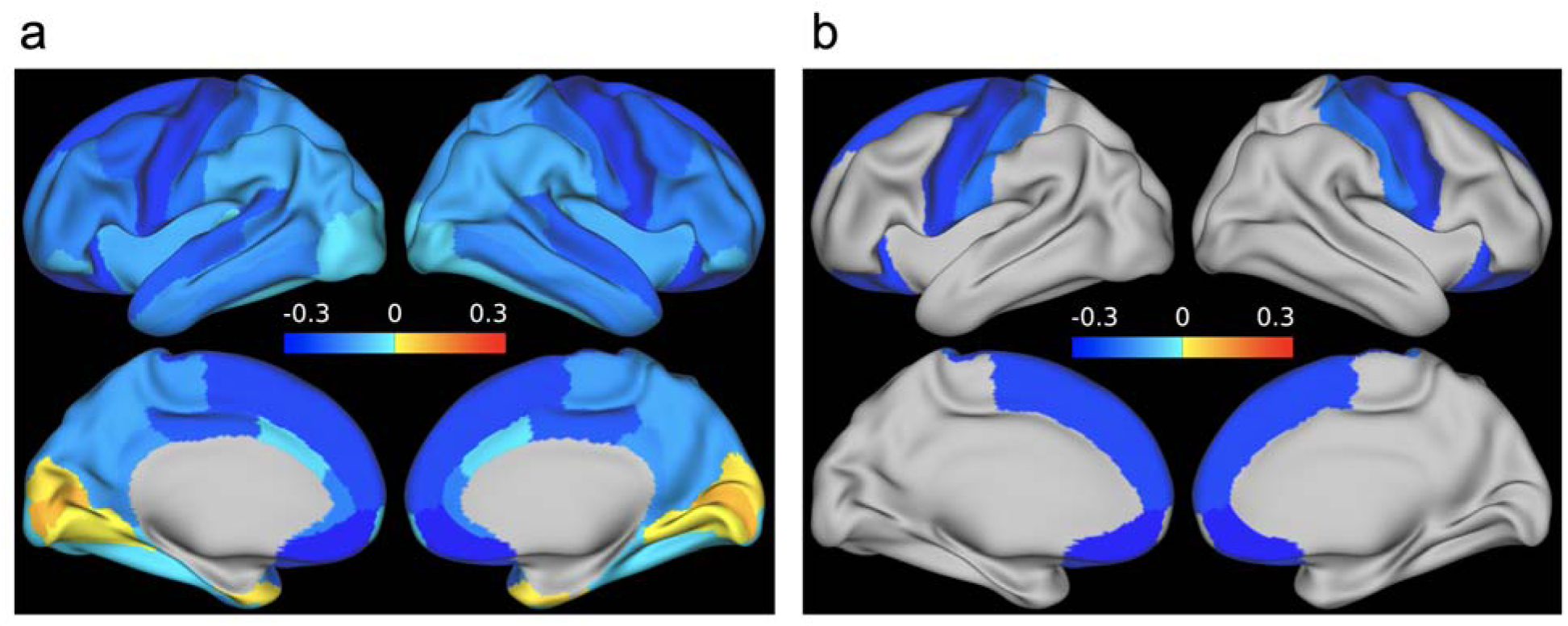
Illustration of the association between cortical region volumes, as defined by the Desikan-Killiany atlas, and gestational week. **a** color-coded to reflect the standardized regression coefficients derived from linear mixed effects models, which were fitted for each cortical region volume against gestational week while covarying for age and total intracranial volume (Equation 2). **b** selectively presents the cortical regions that exhibited a significant volume decrease throughout pregnancy. No multiple comparison correction was implemented.

#### White Matter Microstructure Changes

In our investigation of changes in white matter microstructure, we implemented LME models for each of the NODDI metrics (NDI and ODI) across each of the 50 tracts generated by TractSeg with gestational week as the fixed effect, and hemisphere (where applicable), age, and relative motion as covariates. To facilitate the understanding of the global pattern of metric changes, a heat map was generated, representing the standardized regression coefficients for all segmented tracts (see Fig. S3). Throughout the gestation period, we observed a predominant increase in NDI across most tracts, whereas ODI metrics remained relatively stable. Notably, the most pronounced increases in NDI were observed in projection tracts such as the corticospinal tract (CST), fronto-pontine tract (FPT), parieto-occipital pontine (PORT), in addition to the striato-premotor tract (ST_PREM) and arcuate fascicle (AF) (see Fig. 3). In line with these NDI findings, our analysis of DTI metrics revealed a predominant trend of increasing FA. Tracts that exhibited the most significant FA increases largely overlapped with those displaying notable NDI increases. Comprehensive statistical results for NDI and ODI metrics are available in Tables S6 and S7, and analyses of DTI metrics can also be found in the supplementary materials results section. Lastly, an examination of average measures across all segmented tracts revealed no significant global changes in NDI and ODI metrics during pregnancy, consistent with our secondary method for assessing global diffusion metrics (see supplementary materials results section).

**Fig. 3.**
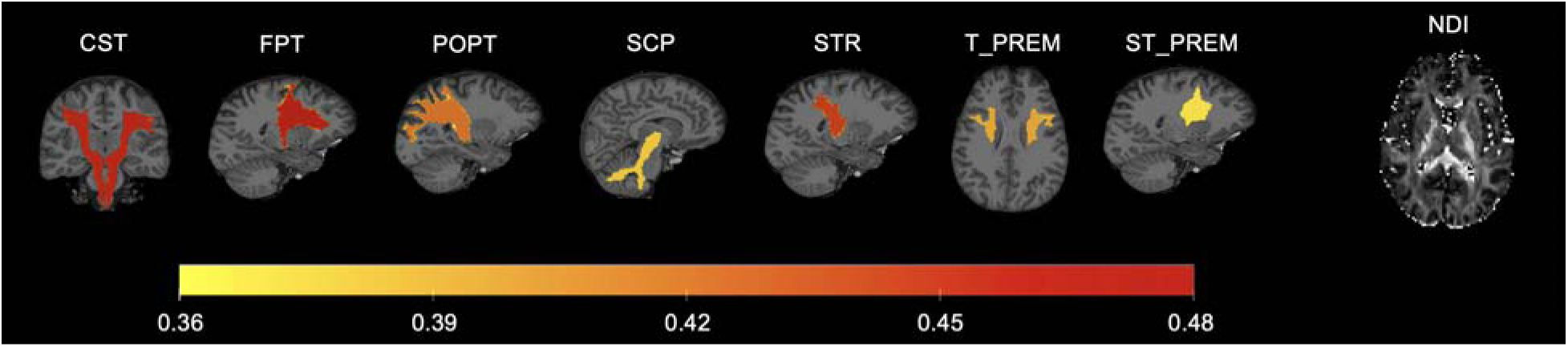
Illustration of the association between Neurite Density Index (NDI) and the progression of pregnancy. It was color-coded to represent the standardized regression coefficients from linear mixed effects models fitted for NDI against gestational week while covarying for age and relative motion (Equation 2). CST: Corticospinal Tract, FPT: Fronto-Pontine Tract, POPT: Parieto-Occipital Pontine, SCP: Superior Cerebellar Peduncle, STR: Superior Thalamic Radiation, T_PREM: Thalamo-Premotor, ST_PREM: Striato-Premotor.

#### CSF Changes

We observed a statistically significant increase in total intracranial CSF (▱ = 0.187, 95% CI [0.021, 0.353], *p* = .030) after covarying for age and TIV (see Fig. 4). Using a leave-one-out cross-validation approach, the ▱ values ranged from 0.145 to 0.267 (see Table S4). This pattern was consistent when varying the threshold of CSF volume fraction between 60% and 95% (see Table S5).

**Fig. 4.**
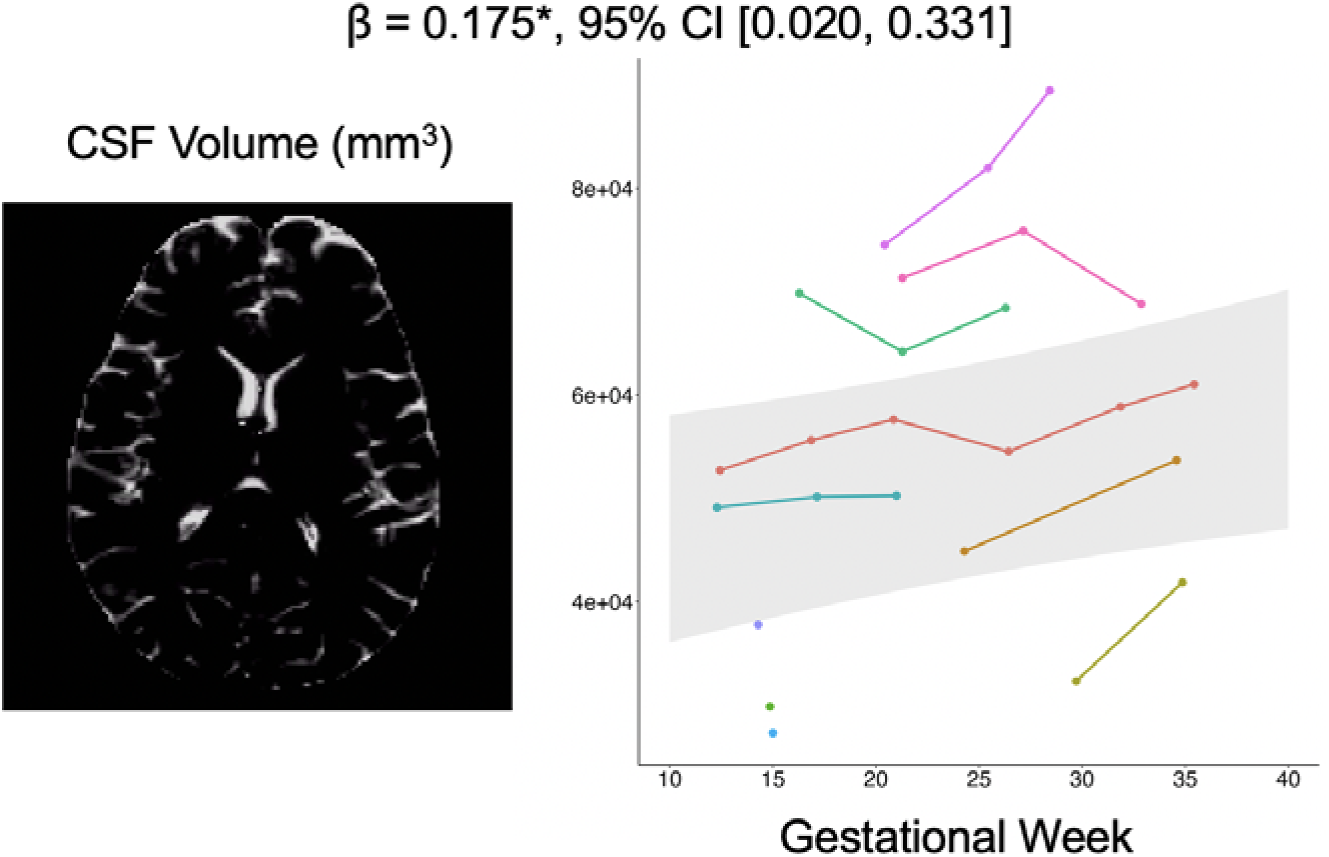
The increase in total intracranial cerebrospinal fluid (CSF) was observed across pregnancy. The plot is based on the linear mixed effects model shown in Equation 1. Specifically, CSF was the dependent variable and gestational week was the predictor, covarying for age and total intracranial volume. The shaded area represents the 95% confidence interval (95% CI) for the fitted line. **Note**. * *p* < .05.

### Associations Between Brain Dynamics and Salivary Hormones in Pregnancy

#### Salivary Hormones and Inflammatory Cytokine Changes during Pregnancy

Covarying for age, we observed statistically significant increases in waking salivary levels of progesterone (0= 0.717, 95% CI [0.391, 1.042], *p* < .001), estradiol (▱ = 0.792, 95% CI [0.533, 1.050], *p* < .001), and testosterone (▱ = 0.393, 95% CI [0.061, 0.724], *p* = .024) across pregnancy. However, the increase in cortisol levels was not statistically significant (▱ = 0.198, 95% CI [-0.014, 0.410], *p* = .065). Additionally, we did not observe statistically significant changes in C-reactive protein (CRP) levels throughout the course of pregnancy (▱ = -0.067, 95% CI [-0.248, 0.114], *p* = .441; see Fig. 5a).

**Fig. 5.**
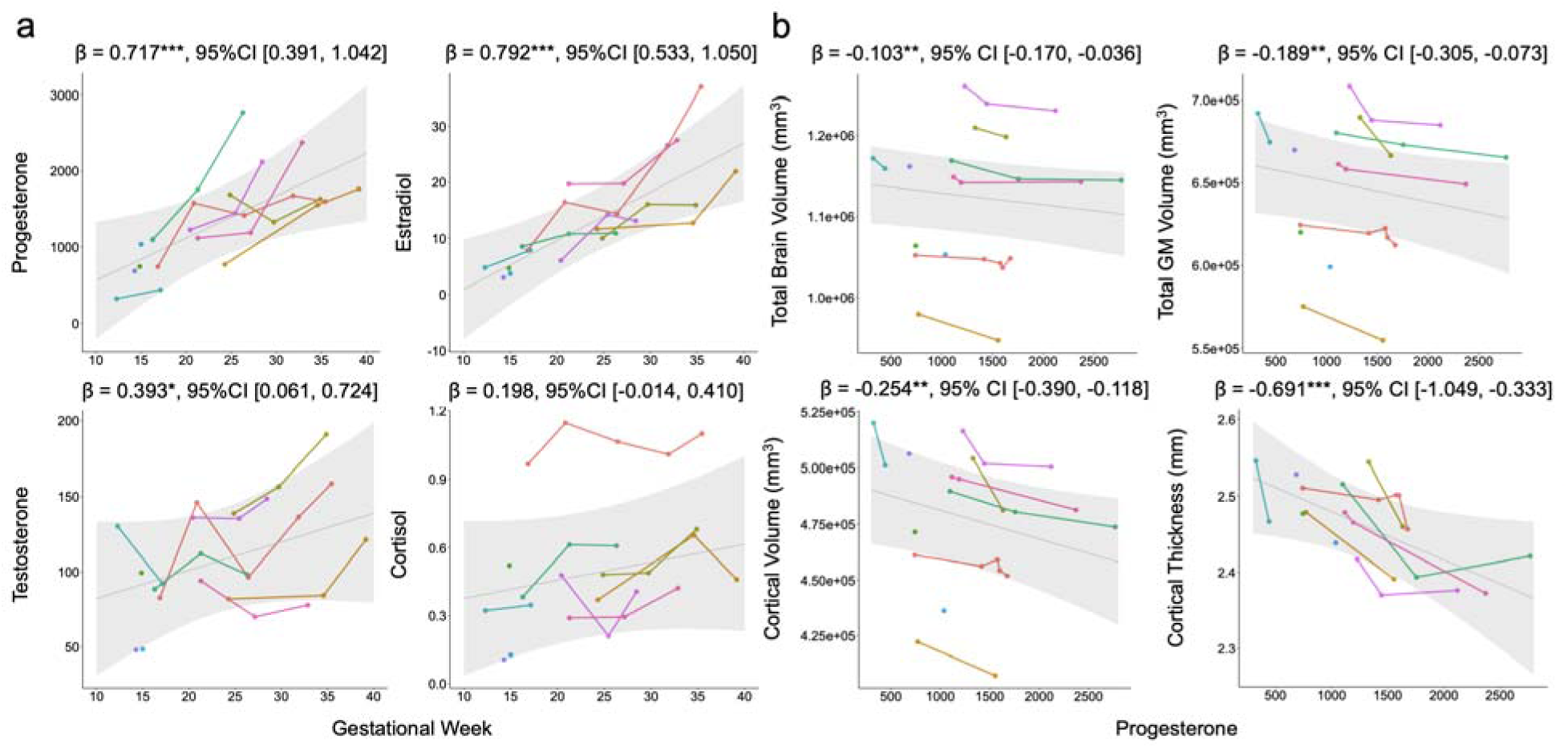
The observed changes in hormone levels during pregnancy and associations between progesterone levels and brain structural metrics. **a** The observed changes in hormone levels (progesterone, estradiol, testosterone, and cortisol) during pregnancy. Linear mixed effects models were applied with hormone levels as the dependent variable and gestational week as the predictor, covarying for age. **b** Associations between progesterone levels and brain structural metrics, including total brain volume, total gray matter volume, total cortical volume, and cortical thickness. The plot is based on Equation 3. Specifically, brain metrics were the dependent variable and progesterone was the predictor, covarying for age and total intracranial volume. The shaded area represents the 95% confidence interval (95% CI) for the fitted line. **Note**. ** *p* < .01. *** *p* < .001.

#### Associations Between Salivary Hormones and Brain Structure

After covarying for TIV and age, we found statistically significant negative associations between total brain volume and salivary levels of progesterone (▱ = -0.103, 95% CI [-0.170, -0.036], *p* = .006) and cortisol (▱ = -0.213, 95% CI [-0.393, -0.032], *p* = .025), indicating that increasing salivary levels of progesterone and cortisol are associated with decreasing total brain volume. Associations between total brain volume and either salivary estradiol or testosterone levels were not statistically significant (*p*s > .144). Additionally, total gray matter volume was negatively associated with salivary progesterone (▱ = -0.189, 95% CI [-0.305, -0.073], *p* = .003), but not with salivary estradiol, testosterone, or cortisol (all *p*s > .134). Total cortical volume (▱ = -0.254, 95% CI [-0.390, -0.118], *p* = .002) and mean cortical thickness (▱ = -0.691, 95% CI [-1.049, -0.333], *p* = .001) were also negatively associated with salivary progesterone (see Fig. 5b), but not with salivary estradiol, testosterone, and cortisol levels (all *p*s > .137).

#### Associations Between Salivary Hormones and Diffusion Metrics

Fig. S8 shows the standardized regression coefficients of hormone levels across all segmented tracts, illustrating the associations between NDI and salivary hormone levels. Most notably, our results indicated that NDI across nearly all tracts was positively associated with salivary estradiol levels and moderately with salivary progesterone (see Fig. 6). Moreover, NDI demonstrated a negative link with salivary cortisol levels, while no significant association was found with salivary testosterone. Detailed statistical outcomes of these analyses are available in Table S12–S15.

**Fig. 6.**
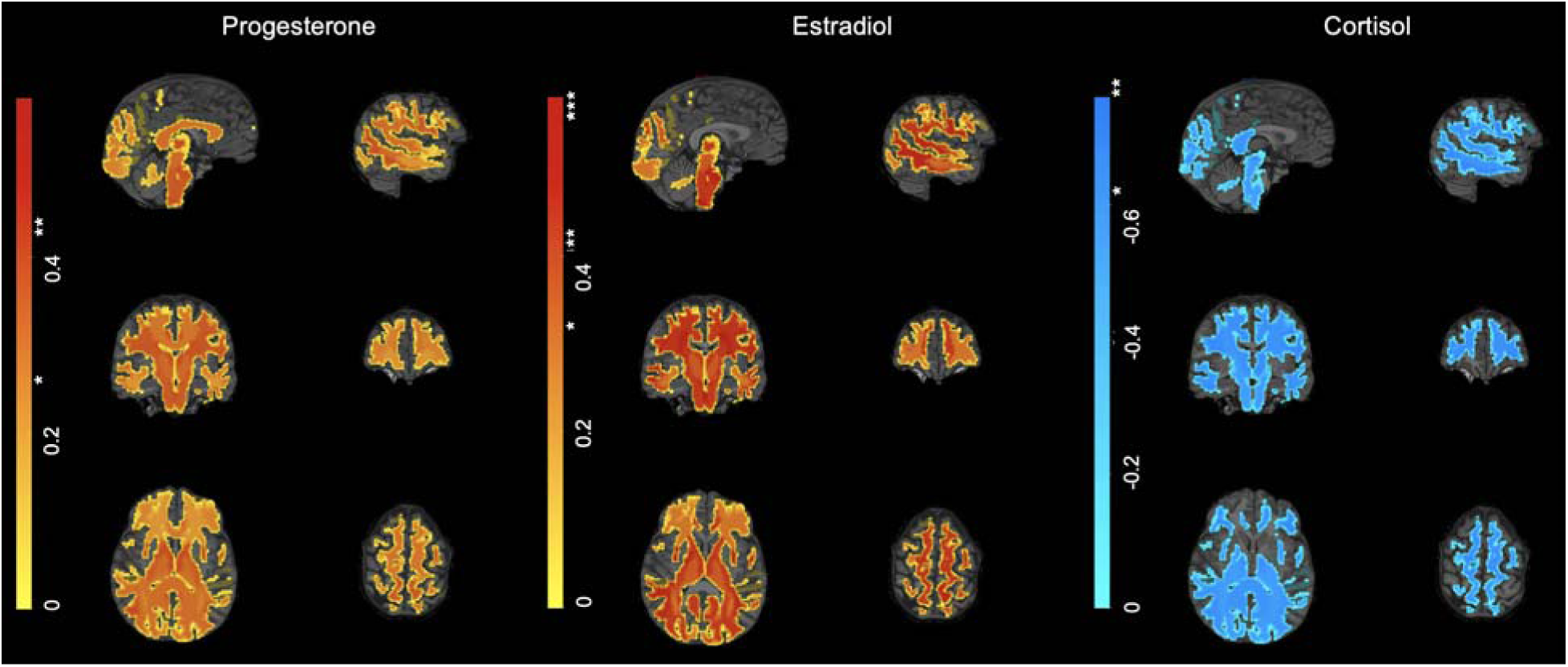
The association between Neurite Density Index (NDI) and hormone levels (specifically progesterone, estradiol, and cortisol). The color-coding reflects the standardized regression coefficients obtained from linear mixed effects models, which were used to analyze the relation between NDI and each of the three hormones while covarying for age and relative motion (Equation 4). **Note**. * *p* < .05. ** *p* < .01. *** *p* < .001. An asterisk in the color bar indicates the level of significance for standardized regression coefficients.

#### Associations Between Salivary Hormones and CSF

Covarying for TIV and age, no statistically significant associations were detected between CSF volume changes and salivary hormonal levels, including progesterone (▱ = 0.130, 95% CI [-0.060, 0.319], *p* = .160), estradiol (▱ = 0.151, 95% CI [-0.060, 0.361], *p* = .143), testosterone (▱ = 0.153, 95% CI [-0.119, 0.425], *p* = .242), and cortisol (▱ = 0.187, 95% CI [-0.284, 0.668], *p* = .401; see Table S16–S19).

#### Associations Between CSF and Brain Structure

There were statistically significant negative associations between total intracranial CSF and total brain volume (▱ = -0.277, 95% CI [-0.452, -0.102], *p* = .004), total gray matter volume (▱ = -0.388, 95% CI [-0.678, -0.097], *p* = .013), and total cortical volume (▱ = -0.391, 95% CI [-0.754, -0.028], *p* = .037), suggesting that reductions in brain volumetrics may be associated with the increases in the total intracranial CSF volume. However, the link between total intracranial CSF and mean cortical thickness was not statistically significant (▱ = -0.515, 95% CI [-1.069, 0.039], *p* = .066).

#### Associations Between CSF and Diffusion Metrics

Across all segmented tracts, there were no statistically significant associations between total intracranial CSF and NDI. Detailed statistical outcomes of these analyses are available in Table S20.

## Discussion

In a sample of 10 pregnant participants who underwent multimodal MRI scans, hormone assessments, and inflammatory cytokine assessments between 1 and 6 times during pregnancy (median = 3 times), we explored whether, and to what degree, neurobiological changes occur during pregnancy. There were four main findings. First, there were pronounced reductions in overall brain and gray matter volume, while an increase was found in the total intracranial CSF volume. Second, using NODDI metrics to assess white matter microstructure, we found an increase in NDI across the majority of segmented tracts, with the most pronounced increase observed in tracts associated with sensorimotor processing. Third, there were significant correlations between pregnancy-related salivary hormones and observed brain changes (i.e., volumetric reduction in overall brain and gray matter, and increased NDI in segmented tracts). Fourth, the increase in the total intracranial CSF volume was associated with the decrease in brain volumetrics, but not with NDI changes.

The decreases in estimates of total brain volume in the present study were consistent with earlier evidence involving both humans^18–22^ and non-human animal models^15–17^. Comparing mothers before and after their first pregnancy, Hoekzema et al.^18,19^ reported a reduction in gray matter volume across multiple brain regions, notably within the anterior and posterior midline structures. Another recent study found that, during late pregnancy, mothers exhibited smaller global cortical volume and thickness compared to nulliparous controls, with the most pronounced differences in the somatomotor, attentional, and default mode networks^20^. A recent study by Pritschet et al.^22^ focused on neuroanatomical changes from preconception to two years postpartum in a single individual, documenting significant reductions in gray matter volume and cortical thickness. Non-human animal studies suggest that the observed brain volume reduction may be linked to decreased microglial proliferation, as evidenced by the reduced microglial cell density in pregnant rats during late gestation and the early postpartum period compared to virgin rats^17^. Expanding upon previous pre- and post-pregnancy comparisons and cross-sectional analyses, our findings demonstrate a significant rate of reduction in gray matter volume during pregnancy. Additionally, we observed a significant correlation between CSF volume and overall brain and gray matter volumes. This association suggests that intracranial CSF volume increases during pregnancy, consistent with the prior report of ventricular volume expansion^21^, may exert pressure on the brain and compress the cortex, possibly contributing to the observed decreases in brain volume^31,32^.

Additionally, we observed an increase in NDI across tracts, especially those linked with sensorimotor processing, whereas ODI remained largely unchanged. NDI quantifies the tissue volume fraction occupied by neurites, such as axons and dendrites, and is considered a proxy for axonal density in white matter^33,34^. Meanwhile, ODI reflects the variance in neurite orientation. Our findings suggest that across pregnancy there may be an increase in axonal density, potentially stemming from increased myelination and/or axonal diameter, without significant shifts in white matter geometric complexity. This aligns with prior non-human animal studies that have documented an increase in myelinated axons during pregnancy, potentially as a result of amplified proliferation of oligodendrocytes^24,25^. However, there is a discrepancy between our results and those reported by Hoekzema et al.^19^ who did not observe significant changes in DTI metrics during pregnancy. The discrepancy may arise from differences in imaging sequence parameters. For instance, Hoekzema et al.^19^ used single shell protocol (b = 1000) in contrast to the multi-shell protocol (b = 500, 1500, 2500) in the present study, which has been shown to affect the estimated scalar diffusion metrics^35^. Another consideration is the methodological divergence between the two studies. Our study adopted a longitudinal design, capturing dynamic changes over the course of pregnancy, while Hoekzema et al.^19^ employed a pre-post pregnancy comparison approach. Their post-pregnancy data acquisition took place, on average,

100 days after childbirth coupled with a wide variability (*SD* = 70 days). This extended interval could potentially obscure white matter alterations occurring during pregnancy. Indeed, the recent study by Pritschet et al.^22^ demonstrated an increase in global quantitative anisotropy (QA), an index of white matter microstructural integrity, during gestation, which was concomitant with rises in estradiol and progesterone. However, extended analyses into the postpartum period revealed non-linear patterns of global QA, with increases during the first and second trimesters, returning to baseline levels postpartum. Thus, it is possible that white matter quickly reverts to its pre-pregnancy state, thereby remaining undetected in the delayed post-pregnancy assessment. Alternatively, the postpartum period itself, characterized by hormonal fluctuations, physiological adaptations, and new caregiving demands, might induce its own set of white matter microstructural adaptations, which might counteract those taking place during pregnancy. Additionally, the absence of global changes in white matter microstructural metrics, specifically NDI, may be due to regional variability. While we observed increases in NDI across most tracts, other tracts exhibited small to subtle increases. As such, the effects may not be detectable at the global level. Although it is also possible that our study is underpowered to detect subtle global changes that may indeed be present, this observation, if generalized, highlights the importance of understanding focal tract changes in NDI that occur during pregnancy in future studies.

The current study also explored candidate mechanisms underlying the observed neurobiological changes during pregnancy. First, our data revealed a correlation between pregnancy-associated hormones and brain alterations in both gray matter and white matter. Estradiol and progesterone are pivotal neuroprotective agents, modulating processes like neurogenesis^36,37^, dendritic spine formation and branching^38,39^, myelination^40,41^, axonal growth, and synaptogenesis^42^. Our findings of positive correlations between white matter NDI and progesterone and estradiol levels are consistent with such findings from animal studies.

However, the observed inverse associations between pregnancy-related hormones and gray matter volumes are somewhat counterintuitive, especially when considering that non-human animal research suggests estradiol and progesterone promote neurogenesis^43,44^. It might be that increased dendritic length, branching, and spine density in gray matter due to pregnancy-related hormones^38,38,39,42,45^ and/or myelination lead to misclassification of voxels at tissue interfaces, thereby appearing as reductions in gray matter volume^46,47^, either in addition to or instead of fluid-related compression. It is also possible that hormones influence other physiological functions that are linked to a decrease in gray matter volume. For example, suppressed microglial proliferation might underlie the connections between hormonal shifts and grey matter reduction during pregnancy^47^. Furthermore, the observed inverse associations raise the possibility of a non-linear, U-shaped relationship, where progesterone’s links to brain structure may not be straightforward and could vary depending on concentration levels^48^. Indeed, inverted U-shaped relationships have also been observed for other hormones such as estradiol^49^ and cortisol^50^ in humans.

Additionally, it is interesting that we observed a link between brain morphometry and progesterone, but not estradiol. However, we should acknowledge that salivary hormone assays can be affected by numerous factors, including collection time and sample storage^51,52^. If these effects generalize, it might suggest that progesterone and estradiol have distinct links to brain morphometry. This distinction has been evidenced by their different effects on receptors^53^ and functional network organization^54^, as well as their complex interactions in synaptic plasticity and neuroprotection^55^. It has been reported that intrinsic fluctuations in progesterone, but not estradiol, were associated with medial temporal lobe morphology across the human menstrual cycle^56^. In mice, during pregnancy, both estradiol and progesterone affect the galanin (Gal)- expressing neurons in the mouse medial preoptic area (MPOA), but in different ways—estradiol silences MPOA^Gal^ neurons and increases neuronal excitability, while progesterone promotes dendritic spine formation. Those effects on MPOA^Gal^ neurons are linked to increased selectivity for pup stimuli^57^. However, whether these previously mentioned findings from animal studies translate to humans remains unclear, given the challenges inherent in comparing animal models to human experiences^58^. It is important to note that our findings reflect associations and cannot be interpreted as evidence of causality. The observed relations between hormone levels and brain structure could be due to a variety of factors, such as underlying physiological changes that occur during pregnancy and/or the influence of other biological processes not measured in the current study.

Moreover, the influence of pregnancy-related hormones likely represents only one of the factors contributing to observed brain changes. Other pregnancy-associated alterations, such as increased water content^59,60^ and blood flow variations^61,62^ during pregnancy, may also play significant roles. For example, studies have documented a significant and progressive rise in total body water, as well as extracellular and intracellular water values throughout pregnancy^59,60^, which might help explain the observed elevation in NDI. Yet, it is important to note that in vivo MRI methods offer insights at a macroscopic level and inherently lack the resolution to elucidate the cellular and molecular mechanisms at play during pregnancy. Future research across multiple levels of analysis is needed to supplement this work and enhance our understanding of the neurobiological changes observed during pregnancy.

Although cortisol levels in our study did not reach statistical significance at the .05 level, they demonstrated a small to moderate increasing trend over the course of pregnancy. This observation is consistent with previous literature indicating that salivary cortisol levels progressively rise during pregnancy^63,64^. Additionally, we observed negative associations between cortisol levels and total brain volume, as well as between cortisol levels and white matter NDI. These findings align with existing research that implicates cortisol in neurodevelopmental processes, including neurogenesis^65,66^, synaptic plasticity^67,68^, and myelination^69^. Higher cortisol levels have been linked to reduced brain volume and alterations in white matter microstructures^70^. Therefore, despite the lack of statistical significance in the observed cortisol changes, the upward trend might still hold biological relevance in influencing brain structure in pregnant individuals. However, it is important to acknowledge the limitations of the waking salivary cortisol assay in our study, as it was assessed only once upon waking^71,72^, and hormone assays can be affected by various factors, as mentioned above. As such, interpretations should be made with caution. Future research may benefit from more frequent cortisol measurements to better capture dynamic changes during pregnancy and explore their implications for brain structure.

Additionally, we did not observe a significant change in CRP levels during the observed period (12 to 39 gestational weeks). This finding aligns with previous literature, which demonstrates that CRP levels can vary widely among individuals during pregnancy—some studies report an increase^73–75^, some report consistency^73,76,77^, and others report a decrease^73,74^ in CRP levels across gestation. Therefore, while CRP is an important marker of inflammation^78^, it has not been consistently shown to track with gestation and is likely influenced by various factors.

While not the focus of the current study, there are multiple hypotheses regarding the potential implications of pregnancy-related brain changes. While some contend that changes to the pregnant person’s brain are most likely byproducts of promoting fetal survival and childbirth^4^, others point to the possibility that these changes are in the service of promoting the parent–child relationship^19,79,80^. There is substantial evidence that interactions with children post-birth shape the brain. Indeed, evidence from non-birthing parents, which allows for distinctions between pregnancy-specific vs. caregiving-related brain changes, shows brain alterations following childbirth^81–84^ and child rearing^85,86^. Additionally, research has shown that brain changes can persist up to six years postpartum^87^, and persistent changes may relate to the cognitive and emotional challenges of caring for a child^88,89^. Future work would benefit from careful consideration of the potential functions of these changes, with designs that help to rule out plausible rival hypotheses.

Several limitations merit consideration. First, our study was constrained by a small sample size. The power to detect subtle effects is therefore limited as well as our ability to adjust for potential confounds and examine the potential effect of moderators. For example, prior research suggests that the number of live births is related to brain age^88^. It is possible that the number of live births might moderate the neurobiological changes during pregnancy, such that individuals with more births experience less (or more) neurobiological changes (see supplementary materials results section for our exploration of these associations within our sample). Given that the current literature has focused nearly exclusively on first-time mothers, it is especially important for future research to consider the number of prior births on pregnancy-related brain changes. Second, our study is limited by the number of data points available (27 unique scan sessions from 10 individuals), with some participants attending only a single session. For example, while our calculation of percentage declines in brain morphometry using estimated LME models offers a useful estimate of average changes during pregnancy, it is important to interpret these data with caution. The focus on the observed range of gestational weeks (i.e., 12–39 weeks) is intentional to accommodate the limited coverage of gestational weeks and small sample size, which could nevertheless affect the precision of this estimate.

Third, our study does not have benchmarking data. Including such comparisons in future research would provide additional context for interpreting the uniqueness of pregnancy-related neurobiological changes. For example, benchmarking against non-birthing parents could help isolate changes unique to pregnancy from those due to shared environmental or experiential factors, offering a more nuanced understanding of the impacts of pregnancy. Fourth, our sample did not cover the entire duration of pregnancy. Without preconception data, we have an incomplete picture of when during pregnancy such alterations become evident. Fifth, given the preliminary nature of our investigation, we opted not to implement corrections for multiple comparisons. Although this decision enhances our sensitivity to potential effects, it simultaneously raises the risk of type I errors. However, the consistency in patterns and lack of randomness suggest that the results may not be artifacts but are likely indicative of genuine underlying phenomena. It is important for subsequent, larger-scale studies to incorporate appropriate corrections to further validate and replicate our observations. Sixth, our exploration of the underlying mechanisms of neural dynamics primarily focused on hormonal fluctuations during pregnancy. Although these hormonal shifts are important, they might represent just one aspect of a more multifaceted interplay. Other potential factors, such as metabolic changes, vascular dynamics, and alterations in neurotransmitter levels, may also significantly contribute. Future research adopting an interdisciplinary approach, integrating insights from these domains would shed light on the multifaceted nature of brain changes during pregnancy. Lastly, careful monitoring of pregnancy complications (e.g., preeclampsia and eclampsia) which may significantly affect brain structure^21^, is crucial to ensure accurate interpretation of the findings in the future.

The present study elucidates neurobiological changes occurring across pregnancy. Specifically, we observed a decline in total brain and gray matter volume, along with increases in total intracranial CSF volume and NDI in white matter tracts across pregnancy. Our data also suggest that increases in CSF and pregnancy-related hormones may, at least partially, contribute to these observed neural alterations. These findings highlight the neuroplasticity associated with pregnancy, establishing a foundation for further investigations in this essential yet understudied area of research.

## Methods

### Participants

A total of 10 pregnant women aged between 20.00 and 39.25 years (*M* = 28.97, *SD* = 6.50) years were recruited for the current longitudinal study. In terms of race, 7 identified as White, 2 as Black or African American, and 1 selected “Other.” In terms of ethnicity, 1 participant identified as Hispanic/Latine. Participants were recruited through online advertisements and referral/word of mouth and 1 participant was the senior author. Participant eligibility was initially screened via telephone. Inclusion criteria required participants to be pregnant, at least 18 years old, English-proficient, and have no immediate plans to relocate. Exclusion criteria encompassed a medical history of neurological injury or impairment, a history of psychosis, non-removable metal or electronic implants, having undergone in vitro fertilization (IVF), and use of medication(s) that could potentially influence hormonal or neurological changes during pregnancy (e.g., progesterone). The mean number of prior pregnancies carried to term was 0.50 (*SD* = 0.71, range = 0–2). The mean weeks gestation at data collection was 23.90 (*SD* = 7.70, range = 12.29–39.14). The number of data collection sessions for each participant ranged from 1 and 6 times (mean number of repeated measures = 2.70 times, *SD* = 1.49), contingent on individual circumstances and pregnancy progression. Fig. 7 provides a detailed illustration of the gestational timing range captured for each participant. Each session included an MRI scan and collection of hair samples and dried blood spots. Additionally, participants were asked to collect saliva samples at their homes on the morning prior to and the morning of each session. All participants provided written informed consent prior to participation. Study procedures were approved by the Vanderbilt University Institutional Review Board. All ethical regulations relevant to human research participants were followed.

**Fig. 7.**
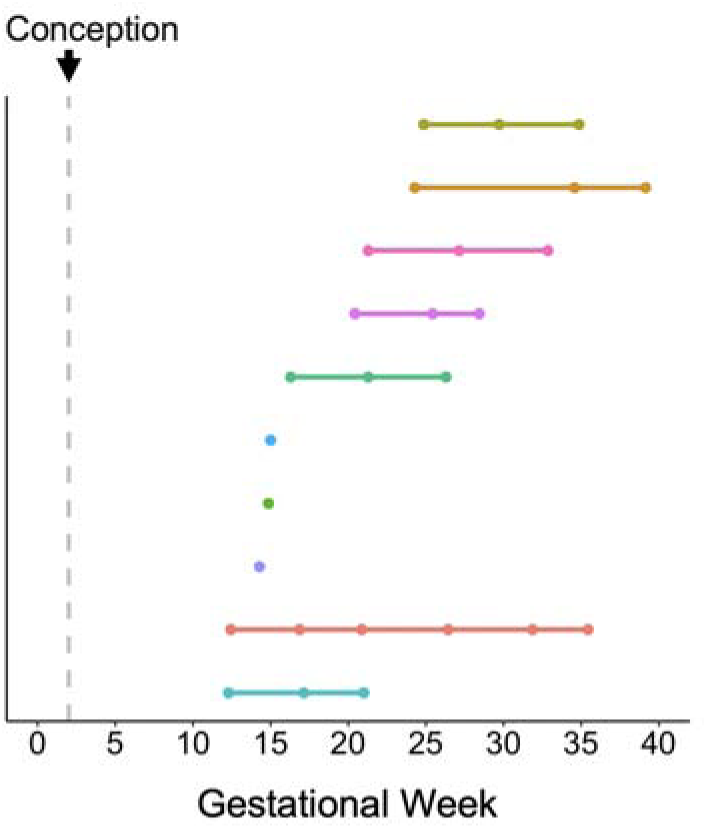
Illustration of the gestational weeks range captured for participants. Zero on the x-axis denotes the first day of the last menstrual period, with conception following approximately two weeks later.

### Hormone Assessments

Hormonal assessments were performed by measuring levels of progesterone, estradiol, testosterone, and cortisol in saliva samples obtained from participants, in accordance with Salimetrics collection and assaying procedures. Two saliva samples were collected for each participant: one on the morning prior to the MRI scan and another on the day of the scan. The obtained values were then averaged for analysis. Additionally, we obtained estimates of cortisol, cortisone, and progesterone from hair samples. Further details on the saliva and hair sample collection and assay procedures can be found in the supplementary materials sections titled Salivary hormone assessments and Hair hormone assessments. We primarily reported on salivary hormones, and results from hair hormone analysis were included in the supplementary materials results section.

### Inflammatory Cytokines Assessments

Inflammatory cytokine assessments were performed by measuring levels of Interleukin-6 (IL-6), Interleukin-10 (IL-10), Tumor Necrosis Factor alpha (TNF□), and CRP in dried blood spots (DBS) obtained from participants. At each visit, up to 5 drops of capillary whole blood were collected on filter paper (Whatman 903) following a finger stick with a sterile microlancet. Blood was then dried at room temperature and stored at -20 degrees Celsius in accordance with protocols established by Northwestern University’s Laboratory for Human Biology Research^90,91^, which performed assays on DBS samples. Further details on the DBS collection and assay procedures can be found in the supplementary materials section titled Inflammatory cytokines assessments. The usability of the samples for IL-6, IL-10, and TNF□ was constrained by the high volume requirements, resulting in a low yield of successfully assessed data for these cytokines (less than 50%). Thus, only CRP was utilized in the analyses. Detailed descriptive statistics are provided in Supplementary Table S1.

### MRI Acquisition

MRI data acquisition was conducted on a 3T Philips Achieva scanner using a 32-channel head coil at Vanderbilt University Institute of Imaging Science. High-resolution T1- and T2-weighted images were obtained for each participant through 3D-quantification using an interleaved Look–Locker acquisition sequence with a T2 preparation pulse (3D-QALAS) sequence^92^ with the following parameters: repetition time (TR) of 5.0 ms, echo time (TE) of 2.3 ms, a flip angle of 4 degrees, a field of view (FOV) of 230 x 193 x 150 mm, a matrix size of 224 x 224 x 146, and a voxel size of 1 x 1 x 1 mm. The acquisition parameters of diffusion-weighted imaging (DWI) were as follows: TR of 4300 ms, TE of 91.5 ms, flip angle of 90 degrees, FOV of 240 x 240 x 135 mm, matrix size of 112 x 112 x 63, yielding a voxel size of 2 x 2 x 2 mm. The diffusion weighting was distributed across 23 directions with a b-value of 500 s/mm², 47 directions with a b-value of 1500 s/mm², and 70 directions with a b-value of 2500 s/mm². The scanner used in this study underwent regular calibrations and quality control assessments at least weekly, conducted by the Vanderbilt University Institute of Imaging Science. Additionally, vendor engineers performed quarterly preventative maintenance and periodic image quality tests to assess system performance across various specifications, including SNR, geometry, scale, and uniformity. Furthermore, the scanner was remotely monitored by Philips, with daily performance reports covering parameters such as cryogenic system stability, noise levels, coil functionality, hardware integrity, cooling efficiency, and environmental factors like temperature and humidity. Throughout the study, all reports and test results indicated no significant deviations in scanner performance. Further details can be found in the supplementary materials section titled MRI acquisitions.

### MRI Processing

#### Anatomical MRI

Synthetic T1-weighted and T2-weighted images were found from 3D-QALAS images using SyntheticMR software (version 0.45.35; Linkoping, Sweden). Both T1-weighted and T2-weighted anatomical images were then used to extract structural MRI data through FreeSurfer v7.1.0’s longitudinal pipeline^93^. Scans at every time point were independently processed for each participant (including skull stripping, Talairach transformation, subcortical structure labeling, surface extraction, spherical registration, and cortical parcellation), followed by the creation of an unbiased within-subject template space using all the subject’s time points^94^.

Note that the FreeSurfer longitudinal pipeline assumes a fixed head size over time, meaning that TIV is considered constant across time in the final longitudinal output. By accounting for the expected stability, the longitudinal pipeline is particularly sensitive to detecting small, gradual changes over time. To ensure that TIV remained relatively stable and to better support the inferences made from our subsequent morphological analyses, we used the cross-sectional estimates, treating each dataset as a separate scan before feeding it into the longitudinal pipeline. The results showed that total TIV did not exhibit a meaningful or statistically significant change over the course of pregnancy (Δ = -0.014, 95% CI [-0.060, 0.033], *p* = .539; see Fig. S1). Following the automated processing steps, all the cortical reconstructions were visually inspected for potential inaccuracies. We observed no processing failures and no manual edits were required. Please refer to the supplementary materials for the Euler number report and example anatomical images in the section titled Anatomical MRI. Subsequently, we obtained measurements of volume for 34 cortical regions in both the left and right hemispheres (68 regions in total) using FreeSurfer’s cortical parcellation based on the Desikan-Killiany atlas^95^, which were used for cortical morphometric analyses.

### CSF

To quantify the total CSF volume in the entire brain, we used the aggregated volume fraction of the CSF image generated by SyntheticMR software from the 3D-QALAS images. Specifically, we first applied a threshold to retain voxels in which the CSF volume faction exceeded this threshold. Then, we multiplied the CSF volume fraction by the voxel volume across the brain. For our primary analyses, we set a threshold at 80%, meaning that a voxel was classified as containing CSF if it has 80% or more CSF. To test if this threshold influenced findings, we repeated analyses using different thresholds across a range of 60% to 95% (reported in Table S4). We observed consistent results across these different probability thresholds.

### Diffusion-weighted MRI

The raw diffusion data were processed using MRtrix3^96^ and FMRIB’s Software Library (FSL)^97^. Preprocessing steps involved noise estimation and removal via the dwidenoise function from MRtrix3^98,99^. The eddy and susceptibility distortion corrections were performed using the dwifslpreproc function in MRtrix3^97,100–102^. Subsequent to preprocessing, we computed the NDI and ODI using the NODDI Matlab Toolbox^30^. All preprocessing steps and NODDI metrics were carried out in native space. For comparison, we also fitted the standard DTI model^103^. In order to analyze changes across tracts, we first estimated fiber orientation distributions (FOD) using MRtrix3’s dwi2fod msmt_csd and then normalized them using MRtrix3’s mtnormalise^104,105^. The normalized FODs were then fed to TractSeg to automatically segment fiber tracts, resulting in NODDI and DTI metrics for each tract across 50 established tracts, which were subsequently used in our statistical analyses^106,107^. Lastly, we characterized the global changes for each NODDI metric and investigated if observed tract-level changes reflected a general alteration across all segmented tracts using two different approaches. For a more detailed description of diffusion data processing and metric quantification, please refer to the supplementary materials section titled Diffusion-weighted MRI.

#### Statistics and reproducibility

Given the longitudinal nature of our data, analyses were conducted using LME models, utilizing the “lme4” package in R^108^. One of the main advantages of using LME models with longitudinal data is their flexibility and robustness in handling missing data points and designs with variable numbers and intervals between measurements^109^. This approach can weigh the data points based on the number of observations per subject, thus subjects with different numbers of observations can be included without biasing the overall estimates, enabling us to use all available data points^110^. Note that we included all available time points, including those from subjects with only a single time point, to maximize data usage.

While individuals with data at only a single-time point do not contribute to the estimation of within-subject changes, they provide information about between-subject variability, and inclusion results in improved intercept estimation and overall model accuracy^111^.

We organized analyses into two primary sections. First, we examined alterations in brain metrics throughout the course of pregnancy, where brain metrics (BrainMetric_ij_) were the dependent variable and gestational week (GesWeek_ij_) was the predictor. We also included age (Age_j_) and total intracranial volume (TIV_j_) as covariates for volumetric analyses (see Equation 1), and included Age_j_ and relative motion (Motion_ij_) as covariates for diffusion metric analyses^112^; see Equation 2.

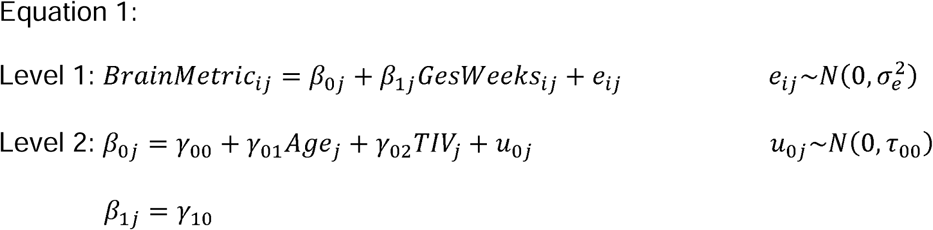

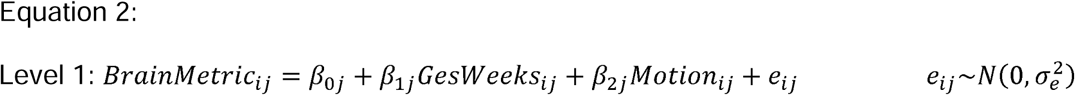

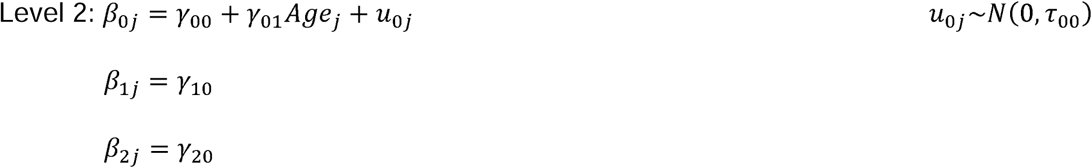

where y_10_ is the fixed effect of gestational weeks (i.e., slope), and y_00_ and u_0j_ are the fixed and random effects of the model intercept which allow each subject to have their own baseline level. To calculate the average percentage change in brain morphometry, we used the estimated LME models to predict the average change in brain morphometry over the range of observed gestational weeks (i.e., 12–39 weeks). Specifically, we used the estimated model to predict the brain metric at 12 and 39 weeks, respectively, and then computed the percentage change as:

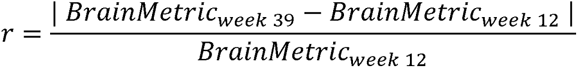

The second set of analyses focused on the potential mechanisms underlying the observed brain alterations by examining the associations between brain metrics and potential influencing factors, including hormone levels, CRP, and CSF volume. We first applied LME models to examine variations in hormone levels (progesterone, estradiol, testosterone, and cortisol) and CRP throughout pregnancy, with hormone levels and CRP as the dependent variable and gestational week as the fixed effect. CRP was log-transformed using a base 10 logarithm to normalize their distribution^113^. For brain metrics demonstrating significant alterations during pregnancy, we examined potential links with hormone levels and the total intracranial CSF volume using LME models in which BrainMetric_ij_ was the dependent variable and hormone levels (Hormone_ij_) or CSF volume was the predictor. Given the absence of significant changes in CRP, associational analyses between CRP levels and observed changes in brain metrics were not implemented. Moreover, we included Age_j_ and total TIV_j_ as covariates for volumetric analyses (see Equation 3); we included Age_j_ and Motion_ij_ as covariates for diffusion metric analyses (see Equation 4).

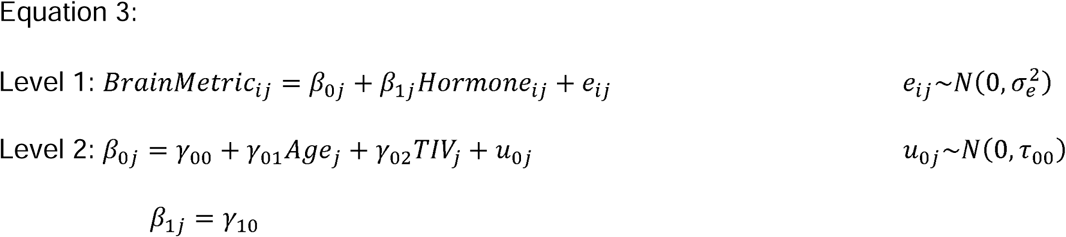

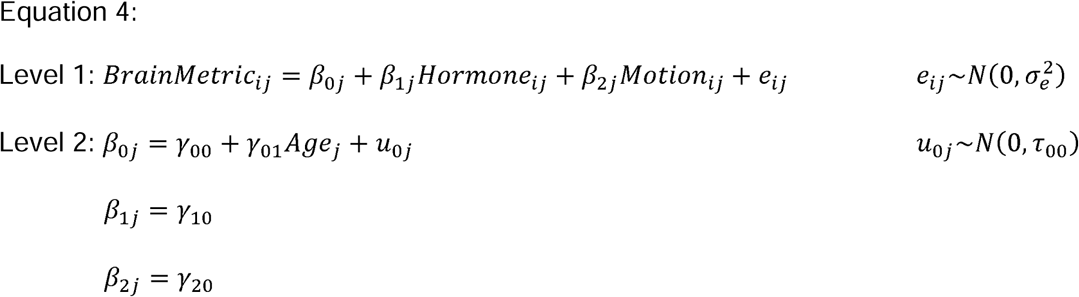

where *y*_10_ is the fixed effect of gestational weeks (i.e., slope), and *y*_00_ and *u*_0j_ are the fixed and random effects of the model intercept. We reported standard beta values and 95% confidence intervals (95% CI), with p-values tested based on a two-sided significance test.

Given the small sample size and the exploratory nature of our study, we did not correct for multiple comparisons^114^. Results with *p*-values < .05 were considered statistically significant. This decision was made to prioritize sensitivity over specificity, as it is particularly important in exploratory research to identify potential effects for further investigation in larger studies. Our results should therefore be interpreted with caution. We mitigated potential concerns by emphasizing our interpretation of the direction and magnitude of effect sizes. Additionally, recognizing the possibility that individual outliers could disproportionately influence the outcome in our small sample, we implemented leave-one-out analyses for our main findings, in which each participant was systematically removed and the analyses recomputed. Specifically, a custom R script was programmed to loop through the subject IDs, sequentially removing each participant by instructing the script to filter out the subject ID for each iteration. In each iteration, one subject was automatically excluded from the dataset and the LME models were estimated using the remaining subjects. This approach helps to validate that the observed effects are consistent and not artifacts of unique data points, thus enhancing the reliability of our findings^115^.

Additionally, we included all available data, including those with single time points, in our LME models to maximize the use of the data and increase the reliability of the findings^110^. To further validate our approach, we have also included results excluding the three participants with only a single time point in the supplementary materials results section. These results are consistent with those presented in the main text. We also considered the patterns of results across related dependent measures, as opposed to individual tests in isolation. The results of our study thus serve as a preliminary exploration into the complex domain of neurobiological dynamics during pregnancy.

## Data availability

Data supporting the findings and figures in this study are available at https://github.com/vanderbiltsealab/NCAP. All original data in this study are also available from the corresponding author upon reasonable request.

## Code availability

The code utilized in this study is publicly available at https://github.com/vanderbiltsealab/NCAP.

## ACKNOWLEDGMENTS

We extend our sincere gratitude to all participants for their contributions to this study. This research was supported by the Peabody Small Grant, Vanderbilt Brain Institute Trans-Institutional Program (TIPs) Grant, Vanderbilt Institute for Clinical and Translational Research Grant (VR61621) to KLH, and the Vanderbilt University Institute of Imaging Science Center for Human Imaging (1 S10OD021771 01). Author contributions were supported in part by T32MH18921 (HAP; LGB).

## Author Contributions

K.L.H., conceived and designed the study; H.A.P., S.R., W.B. collected data; Y.N., B.N.C., M.C.C., M.K.M. cleaned and processed the data; Y.N., D.A.C., L.G.B., M.C.C. performed data analysis; Y.N., M.C.C., K.L.H., B.N.C. interpreted results; Y.N. wrote the first draft of the paper; all authors (Y.N., B.N.C., M.C.C., S.R., H.A.P., L.G.B., W.B., M.K.M., D.A.C., E.W.C., S.S.O., S.A.S., A.K., K.L.H.) contributed to the manuscript final writing, editing, and reviewing.

## Supplementary methods

### Salivary hormone assessments

Saliva samples, collected immediately upon participants waking while participants remained in bed, were used for hormonal measurements. Participants were given instructions on the saliva collection process verbally, in writing, and through video demonstrations. They were also sent reminder texts the night before and the morning of the planned saliva collection. Participants were instructed to collect their saliva samples immediately upon waking, before consuming any food or drink that could potentially contaminate the samples. During collection, participants allowed saliva to pool on the floor of their mouths and then dribbled it into the provided collection vials, thus ensuring a consistent collection procedure. Participants recorded wake and collection times and stored samples in their home freezers. Participants were instructed to transport their samples frozen using ice baths/ice packs and an insulated bag to each session, where the samples were subsequently stored in a -20 degrees Fahrenheit laboratory freezer until they were shipped with dry ice to Salimetrics’ SalivaLab (Carlsbad, CA) for analysis. If participants failed to collect the sample immediately upon waking, they were instructed to defer sample collection until the following day to ensure adherence to the prescribed protocol. The hormones assessed included progesterone, estradiol, testosterone, and cortisol. Assays were conducted using the respective Salimetrics Salivary Assay Kits. Prior to the assays, samples were thawed to room temperature, vortexed, and centrifuged for 15 minutes at approximately 3,000 RPM (1,500 x g). The immunoassays were performed using sample test volumes of 50 μl for progesterone, 100 μl for estradiol, and 25 μl each for testosterone and cortisol. The average intra-assay and inter-assay coefficients of variation conformed to Salimetrics’ guidelines for accuracy and repeatability in Salivary Bioscience. To ensure synchronization between hormonal and MRI data, the samples were collected on the morning preceding and the morning of each MRI session. The measurements from these two time points were then averaged to be used in the analyses.

### Hair hormone assessments

Hair samples were collected from participants using hair scissors to cut a section of hair about 3 to 5 mm in thickness on the scalp from the posterior vertex position of the head. To reduce the frequency of sample collection, samples were requested at odd number assessments (e.g., first and third). Upon retrieving the sample, hair was secured in aluminum foil and stored in a dark, room temperature location until samples were ready to be assayed. From hair, hormone levels can be estimated at 1cm per month reliably for five months^1^. Thus, up to five values corresponding to the prior five months are estimated. Hair samples were analyzed using a commercially available immunoassay with chemiluminescence detection (CLIA, IBL-Hamburg, Germany). We used a column switching LC–APCI–MS/MS assay, a sensitive, reliable method for quantifying steroids in human hair^1–3^. Hair samples were washed with isopropanol and steroid hormones were extracted from non-pulverized hair by methanol incubation. The intra- and inter-assay coefficients of variation (CVs) for cortisol analysis by this method have been reported to range between 3.7% and 9.1%^4^.

We mapped hormone values to scan dates based on gestational age. For example, if a participant scanned three times four weeks apart and provided a sample at the third scan, but not the first or second, then the values derived from the segment of hair closest to the scalp were mapped onto the third scan, the values derived from the next segment were mapped onto the second scan, and the values derived from the third segment were mapped onto the first scan. Thus, even if participants did not provide hair samples at each scan, we were able to assign hormone values for prior scans. Outliers (*n* = 2), defined as having a z-score greater than 3 or lower than -3, were Winsorized to the next observed value. Following this, estimates were grand-mean centered and standardized to obtain standardized betas. Distance from the scalp was a fixed covariate. In analyzing the changes in hair hormone levels during pregnancy, participant age and distance from the scalp were incorporated as fixed effect covariates. For the associations between hair hormone levels and brain metrics, participant age, distance from the scalp, and total intracranial volume were included as covariates in volumetric analyses.

Participant age, distance from the scalp, and relative motion were included as covariates in the analysis of diffusion metrics.

### Inflammatory cytokines assessments

Blood samples for inflammatory cytokine assessments were collected using dried blood spot (DBS) sampling. The DBS method provides a relatively non-invasive alternative to venipuncture. At each visit, up to 5 drops of capillary whole blood were collected on filter paper (Whatman 903) following a finger stick with a sterile microlancet. The filter paper is certified to meet performance standards for sample absorption and lot-to-lot consistency by the Food and Drug Administration regulations for Class II Medical Devices. After collection, samples were dried at room temperature for approximately 4 hours. Once dried, samples were stored at -20 degrees Celsius prior to shipment. Samples were express shipped to the Northwestern University’s Laboratory for Human Biology Research for analysis. Samples were analyzed for interleukin-6 (IL-6), interleukin-10 (IL-10), tumor necrosis factor alpha (TNF□), and C-reactive protein (CRP) using highly sensitive immunoassay protocols validated for use with DBS samples^5,6^. Briefly, 2 x 5 mm punches of the DBS sample, calibration material, and controls were eluted in 100 uL buffer overnight in a filter plate (Millipore MultiScreen MSHVN4510) and transferred to assay plates for IL-6 (R&D Systems, HS600B), IL-10, and TNF□. Assay protocols were completed following the manufacturer’s instructions. To minimize between-assay variation, all samples were analyzed using a single lot of reagents, and samples were distributed randomly across assay plates. CRP was quantified in DBS samples using a high sensitivity enzyme immunoassay protocol previously developed for use with DBS^7^. The Laboratory conducted all assay analyses on the DBS samples. However, the usability of the samples for IL-6, IL-10, and TNF□ was limited, resulting in a low yield of successfully assessed data for these cytokines (less than 50%). Thus, only CRP was utilized in the analyses. CRP was log-transformed using a base 10 logarithm to normalize its distribution^8^. Detailed descriptive statistics are provided in Table S1.

### MRI acquisitions

MRI data acquisition was conducted on a 3T Philips Achieva scanner using a 32-channel head coil at Vanderbilt University Institute of Imaging Science. Participants were provided with headphones to reduce the noise generated by the magnet. Foam paddings were placed around the midsection to attenuate scanner sounds. Participants were instructed to remain as still as possible throughout the scan to minimize potential motion artifacts.

At Philips, all MRI scanners undergo four preventative maintenance visits per year. During these visits, various quality and safety checks are conducted, some at each visit and others on a rotating schedule, including: (1) earth bonding and grounding checks, (2) torque checks and safety inspections of main, sub, and gradient connections, (3) liquid cooling system checks, including flow regulation, system charge level, system recharge and flush and fill, flow interlock safety check, leak inspections, and strainer and filtration cleaning, along with amplifier flow checks, (4) cryocooler/coldhead checks to ensure consistent and reliable performance, (5) coil inspections and connection cleaning, and (6) body coil transmission characteristics. During each preventative maintenance visit, a Philips engineer performs a periodic image quality test, which assesses system performance across specifications, including signal-to-noise ratio, geometry and scale, and uniformity. Test results are compared against a set of specification files maintained by the Business Unit for each system model, which the engineer verifies and updates at every preventative maintenance visit. Each coil on the system also undergoes an individual image quality test as necessary. Moreover, all contract MRI scanners, including the scanner used in this study, are remotely monitored by Philips, with daily reports on various performance metrics covering parameters such as cryogenic system performance, noise levels, coil functionality, hardware faults, cooling system performance, and site temperature and humidity. Any metrics indicating out-of-specification performance trigger a review by remote engineers or local field engineers. This remote monitoring enables Philips engineers to proactively identify and address potential issues before they impact users, ensuring continuous system performance at the subassembly level.

### Anatomical MRI

All images included in the final analyses had an Euler number of 2, with no holes detected, indicating high-quality surface reconstruction. Specifically, the Euler number describes the topographical complexity of the reconstructed cortical surface. The formula for the Euler number is, where is the number of holes in the surface. Therefore, the Euler number for a flat and smooth surface with no holes is 2. A good surface reconstruction algorithm aims to minimize the number of holes in the surface. This means the closer the surface’s Euler number comes to 2, the better the reconstruction, which implies better data quality^9^. Additionally, to provide a more thorough assessment of data quality, we have included raw T1-weighted images, FreeSurfer-processed data, and aseg files for five randomly selected subjects as shown below.

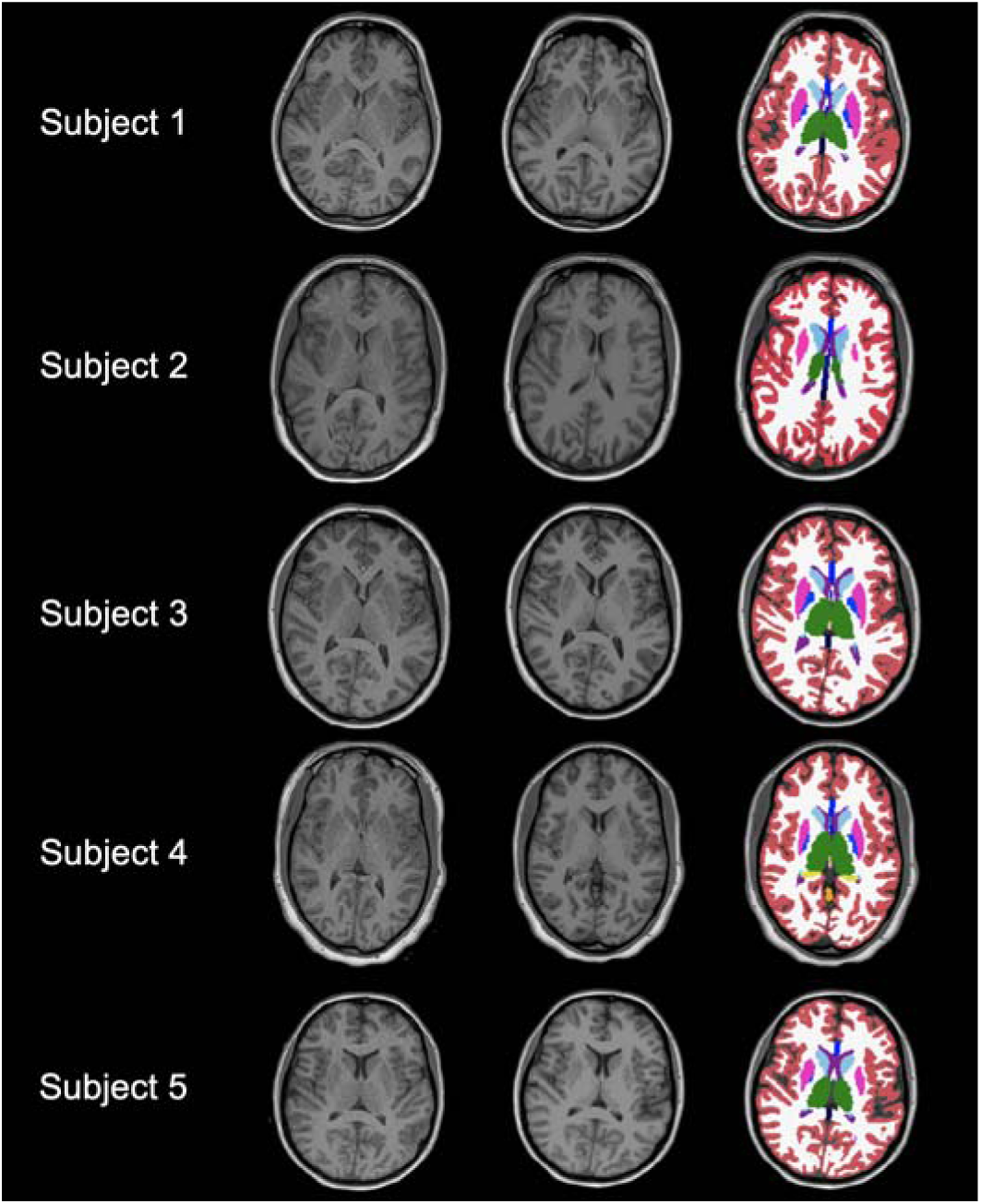

### Diffusion-weighted MRI

The raw diffusion data were processed using MRtrix3^10^ and FSL (FMRIB’s Software Library^11^). Our initial step involved noise estimation and removal via the dwidenoise function from MRtrix3^12,13^. The eddy and susceptibility distortion corrections were performed using the dwifslpreproc function in MRtrix3, which implemented the TOPUP and EDDY tools from FSL to correct for motion and eddy current artifacts, including outlier replacement^11,14–16^.

Subsequent to preprocessing, we computed the neurite density index (NDI) and orientation dispersion indices (ODI) using the NODDI Matlab Toolbox^17^. Although our scanning protocol was primarily configured for NODDI modeling, we supplemented our processing and results with diffusion tensor imaging (DTI) modeling to elucidate a wider spectrum of changes that may occur during pregnancy and also facilitated comparability with other studies that utilize DTI in diffusion research. From DTI, we derived fractional anisotropy (FA), mean diffusivity (MD), axial diffusivity (AD), and radial diffusivity (RD) in MRtrix3 with an iteratively reweighted linear tensor fitting procedure^18,19^.

We then used the Computational Anatomy Toolbox (CAT12^20^) for SPM12 (www.fil.ac.uk/spm/) in Matlab R2021a (The Mathworks, Inc., Natick, MA, USA) for quality assessment of the scalar NODDI and DTI images, allowing us to eliminate potential outliers prior to the statistical analysis. A sample homogeneity check was carried out for each scalar NODDI and DTI image. Any scan classified as an outlier in any scalar image was excluded from all subsequent metrics analyses. The quality assessment revealed significant discrepancies between the overall sample and three scans from three participants (3 outliers), leading to their removal, resulting in 24/27 scans for the final diffusion analysis.

In order to analyze changes across tracts, we first estimated fiber orientation distributions (FOD) using MRtrix3’s dwi2fod msmt_csd and then normalized them using MRtrix3’s mtnormalise^21,22^. The normalized FODs were then fed to TractSeg to automatically segment fiber tracts, resulting in 72 unique fiber bundles^23^. We sampled each pre-generated NODDI and DTI image along each tract using TractSeg’s Tractometry^23,24^. The outcome included NODDI and DTI metrics for each tract across 50 established tracts, which were subsequently used in our statistical analyses.

Lastly, to characterize the global changes for each NODDI metric and investigate if observed tract-level changes reflected a general alteration across all segmented tracts, we adopted two distinct approaches. The first approach was to compute a single averaged value across all segmented tracts. Recognizing the limitations of this method, particularly the neglect of tract size variations, we implemented a supplementary approach. This involved conducting whole-brain streamline tractography using MRtrix3. Subsequently, we transformed the complete set of derived streamlines into a whole-brain density image utilizing MRtrix3’s tckmap^25^. SIFT2 weights were incorporated into this calculation to down-weight the contributions of spurious fibers to the voxel-level average estimate^26^. The density images were multiplied by the subject-specific proportionality coefficient to allow for valid group-level comparisons. The resulting track-weighted image was used to weigh each metric. Each weighted metric imaging was then averaged within individual white matter masks. This approach enabled us to derive a single value for each NODDI and DTI metric, thereby summarizing the global changes in these metrics during pregnancy.

### Whole-brain Streamline Tractography

We used a custom, publicly-available pipeline (https://github.com/conradbn/DWI_Tractography) that was written in Python and built from the MRtrix3 software package^10^. To begin the tractography process, the preprocessed DWI data were passed to an unsupervised method for estimating single-fiber response functions for white matter (WM), gray matter (GM), and cerebrospinal fluid (CSF)^27^. Multi-shell, multi-tissue constrained spherical deconvolution was performed to obtain WM fiber orientation distributions (FODs) as well as for the GM and CSF compartments in all voxels^21^. The FOD maps were then corrected for intensity inhomogeneities and bias fields^28^. Probabilistic tracking was employed to construct a whole-brain tractogram via the tckgen command and the “iFOD2” algorithm^29^. We generated 10 million streamlines which were randomly seeded and terminated at the GM/WM interface as delineated by the FreeSurfer Longitudinal pipeline segmentation files, in a method known as anatomically-constrained tractography (ACT^30^). ACT helps improve the biological validity of streamlines by ensuring that a streamline begins and ends (i.e., “synapses”) at some GM structure. Finally, we used the “spherical-deconvolution informed filtering of tractograms” method (SIFT2), to correct for known biases in tractography by assigning a cross-sectional area weight to each streamline^26^. SIFT2 provides a weighted streamline reconstruction that optimally fits the underlying CSD model at each voxel, effectively down-weighting false-positive streamlines.

## Supplementary results

### White matter microstructure changes

DTI metrics (FA, MD, AD, and RD) were analyzed using LME models. A heatmap was produced (see Fig. S4) illustrating the standardized regression coefficients for all segmented tracts. Across the tracts, we observed a predominant increasing trend in FA, consistent with the observed NDI enhancements. Importantly, tracts exhibiting the most pronounced FA increases largely overlapped with those showing significant NDI increases (see Fig. S5). Specifically, these were projection tracts implicated in sensorimotor functions, such as the corticospinal tract (CST), fronto-pontine tract (FPT), parieto-occipital pontine (PORT), superior cerebellar peduncle (SCP), and thalamo-premotor tract (T_PREM). Additionally, MD and RD metrics demonstrated decreasing trends, particularly in association and projection tracts integral to motor and sensory functions. Conversely, AD metrics remained relatively unchanged, with the exception of the SCP. Detailed statistical results for these DTI metrics (FA, MD, AD, and RD) are available in Supplementary Tables S8-S11.

Lastly, we evaluated whether the tract-level changes that were observed suggested any broad alterations across all the segmented tracts. For this, we used average measurements derived from all segmented tracts. Our analysis revealed that global metrics of NDI (▱ = 0.235, 95% CI [-0.242, 0.713], *p* = .301) and ODI (▱ = -0.089, 95% CI [-0.496, 0.317], *p* = .638) did not demonstrate significant changes over the course of pregnancy. Similar patterns were observed for global DTI metrics, none of which exhibited significant changes during pregnancy, including FA (▱ = 0.310, 95% CI [-0.077, 0.696], *p* = .106), MD (▱ = -0.016, 95% CI [-0.344, 0.312], *p* = .917), AD (▱ = 0.316, 95% CI [-0.128, 0.760], *p* = .146), and RD (▱ = -0.157, 95% CI [-0.479, 0.165], *p* = .306). Moreover, we implemented an alternative method to characterize global measures to examine whether changes in diffusion metrics were global. The results indicated that global track-weighted metrics did not significantly fluctuate over pregnancy, including NDI (▱ = 0.089, 95% CI [-0.325, 0.502], *p* = .646), ODI (▱ = -0.008, 95% CI [-0.235, 0.219], *p* = .940), FA (▱ = 0.022, 95% CI [-0.317, 0.360], *p* = .890), MD (▱ = -0.101, 95% CI [-0.399, 0.196], *p* = .470), AD (▱ = -0.074, 95% CI [-0.411, 0.263], *p* = .639), and RD (▱ = -0.125, 95% CI [-0.384, 0.133], *p* = .310).

### Hair hormones changes

Covarying for age and distance from the scalp, we observed statistically significant increases in hair hormonal levels of cortisol (▱ = 0.730, 95% CI [0.441, 1.019], *p* < .001) and cortisone (▱ = 0.706, 95% CI [0.283, 1.130], *p* = .003) across pregnancy. The increase in progesterone levels was moderate and not statistically significant (▱ = 0.141, 95% CI [-0.333, 0.615], *p* = .536). While sex hormones are known to rise over the gestational period, our hair hormone assessments did not show a statistically significant increase in progesterone levels. Our data did not demonstrate the expected increase, suggesting potential limitations in the method used for measuring progesterone in hair samples or possible issues with sample collection. Previous studies have successfully measured progesterone changes in hair samples during pregnancy, indicating the feasibility of assessing this hormone using hair assays^31,32^.

However, these studies also note variability and potential limitations in using hair samples for hormone measurement. We have included this information here to provide a complete dataset for future examination and reference.

### Associations between hair hormones and brain structure

Having observed significant increases in hair cortisol and cortisone levels, we then explored their associations with brain volumetric measurements. After covarying for TIV, age, and distance from the scalp, no statistically significant associations were identified between hair cortisol and total brain volume (▱ = -0.072, 95% CI [-0.170, 0.027], *p* = .138), total gray matter volume (▱ = -0.103, 95% CI [-0.252, 0.046], *p* = .159), total cortical volume (▱ = -0.135, 95% CI [-0.319, 0.049], *p* = .137), and mean cortical thickness (▱ = -0.208, 95% CI [-0.752, 0.335], *p* = .423). Similarly, there were no significant relations between hair cortisone and total brain volume (▱ = -0.062, 95% CI [-0.136, 0.011], *p* = .090), total gray matter volume (▱ = -0.096, 95% CI [-0.205, 0.013], *p* = .078), total cortical volume (▱ = -0.103, 95% CI [-0.243, 0.037], *p* = .136), and mean cortical thickness (▱ = -0.155, 95% CI [-0.573, 0.262], *p* = .436).

### Associations between hair hormones and diffusion metrics

NDI demonstrated moderate positive links with both hair cortisol and cortisone levels. Detailed statistical outcomes of analyses are available in Table S21-S22.

### Brain morphometry changes and associations between salivary hormones and brain morphometry after excluding participants with single time point data

After excluding three participants with only a single time point, the results were consistent with those from the full sample. Specifically, total brain volume (▱ = -0.098, 95% CI [-0.159, -0.037], *p* = .004), total gray matter volume (▱ = -0.188, 95% CI [-0.295, -0.082], *p* = .002), total cortical volume (▱ = -0.230, 95% CI [-0.364, -0.095], *p* = .003), subcortical volume (▱ = -0.099, 95% CI [-0.199, 0.001], *p* = .053), and cortical thickness (▱ = -0.520, 95% CI [-0.923, -0.117], *p* = .015) exhibited decreases across pregnancy. Cortical surface area (▱ = -0.014, 95% CI [-0.047, 0.019], *p* = .375) and total white matter volume (▱ = -0.026, 95% CI [-0.108, 0.056], *p* = .505) did not exhibit a statistically significant change across pregnancy. Moreover, there was a statistically significant increase in total intracranial CSF (▱ = 0.187, 95% CI [0.001, 0.373], *p* = .049).

Additionally, there were statistically significant negative associations between salivary levels of progesterone and total brain volume (▱ = -0.101, 95% CI [-0.166, -0.035], *p* = .006), total gray matter volume (▱ = -0.189, 95% CI [-0.306, -0.073], *p* = .004), total cortical volume (▱ = -0.247, 95% CI [-0.386, -0.108], *p* = .002) and mean cortical thickness (▱ = -0.661, 95% CI [-1.030, -0.291], *p* = .002).

### Interaction effect of gestational age by the number of live births on brain morphometry changes

We explored the interaction effect of gestational age by the number of live births and found a significant interaction effect on total brain volume (▱ = 0.063, 95% CI [0.005, 0.122], *p* = .036), total gray matter volume (▱ = 0.134, 95% CI [0.048, 0.221], *p* = .005), total cortical volume (▱ = 0.169, 95% CI [0.061, 0.277], *p* = .005), and cortical thickness 0 = 0.373, 95% CI [0.032, 0.715], *p* = .034), but not on subcortical volume (▱ = 0.010, 95% CI [-0.100, 0.119], *p*= .851), cortical surface area (▱ = 0.016, 95% CI [-0.019, 0.051], *p* = .346) and total white matter volume (▱ = -0.005, 95% CI [-0.098, 0.088], *p* = .911). These findings suggest that individuals with a greater number of births experience less reduction in total brain volume, total gray matter volume, total cortical volume, and cortical thickness. However, it is important to note that our already small sample size makes the model underpowered when incorporating the interaction term. Therefore, caution is needed when interpreting these findings.

### Brain morphometry changes covarying for the number of live births

To further validate our results, we further included the number of live births as a covariate in addition to the original covariates. The results were consistent with those from the full sample. Specifically, total brain volume (▱ = -0.105, 95% CI [-0.169, -0.040], *p* = .004), total gray matter volume (▱ = -0.192, 95% CI [-0.302, -0.082], *p* = .002), total cortical volume (▱ = -0.238, 95% CI [-0.375, -0.101], *p* = .002), subcortical volume (▱ = -0.095, 95% CI [-0.197, 0.008], *p* = .067), and cortical thickness (▱ = -0.520, 95% CI [-0.921, -0.119], *p* = .015) exhibited decreases across pregnancy. Cortical surface area (▱ = -0.017, 95% CI [-0.052, 0.018], *p* = .313) and total white matter volume (▱ = -0.030, 95% CI [-0.119, 0.058], *p* = .473) did not exhibit a statistically significant change. Moreover, there was a statistically significant increase in total intracranial CSF (▱ = 0.243, 95% CI [0.083, 0.403], *p* = .006).

Additionally, there were statistically significant negative associations between salivary levels of progesterone and total brain volume (▱ = -0.102, 95% CI [-0.169, -0.034], *p* = .006), total gray matter volume (▱ = -0.190, 95% CI [-0.307, -0.073], *p* = .004), total cortical volume (▱ = -0.253, 95% CI [-0.390, -0.116], *p* = .002) and mean cortical thickness (▱ = -0.685, 95% CI [-1.055, -0.314], *p* = .002).

---

**Table S1.** Descriptive statistics for inflammatory cytokines. IL-6: interleukin-6, IL-10: interleukin-10, TNF□: tumor necrosis factor alpha, CRP: C-reactive protein.

| Inflammatory Marker | ***N***<br>Obtained<br>Samples | ***N***<br>Usable<br>Samples | ***M*** | ***SD*** | Range |
| --- | --- | --- | --- | --- | --- |
| IL-6 (pg/mL) | 27 | 13 | 0.34 | 0.36 | 0.09–1.50 |
| IL-10 (pg/mL) | 27 | 13 | 0.28 | 0.23 | 0.01–0.86 |
| TNFα (pg/mL) | 27 | 13 | 2.43 | 0.78 | 1.53–4.04 |
| CRP (mg/L) | 27 | 26 | 2.08 | 1.30 | 0.19–4.25 |

**Fig. S1.**
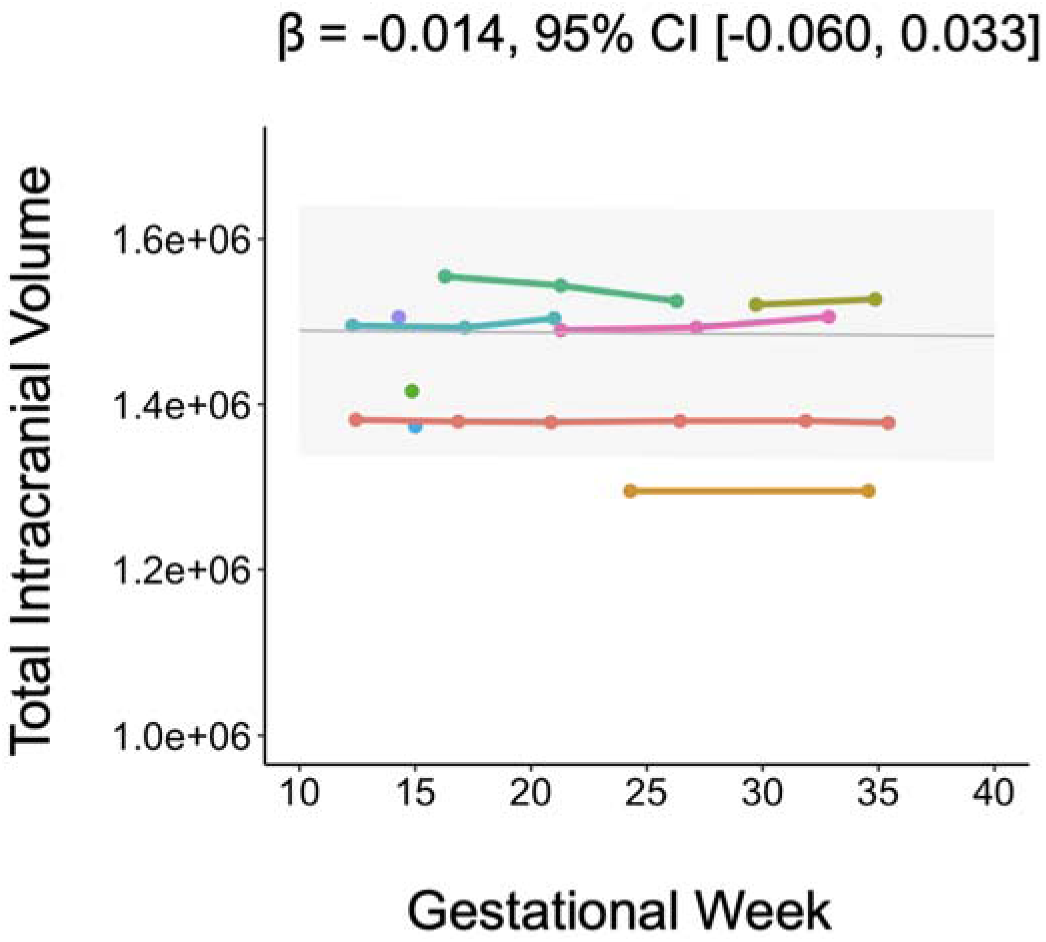
Changes in total intracranial volume throughout pregnancy. Total intracranial volume did not exhibit a meaningful or statistically significant change over the course of pregnancy (▱ = -0.014, 95% CI [-0.060, 0.033], *p* = .539).

**Fig. S2.**
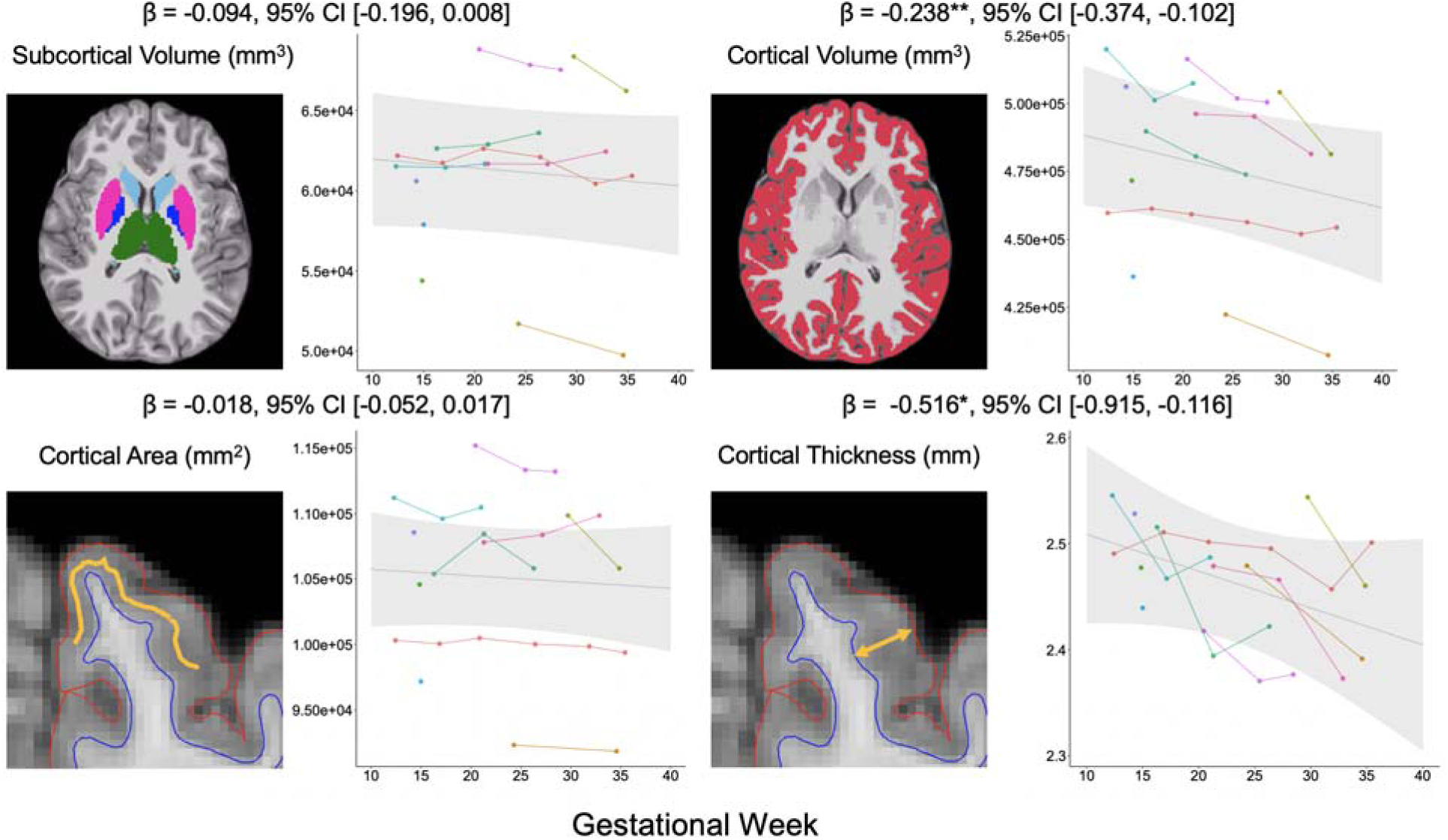
Changes in subcortical volume, cortical volume, cortical area, and cortical thickness during pregnancy. Total cortical volume significantly declined over the course of pregnancy (▱ = -0.238, 95% CI [-0.374, -0.102], *p* = .002), whereas total subcortical volume showed a non-significant decreasing trend (▱ = -0.094, 95% CI [-0.196, 0.008], *p* = .068). This translates to a 4.9% decline over the observed gestational age range in this study (i.e., 12–39 weeks). Subsequently, we assessed changes in mean cortical surface area and cortical thickness respectively. We found that cortical thickness decreased during pregnancy (▱ = -0.516, 95% CI [-0.915, -0.116], *p* = .015), and cortical surface area remained unchanged (▱ = -0.018, 95% CI [-0.052, 0.017], *p* = .297). This translates to a 3.7% decline over the observed gestational age range in this study. **Note**. * *p* < .05. ** *p* < .01.

**Table S2.**
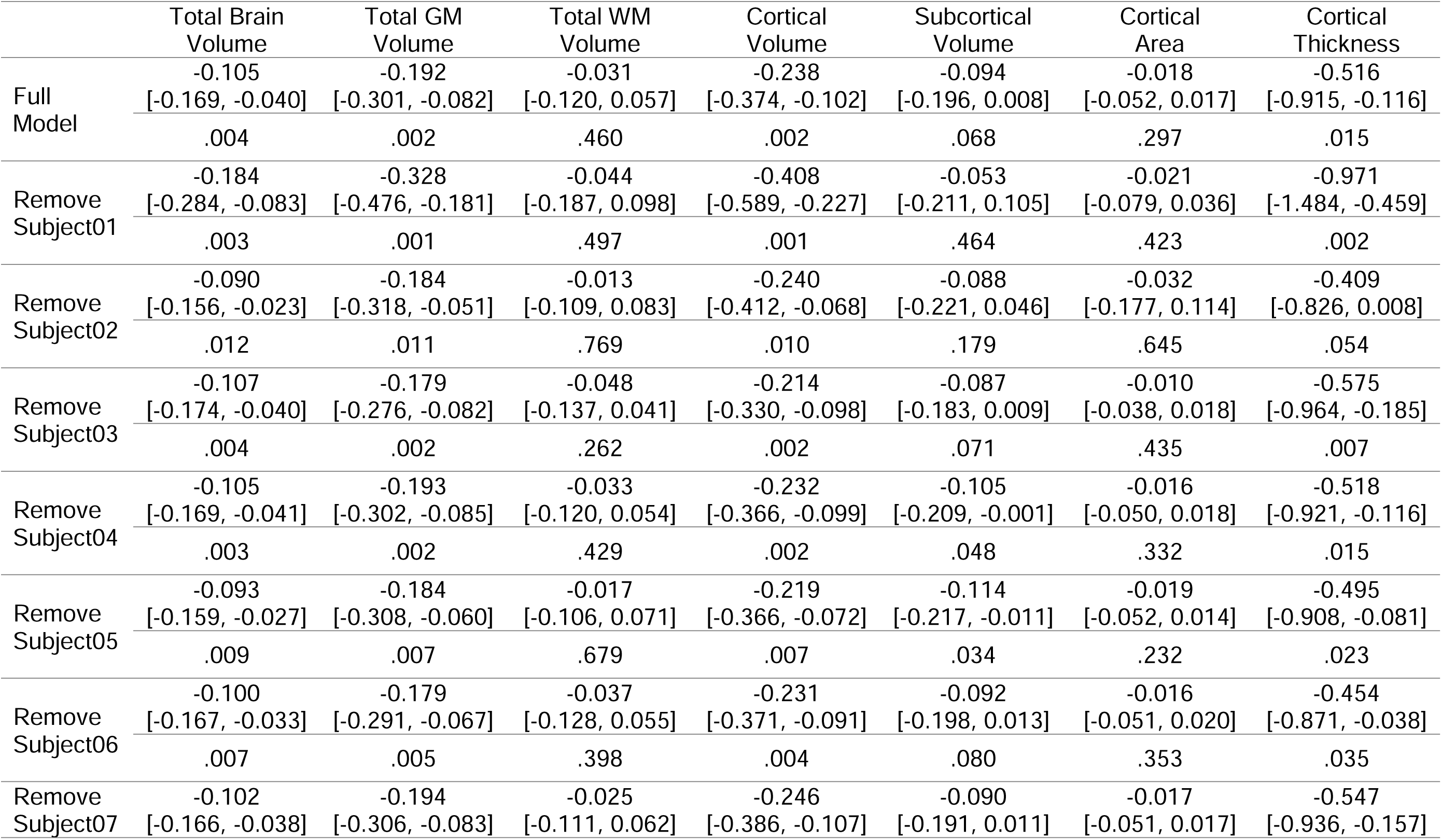
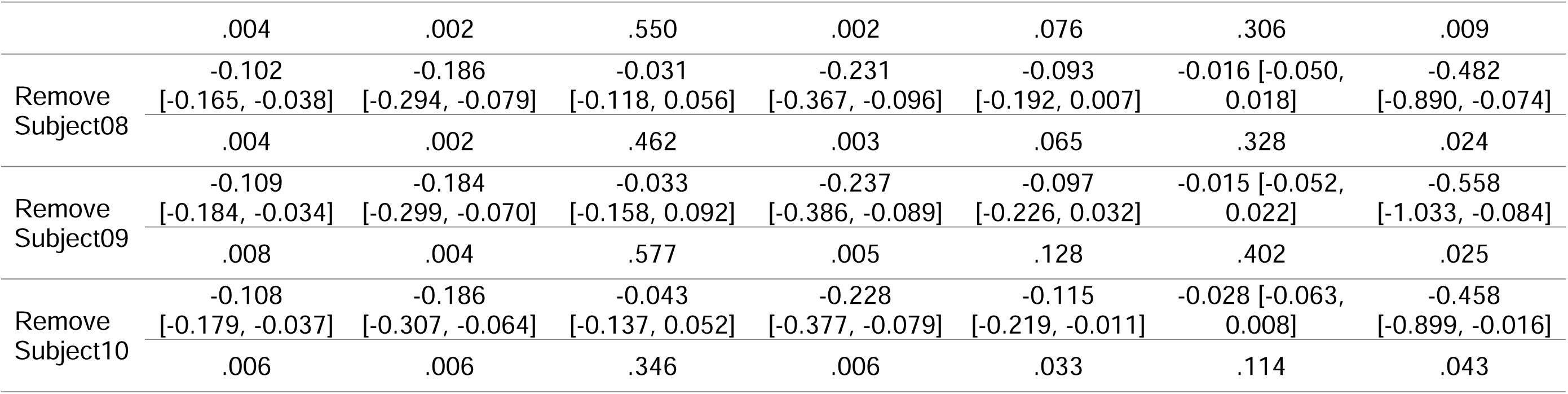
Statistical outcomes for leave-one-out cross-validation analyses of brain volumes. The columns specified the type of analysis conducted with the leave-one-out cross-validation (i.e., total brain volume, total gray matter volume, total white matter volume, cortical volume, subcortical volume, cortical area, and cortical thickness). Each row listed the participant who was excluded from the analysis. Within each cell, the standardized regression coefficients were provided at the top, followed by the 95% confidence intervals in parentheses, and the *p*-values were listed at the bottom. GM: gray matter, WM: white matter.

|  | Total Brain<br>Volume | Total GM<br>Volume | Total WM<br>Volume | Cortical<br>Volume | Subcortical<br>Volume | Cortical<br>Area | Cortical<br>Thickness |
| --- | --- | --- | --- | --- | --- | --- | --- |
| Full<br>Model | -0.105<br>[-0.169, -0.040] | -0.192<br>[-0.301, -0.082] | -0.031<br>[-0.120, 0.057] | -0.238<br>[-0.374, -0.102] | -0.094<br>[-0.196, 0.008] | -0.018<br>[-0.052, 0.017] | -0.516<br>[-0.915, -0.116] |
|  | .004 | .002 | .460 | .002 | .068 | .297 | .015 |
| Remove<br>Subject01 | -0.184<br>[-0.284, -0.083] | -0.328<br>[-0.476, -0.181] | -0.044<br>[-0.187, 0.098] | -0.408<br>[-0.589, -0.227] | -0.053<br>[-0.211, 0.105] | -0.021<br>[-0.079, 0.036] | -0.971<br>[-1.484, -0.459] |
|  | .003 | .001 | .497 | .001 | .464 | .423 | .002 |
| Remove<br>Subject02 | -0.090<br>[-0.156, -0.023] | -0.184<br>[-0.318, -0.051] | -0.013<br>[-0.109, 0.083] | -0.240<br>[-0.412, -0.068] | -0.088<br>[-0.221, 0.046] | -0.032<br>[-0.177, 0.114] | -0.409<br>[-0.826, 0.008] |
|  | .012 | .011 | .769 | .010 | .179 | .645 | .054 |
| Remove<br>Subject03 | -0.107<br>[-0.174, -0.040] | -0.179<br>[-0.276, -0.082] | -0.048<br>[-0.137, 0.041] | -0.214<br>[-0.330, -0.098] | -0.087<br>[-0.183, 0.009] | -0.010<br>[-0.038, 0.018] | -0.575<br>[-0.964, -0.185] |
|  | .004 | .002 | .262 | .002 | .071 | .435 | .007 |
| Remove<br>Subject04 | -0.105<br>[-0.169, -0.041] | -0.193<br>[-0.302, -0.085] | -0.033<br>[-0.120, 0.054] | -0.232<br>[-0.366, -0.099] | -0.105<br>[-0.209, -0.001] | -0.016<br>[-0.050, 0.018] | -0.518<br>[-0.921, -0.116] |
|  | .003 | .002 | .429 | .002 | .048 | .332 | .015 |
| Remove<br>Subject05 | -0.093<br>[-0.159, -0.027] | -0.184<br>[-0.308, -0.060] | -0.017<br>[-0.106, 0.071] | -0.219<br>[-0.366, -0.072] | -0.114<br>[-0.217, -0.011] | -0.019<br>[-0.052, 0.014] | -0.495<br>[-0.908, -0.081] |
|  | .009 | .007 | .679 | .007 | .034 | .232 | .023 |
| Remove<br>Subject06 | -0.100<br>[-0.167, -0.033] | -0.179<br>[-0.291, -0.067] | -0.037<br>[-0.128, 0.055] | -0.231<br>[-0.371, -0.091] | -0.092<br>[-0.198, 0.013] | -0.016<br>[-0.051, 0.020] | -0.454<br>[-0.871, -0.038] |
|  | .007 | .005 | .398 | .004 | .080 | .353 | .035 |
| Remove<br>Subject07 | -0.102<br>[-0.166, -0.038] | -0.194<br>[-0.306, -0.083] | -0.025<br>[-0.111, 0.062] | -0.246<br>[-0.386, -0.107] | -0.090<br>[-0.191, 0.011] | -0.017<br>[-0.051, 0.017] | -0.547<br>[-0.936, -0.157] |
|  | .004 | .002 | .550 | .002 | .076 | .306 | .009 |
| Remove Subject08 | -0.102<br>[-0.165, -0.038] | -0.186<br>[-0.294, -0.079] | -0.031<br>[-0.118, 0.056] | -0.231<br>[-0.367, -0.096] | -0.093<br>[-0.192, 0.007] | -0.016 [-0.050, 0.018] | -0.482<br>[-0.890, -0.074] |
|  | .004 | .002 | .462 | .003 | .065 | .328 | .024 |
| Remove Subject09 | -0.109<br>[-0.184, -0.034] | -0.184<br>[-0.299, -0.070] | -0.033<br>[-0.158, 0.092] | -0.237<br>[-0.386, -0.089] | -0.097<br>[-0.226, 0.032] | -0.015 [-0.052, 0.022] | -0.558<br>[-1.033, -0.084] |
|  | .008 | .004 | .577 | .005 | .128 | .402 | .025 |
| Remove Subject10 | -0.108<br>[-0.179, -0.037] | -0.186<br>[-0.307, -0.064] | -0.043<br>[-0.137, 0.052] | -0.228<br>[-0.377, -0.079] | -0.115<br>[-0.219, -0.011] | -0.028 [-0.063, 0.008] | -0.458<br>[-0.899, -0.016] |
|  | .006 | .006 | .346 | .006 | .033 | .114 | .043 |

**Table S3.**
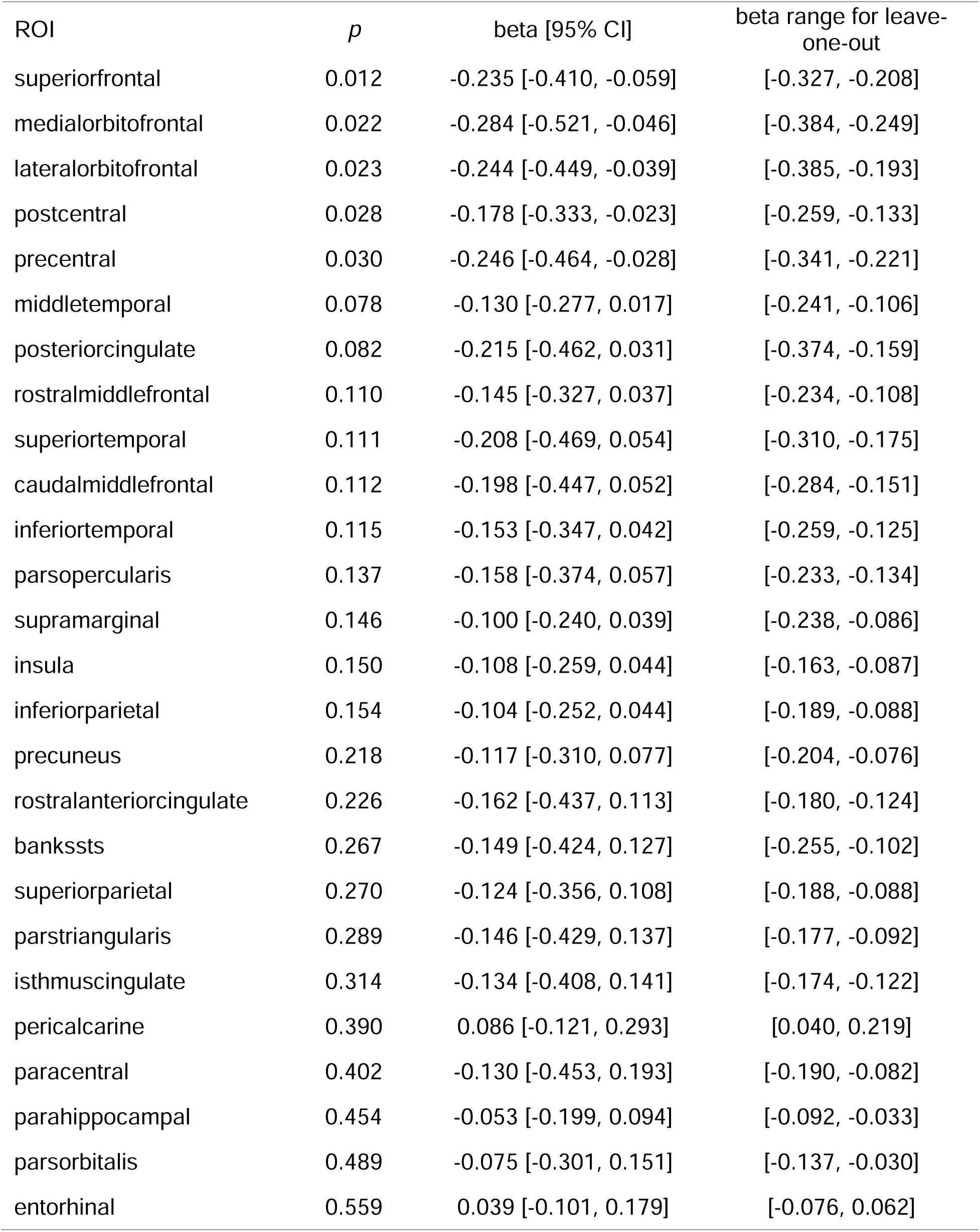
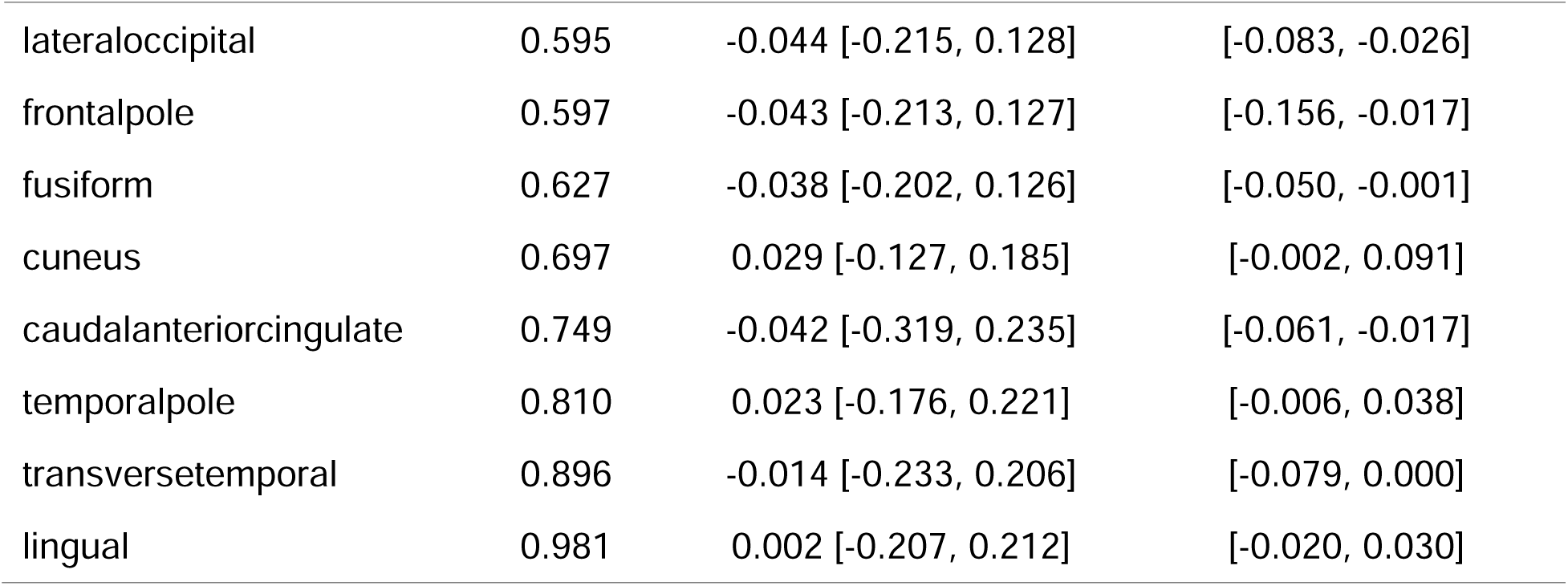
Statistical outcomes of linear mixed effects models that were implemented for the analysis of cortical region volumes. These regions, as defined by the Desikan-Killiany atlas, were assessed against gestational week to evaluate changes during pregnancy. ROI: region of interest.

| ROI | *p* | beta [95% CI] | beta range for leave-one-out |
| --- | --- | --- | --- |
| superiorfrontal | 0.012 | -0.235 [-0.410, -0.059] | [-0.327, -0.208] |
| medialorbitofrontal | 0.022 | -0.284 [-0.521, -0.046] | [-0.384, -0.249] |
| lateralorbitofrontal | 0.023 | -0.244 [-0.449, -0.039] | [-0.385, -0.193] |
| postcentral | 0.028 | -0.178 [-0.333, -0.023] | [-0.259, -0.133] |
| precentral | 0.030 | -0.246 [-0.464, -0.028] | [-0.341, -0.221] |
| middletemporal | 0.078 | -0.130 [-0.277, 0.017] | [-0.241, -0.106] |
| posteriorcingulate | 0.082 | -0.215 [-0.462, 0.031] | [-0.374, -0.159] |
| rostralmiddlefrontal | 0.110 | -0.145 [-0.327, 0.037] | [-0.234, -0.108] |
| superiortemporal | 0.111 | -0.208 [-0.469, 0.054] | [-0.310, -0.175] |
| caudalmiddlefrontal | 0.112 | -0.198 [-0.447, 0.052] | [-0.284, -0.151] |
| inferiortemporal | 0.115 | -0.153 [-0.347, 0.042] | [-0.259, -0.125] |
| parsopercularis | 0.137 | -0.158 [-0.374, 0.057] | [-0.233, -0.134] |
| supramarginal | 0.146 | -0.100 [-0.240, 0.039] | [-0.238, -0.086] |
| insula | 0.150 | -0.108 [-0.259, 0.044] | [-0.163, -0.087] |
| inferiorparietal | 0.154 | -0.104 [-0.252, 0.044] | [-0.189, -0.088] |
| precuneus | 0.218 | -0.117 [-0.310, 0.077] | [-0.204, -0.076] |
| rostralanteriorcingulate | 0.226 | -0.162 [-0.437, 0.113] | [-0.180, -0.124] |
| bankssts | 0.267 | -0.149 [-0.424, 0.127] | [-0.255, -0.102] |
| superiorparietal | 0.270 | -0.124 [-0.356, 0.108] | [-0.188, -0.088] |
| parstriangularis | 0.289 | -0.146 [-0.429, 0.137] | [-0.177, -0.092] |
| isthmuscingulate | 0.314 | -0.134 [-0.408, 0.141] | [-0.174, -0.122] |
| pericalcarine | 0.390 | 0.086 [-0.121, 0.293] | [0.040, 0.219] |
| paracentral | 0.402 | -0.130 [-0.453, 0.193] | [-0.190, -0.082] |
| parahippocampal | 0.454 | -0.053 [-0.199, 0.094] | [-0.092, -0.033] |
| parsorbitalis | 0.489 | -0.075 [-0.301, 0.151] | [-0.137, -0.030] |
| entorhinal | 0.559 | 0.039 [-0.101, 0.179] | [-0.076, 0.062] |
| lateraloccipital | 0.595 | -0.044 [-0.215, 0.128] | [-0.083, -0.026] |
| frontalpole | 0.597 | -0.043 [-0.213, 0.127] | [-0.156, -0.017] |
| fusiform | 0.627 | -0.038 [-0.202, 0.126] | [-0.050, -0.001] |
| cuneus | 0.697 | 0.029 [-0.127, 0.185] | [-0.002, 0.091] |
| caudalanteriorcingulate | 0.749 | -0.042 [-0.319, 0.235] | [-0.061, -0.017] |
| temporalpole | 0.810 | 0.023 [-0.176, 0.221] | [-0.006, 0.038] |
| transversetemporal | 0.896 | -0.014 [-0.233, 0.206] | [-0.079, 0.000] |
| lingual | 0.981 | 0.002 [-0.207, 0.212] | [-0.020, 0.030] |

**Table S4.**
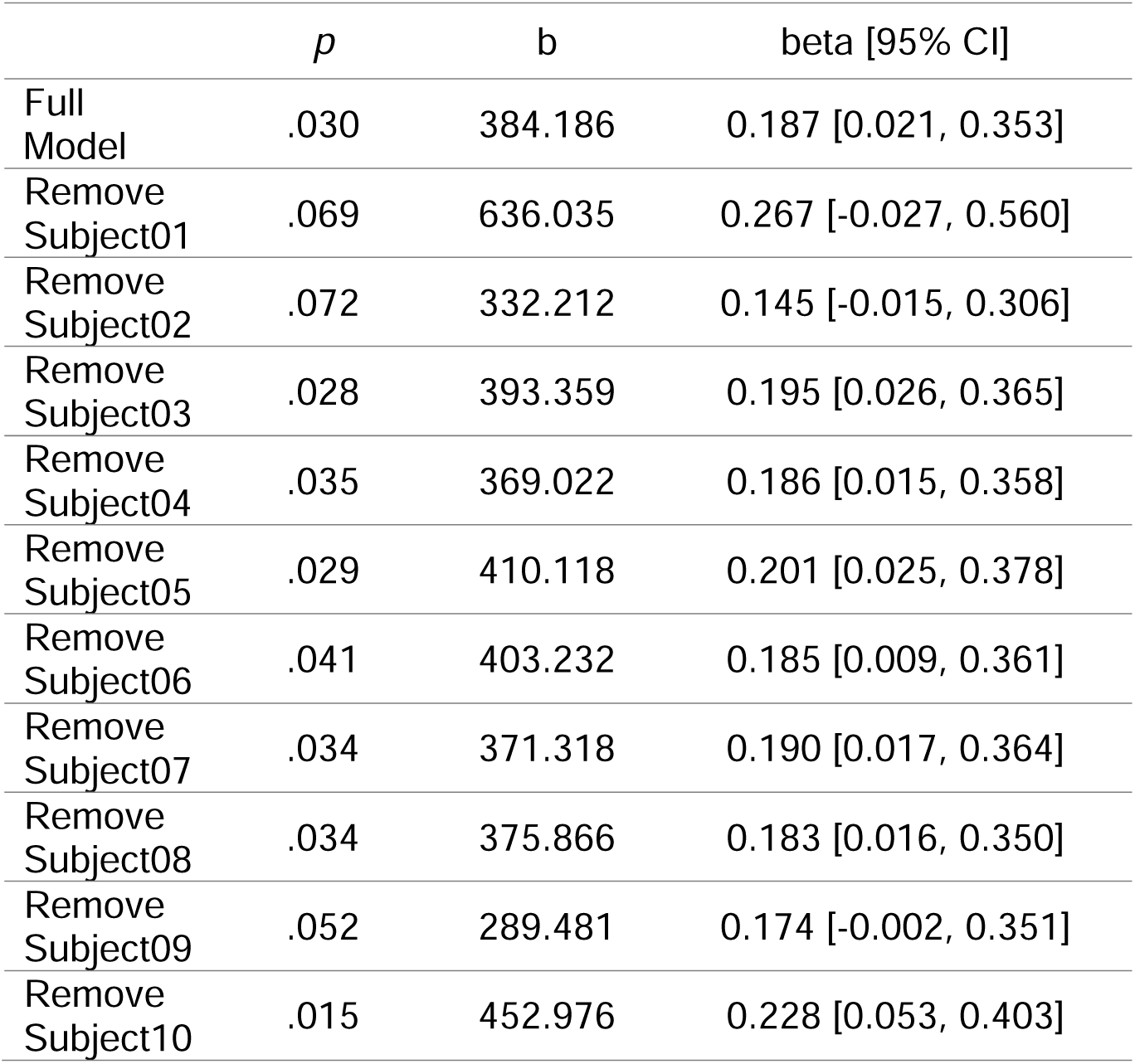
Statistical outcomes for leave-one-out cross-validation analyses of total intracranial cerebrospinal fluid. Model Subject01 Subject02 Subject03 Subject04 Subject05 Subject06 Subject07 Subject08 Subject09 Subject10

|  | *p* | b | beta [95% CI] |
| --- | --- | --- | --- |
| Full Model | .030 | 384.186 | 0.187 [0.021, 0.353] |
| Remove Subject01 | .069 | 636.035 | 0.267 [-0.027, 0.560] |
| Remove Subject02 | .072 | 332.212 | 0.145 [-0.015, 0.306] |
| Remove Subject03 | .028 | 393.359 | 0.195 [0.026, 0.365] |
| Remove Subject04 | .035 | 369.022 | 0.186 [0.015, 0.358] |
| Remove Subject05 | .029 | 410.118 | 0.201 [0.025, 0.378] |
| Remove Subject06 | .041 | 403.232 | 0.185 [0.009, 0.361] |
| Remove Subject07 | .034 | 371.318 | 0.190 [0.017, 0.364] |
| Remove Subject08 | .034 | 375.866 | 0.183 [0.016, 0.350] |
| Remove Subject09 | .052 | 289.481 | 0.174 [-0.002, 0.351] |
| Remove Subject10 | .015 | 452.976 | 0.228 [0.053, 0.403] |

**Table S5.** Statistical outcomes of linear mixed effects models that were implemented for the analysis of total intracranial cerebrospinal fluid computed with different voxel-level thresholds. Intracranial cerebrospinal fluid was assessed against gestational week to evaluate changes during pregnancy.

| Threshold | $p$ | b | beta [95% CI] |
| --- | --- | --- | --- |
| 60% | .016 | 518.510 | 0.206 [0.045, 0.367] |
| 65% | .019 | 486.391 | 0.201 [0.039, 0.363] |
| 70% | .022 | 451.744 | 0.196 [0.033, 0.359] |
| 75% | .025 | 420.094 | 0.192 [0.028, 0.356] |
| 80% | .030 | 384.186 | 0.187 [0.021, 0.353] |
| 85% | .034 | 348.885 | 0.184 [0.016, 0.351] |
| 90% | .040 | 311.048 | 0.181 [0.010, 0.352] |
| 95% | .030 | 284.357 | 0.192 [0.022, 0.362] |

**Fig. S3.**
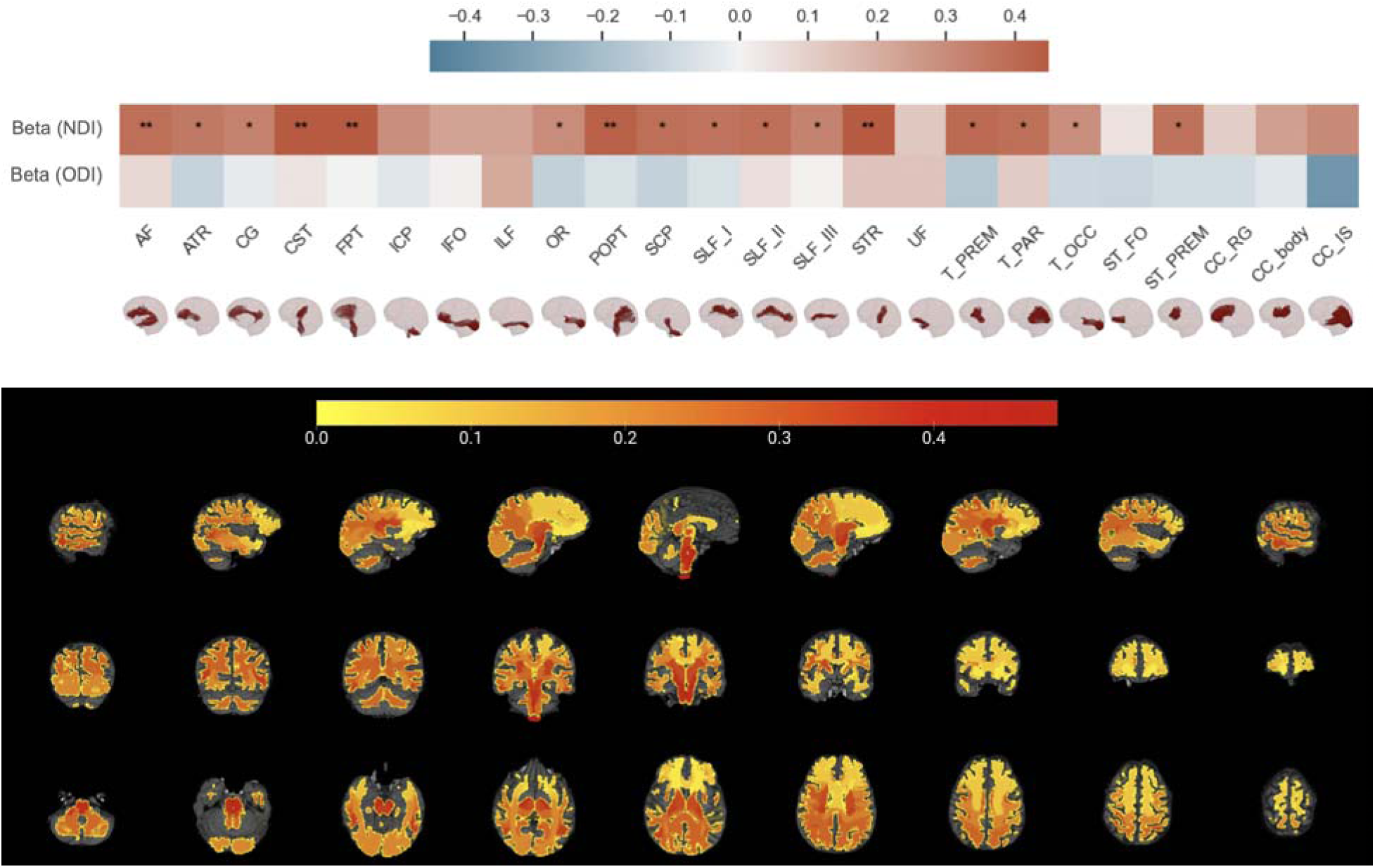
Illustration of the association between neurite density index and gestational week. The upper panel depicts a heatmap of standardized regression coefficients obtained from linear mixed effects (LME) models that were fitted to assess the association between neurite density index and gestation week. The below panel showcases a whole-brain tract map, color-coded to represent the standardized regression coefficients from LME models fitted for neurite density index against gestational age. NDI: neurite density index, ODI: orientation dispersion index, AF: Arcuate Fascicle, ATR: Anterior Thalamic Radiation, CG: Cingulum, CST: Corticospinal Tract, FPT: Fronto-Pontine Tract, ICP: Inferior Cerebellar Peduncle, IFO: Inferior Occipito-Frontal Fascicle, ILF: Inferior Longitudinal Fascicle, OR: Optic Radiation, POPT: Parieto-Occipital Pontine, SCP: Superior Cerebellar Peduncle, SLF_I: Superior Longitudinal Fascicle I; SLF_II: Superior Longitudinal Fascicle II, SLF_III: Superior Longitudinal Fascicle III, STR: Superior Thalamic Radiation, UF: Uncinate Fascicle, T_PREM: Thalamo-Premotor, T_PAR: Thalamo-Parietal, T_OCC: Thalamo-Occipital, ST_FO: Striato-Fronto-Orbital, ST_PREM: Striato-Premotor, CC_RG: Rostrum and Genu, CC_body: Corpus Callosum Body, CC_IS: Isthmus and Splenium.

**Fig. S4.**
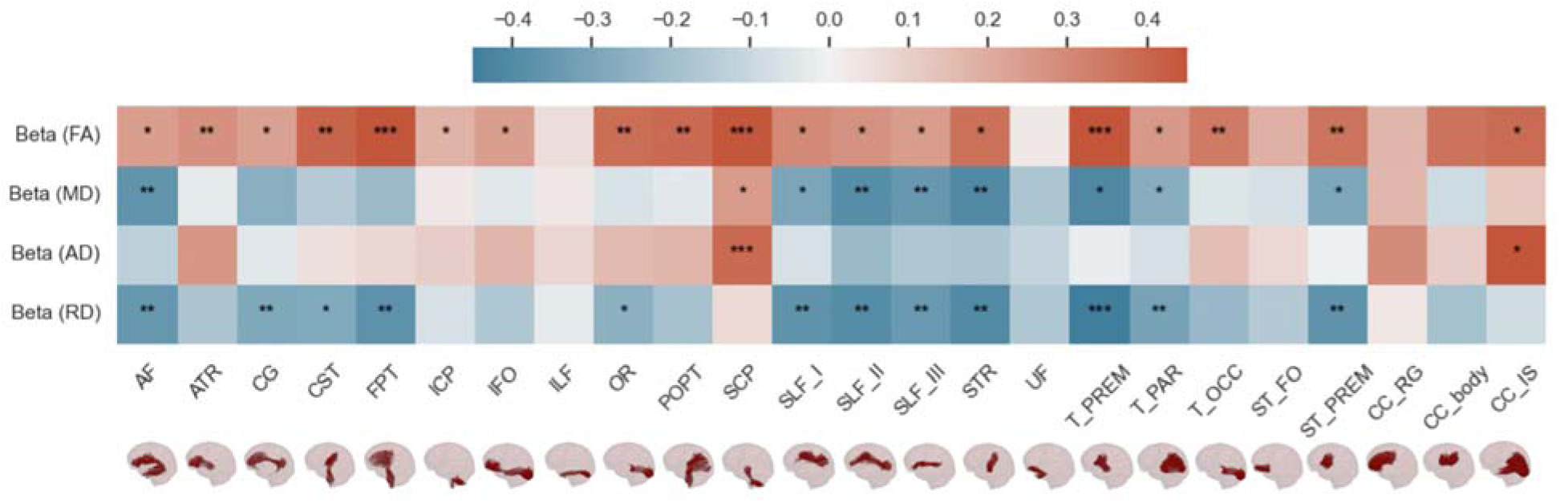
Illustration of the association between diffusion tensor imaging metrics and gestational week. It depicts a heatmap of standardized regression coefficients obtained from linear mixed effects models that were fitted to assess the relations between diffusion tensor imaging metrics and gestational week. FA: fractional anisotropy, MD: mean diffusivity, AD: axial diffusivity, RD: radial diffusivity, AF: Arcuate Fascicle, ATR: Anterior Thalamic Radiation, CG: Cingulum, CST: Corticospinal Tract, FPT: Fronto-Pontine Tract, ICP: Inferior Cerebellar Peduncle, IFO: Inferior Occipito-Frontal Fascicle, ILF: Inferior Longitudinal Fascicle, OR: Optic Radiation, POPT: Parieto-Occipital Pontine, SCP: Superior Cerebellar Peduncle, SLF_I: Superior Longitudinal Fascicle I; SLF_II: Superior Longitudinal Fascicle II, SLF_III: Superior Longitudinal Fascicle III, STR: Superior Thalamic Radiation, UF: Uncinate Fascicle, T_PREM: Thalamo-Premotor, T_PAR: Thalamo-Parietal, T_OCC: Thalamo-Occipital, ST_FO: Striato-Fronto-Orbital, ST_PREM: Striato-Premotor, CC_RG: Rostrum and Genu, CC_body: Corpus Callosum Body, CC_IS: Isthmus and Splenium.

**Fig. S5.**
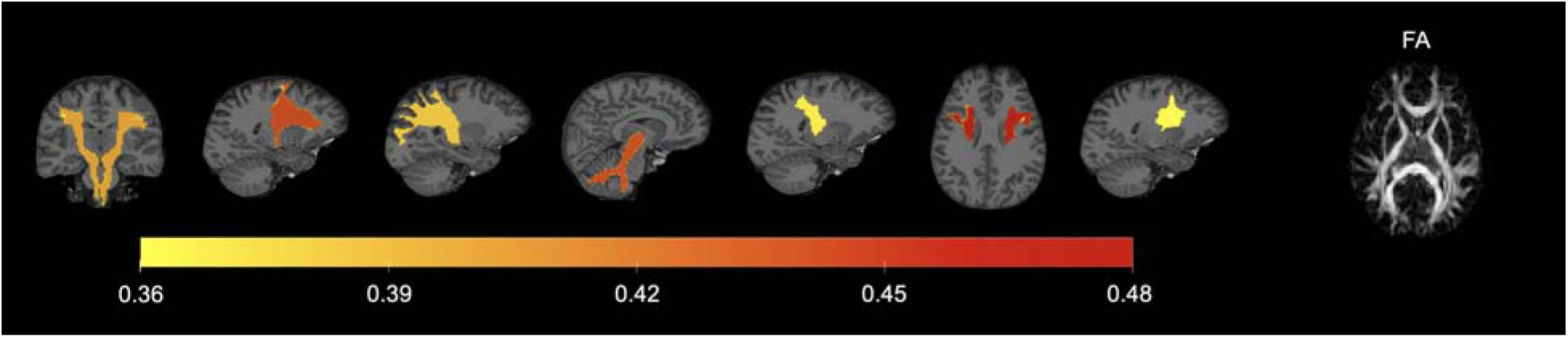
Illustration of the association between fractional anisotropy and gestational week. The color-coding represents the standardized regression coefficients from linear mixed effects models fitted for fractional anisotropy against gestational age. FA: fractional anisotropy.

**Table S6.**
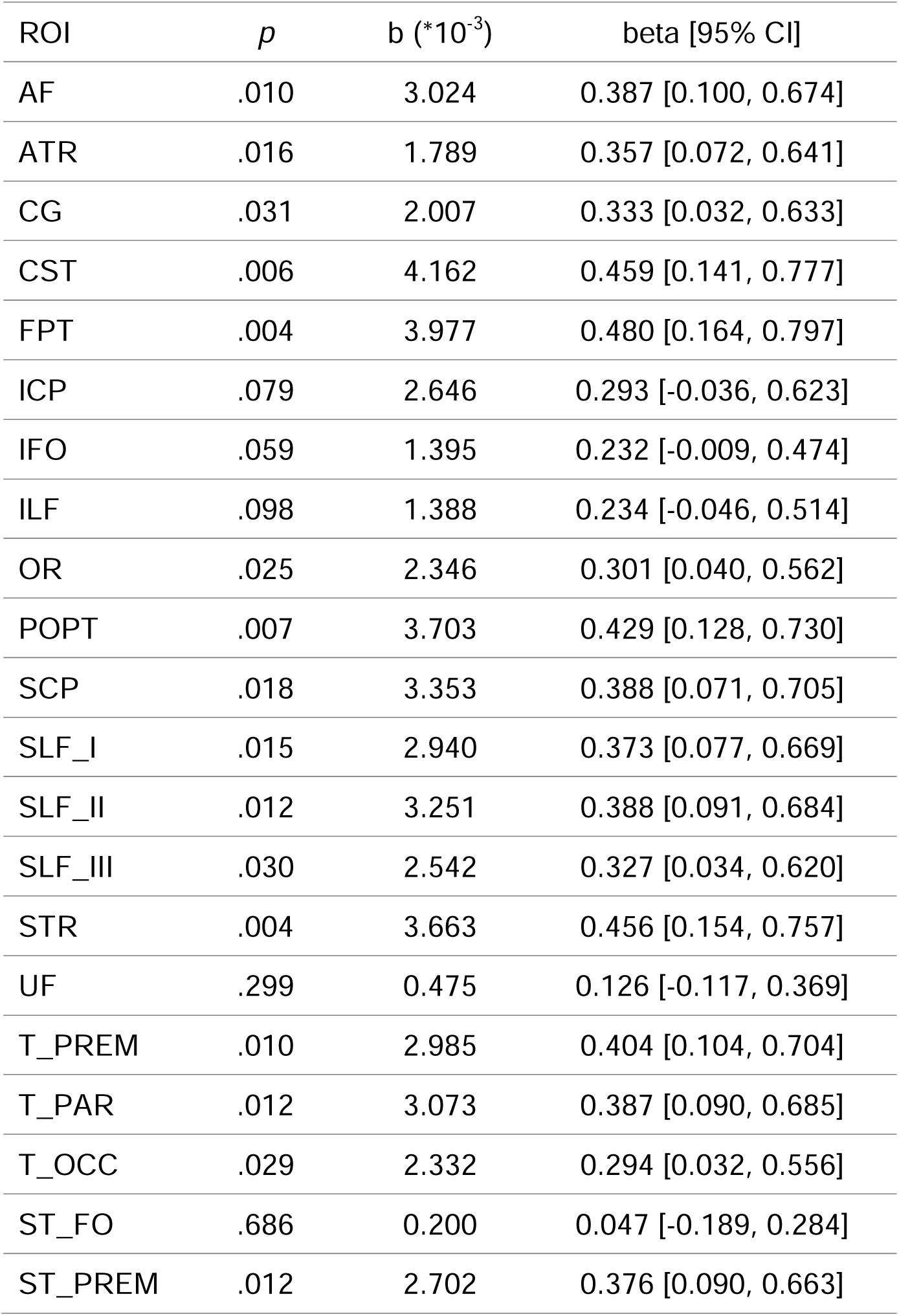
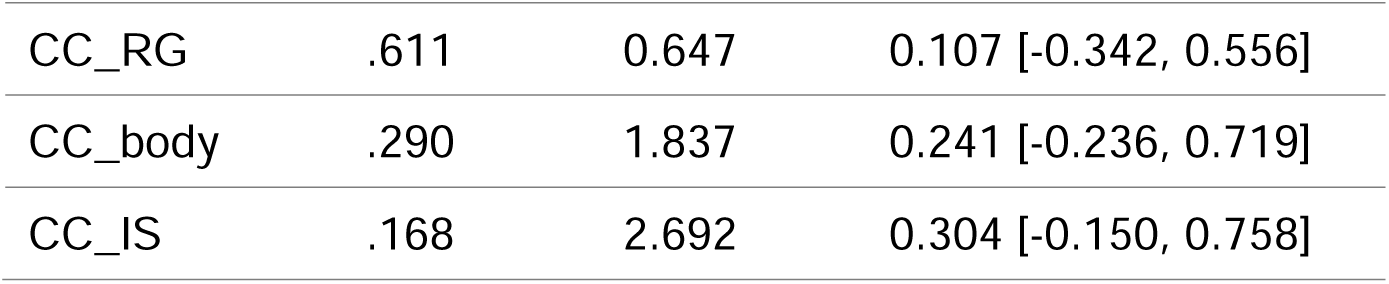
Statistical results of the linear mixed effects models for neurite density index. The models were utilized to assess changes in neurite density index in relation to gestational week throughout the duration of pregnancy. AF: Arcuate Fascicle, ATR: Anterior Thalamic Radiation, CG: Cingulum, CST: Corticospinal Tract, FPT: Fronto-Pontine Tract, ICP: Inferior Cerebellar Peduncle, IFO: Inferior Occipito-Frontal Fascicle, ILF: Inferior Longitudinal Fascicle, OR: Optic Radiation, POPT: Parieto-Occipital Pontine, SCP: Superior Cerebellar Peduncle, SLF_I: Superior Longitudinal Fascicle I; SLF_II: Superior Longitudinal Fascicle II, SLF_III: Superior Longitudinal Fascicle III, STR: Superior Thalamic Radiation, UF: Uncinate Fascicle, T_PREM: Thalamo-Premotor, T_PAR: Thalamo-Parietal, T_OCC: Thalamo-Occipital, ST_FO: Striato-Fronto-Orbital, ST_PREM: Striato-Premotor, CC_RG: Rostrum and Genu, CC_body: Corpus Callosum Body, CC_IS: Isthmus and Splenium. ROI: region of interest.

| ROI | *p* | b (*10<sup>-3</sup>) | beta [95% CI] |
| --- | --- | --- | --- |
| AF | .010 | 3.024 | 0.387 [0.100, 0.674] |
| ATR | .016 | 1.789 | 0.357 [0.072, 0.641] |
| CG | .031 | 2.007 | 0.333 [0.032, 0.633] |
| CST | .006 | 4.162 | 0.459 [0.141, 0.777] |
| FPT | .004 | 3.977 | 0.480 [0.164, 0.797] |
| ICP | .079 | 2.646 | 0.293 [-0.036, 0.623] |
| IFO | .059 | 1.395 | 0.232 [-0.009, 0.474] |
| ILF | .098 | 1.388 | 0.234 [-0.046, 0.514] |
| OR | .025 | 2.346 | 0.301 [0.040, 0.562] |
| POPT | .007 | 3.703 | 0.429 [0.128, 0.730] |
| SCP | .018 | 3.353 | 0.388 [0.071, 0.705] |
| SLF_I | .015 | 2.940 | 0.373 [0.077, 0.669] |
| SLF_II | .012 | 3.251 | 0.388 [0.091, 0.684] |
| SLF_III | .030 | 2.542 | 0.327 [0.034, 0.620] |
| STR | .004 | 3.663 | 0.456 [0.154, 0.757] |
| UF | .299 | 0.475 | 0.126 [-0.117, 0.369] |
| T_PREM | .010 | 2.985 | 0.404 [0.104, 0.704] |
| T_PAR | .012 | 3.073 | 0.387 [0.090, 0.685] |
| T_OCC | .029 | 2.332 | 0.294 [0.032, 0.556] |
| ST_FO | .686 | 0.200 | 0.047 [-0.189, 0.284] |
| ST_PREM | .012 | 2.702 | 0.376 [0.090, 0.663] |
| CC_RG | .611 | 0.647 | 0.107 [-0.342, 0.556] |
| CC_body | .290 | 1.837 | 0.241 [-0.236, 0.719] |
| CC_IS | .168 | 2.692 | 0.304 [-0.150, 0.758] |

**Table S7.**
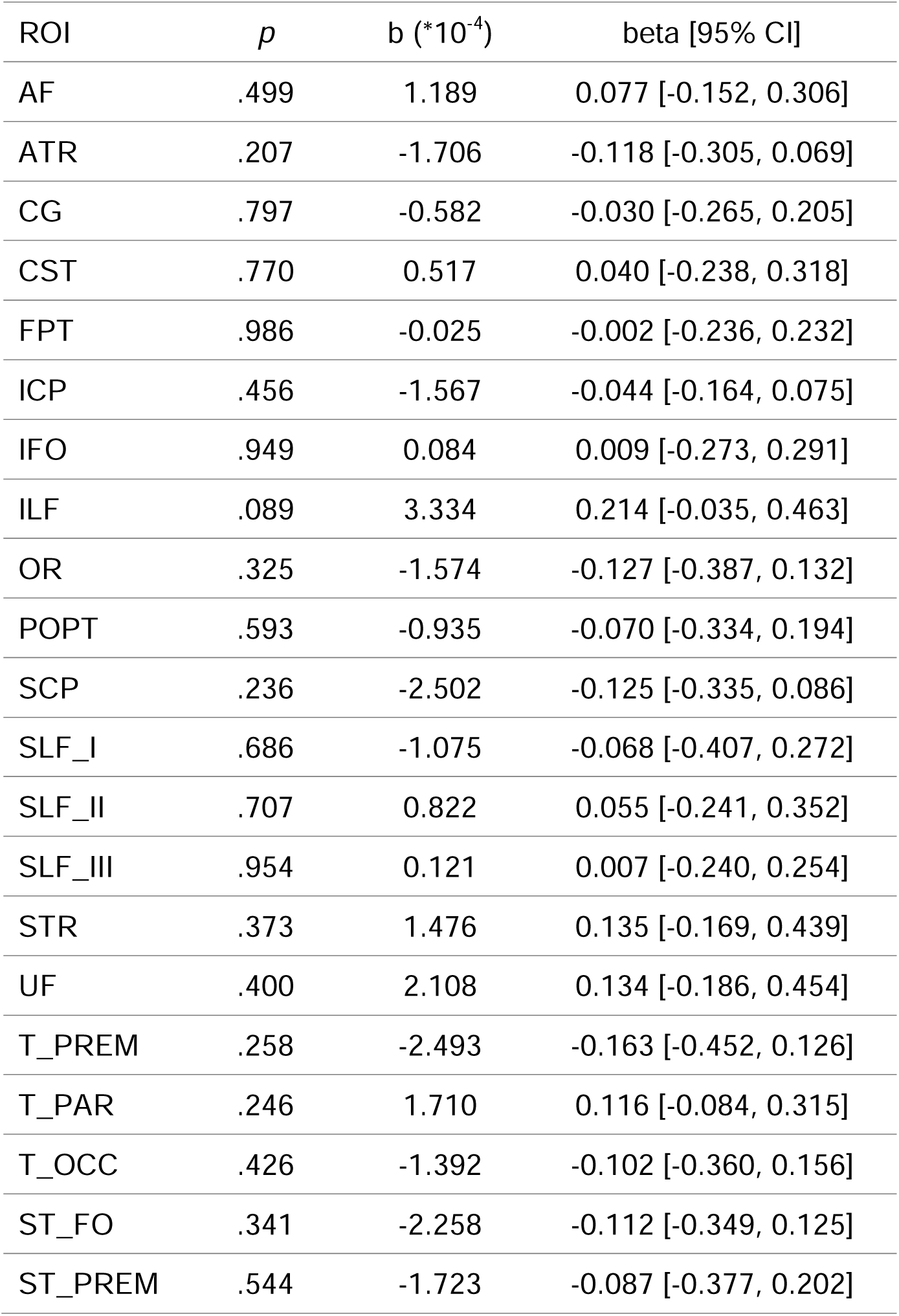
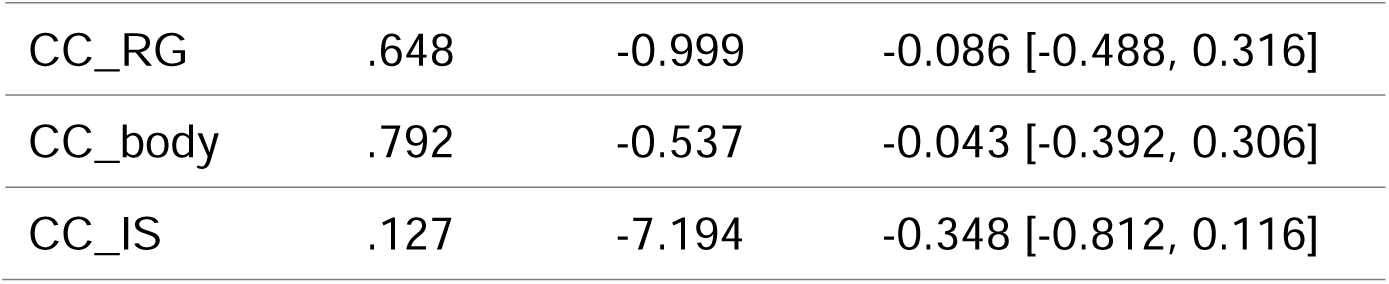
Statistical results of the linear mixed effects models for orientation dispersion index. The models were utilized to assess changes in the orientation dispersion index in relation to gestational week throughout the duration of pregnancy. AF: Arcuate Fascicle, ATR: Anterior Thalamic Radiation, CG: Cingulum, CST: Corticospinal Tract, FPT: Fronto-Pontine Tract, ICP: Inferior Cerebellar Peduncle, IFO: Inferior Occipito-Frontal Fascicle, ILF: Inferior Longitudinal Fascicle, OR: Optic Radiation, POPT: Parieto-Occipital Pontine, SCP: Superior Cerebellar Peduncle, SLF_I: Superior Longitudinal Fascicle I; SLF_II: Superior Longitudinal Fascicle II, SLF_III: Superior Longitudinal Fascicle III, STR: Superior Thalamic Radiation, UF: Uncinate Fascicle, T_PREM: Thalamo-Premotor, T_PAR: Thalamo-Parietal, T_OCC: Thalamo-Occipital, ST_FO: Striato-Fronto-Orbital, ST_PREM: Striato-Premotor, CC_RG: Rostrum and Genu, CC_body: Corpus Callosum Body, CC_IS: Isthmus and Splenium. ROI: region of interest.

| ROI | *p* | *b* (*10<sup>-4</sup>) | beta [95% CI] |
| --- | --- | --- | --- |
| AF | .499 | 1.189 | 0.077 [-0.152, 0.306] |
| ATR | .207 | -1.706 | -0.118 [-0.305, 0.069] |
| CG | .797 | -0.582 | -0.030 [-0.265, 0.205] |
| CST | .770 | 0.517 | 0.040 [-0.238, 0.318] |
| FPT | .986 | -0.025 | -0.002 [-0.236, 0.232] |
| ICP | .456 | -1.567 | -0.044 [-0.164, 0.075] |
| IFO | .949 | 0.084 | 0.009 [-0.273, 0.291] |
| ILF | .089 | 3.334 | 0.214 [-0.035, 0.463] |
| OR | .325 | -1.574 | -0.127 [-0.387, 0.132] |
| POPT | .593 | -0.935 | -0.070 [-0.334, 0.194] |
| SCP | .236 | -2.502 | -0.125 [-0.335, 0.086] |
| SLF_I | .686 | -1.075 | -0.068 [-0.407, 0.272] |
| SLF_II | .707 | 0.822 | 0.055 [-0.241, 0.352] |
| SLF_III | .954 | 0.121 | 0.007 [-0.240, 0.254] |
| STR | .373 | 1.476 | 0.135 [-0.169, 0.439] |
| UF | .400 | 2.108 | 0.134 [-0.186, 0.454] |
| T_PREM | .258 | -2.493 | -0.163 [-0.452, 0.126] |
| T_PAR | .246 | 1.710 | 0.116 [-0.084, 0.315] |
| T_OCC | .426 | -1.392 | -0.102 [-0.360, 0.156] |
| ST_FO | .341 | -2.258 | -0.112 [-0.349, 0.125] |
| ST_PREM | .544 | -1.723 | -0.087 [-0.377, 0.202] |
| CC_RG | .648 | -0.999 | -0.086 [-0.488, 0.316] |
| CC_body | .792 | -0.537 | -0.043 [-0.392, 0.306] |
| CC_IS | .127 | -7.194 | -0.348 [-0.812, 0.116] |

**Table S8.**
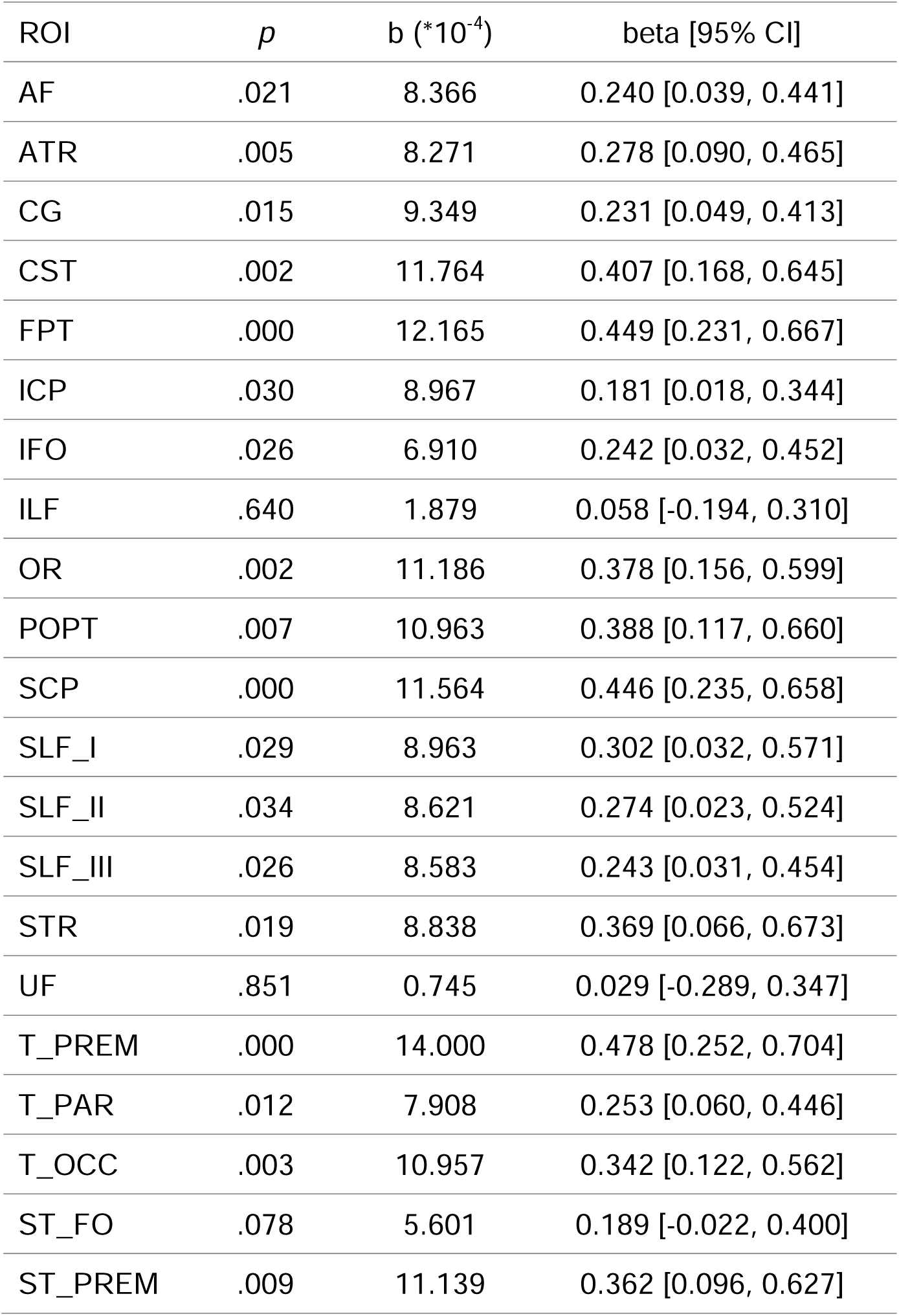
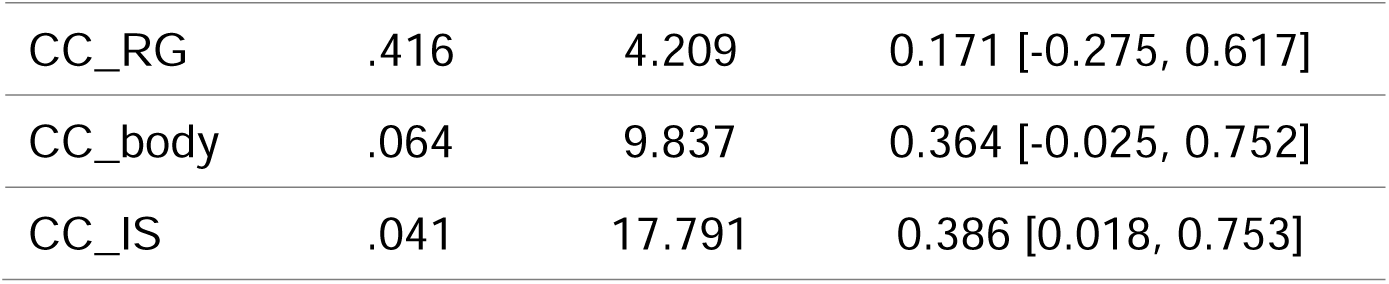
Statistical results of the linear mixed effects models for fractional anisotropy. The models were utilized to assess changes in fractional anisotropy in relation to gestational week throughout the duration of pregnancy. AF: Arcuate Fascicle, ATR: Anterior Thalamic Radiation, CG: Cingulum, CST: Corticospinal Tract, FPT: Fronto-Pontine Tract, ICP: Inferior Cerebellar Peduncle, IFO: Inferior Occipito-Frontal Fascicle, ILF: Inferior Longitudinal Fascicle, OR: Optic Radiation, POPT: Parieto-Occipital Pontine, SCP: Superior Cerebellar Peduncle, SLF_I: Superior Longitudinal Fascicle I; SLF_II: Superior Longitudinal Fascicle II, SLF_III: Superior Longitudinal Fascicle III, STR: Superior Thalamic Radiation, UF: Uncinate Fascicle, T_PREM: Thalamo-Premotor, T_PAR: Thalamo-Parietal, T_OCC: Thalamo-Occipital, ST_FO: Striato-Fronto-Orbital, ST_PREM: Striato-Premotor, CC_RG: Rostrum and Genu, CC_body: Corpus Callosum Body, CC_IS: Isthmus and Splenium. ROI: region of interest.

| ROI | *p* | b (*10<sup>-4</sup>) | beta [95% CI] |
| --- | --- | --- | --- |
| AF | .021 | 8.366 | 0.240 [0.039, 0.441] |
| ATR | .005 | 8.271 | 0.278 [0.090, 0.465] |
| CG | .015 | 9.349 | 0.231 [0.049, 0.413] |
| CST | .002 | 11.764 | 0.407 [0.168, 0.645] |
| FPT | .000 | 12.165 | 0.449 [0.231, 0.667] |
| ICP | .030 | 8.967 | 0.181 [0.018, 0.344] |
| IFO | .026 | 6.910 | 0.242 [0.032, 0.452] |
| ILF | .640 | 1.879 | 0.058 [-0.194, 0.310] |
| OR | .002 | 11.186 | 0.378 [0.156, 0.599] |
| POPT | .007 | 10.963 | 0.388 [0.117, 0.660] |
| SCP | .000 | 11.564 | 0.446 [0.235, 0.658] |
| SLF_I | .029 | 8.963 | 0.302 [0.032, 0.571] |
| SLF_II | .034 | 8.621 | 0.274 [0.023, 0.524] |
| SLF_III | .026 | 8.583 | 0.243 [0.031, 0.454] |
| STR | .019 | 8.838 | 0.369 [0.066, 0.673] |
| UF | .851 | 0.745 | 0.029 [-0.289, 0.347] |
| T_PREM | .000 | 14.000 | 0.478 [0.252, 0.704] |
| T_PAR | .012 | 7.908 | 0.253 [0.060, 0.446] |
| T_OCC | .003 | 10.957 | 0.342 [0.122, 0.562] |
| ST_FO | .078 | 5.601 | 0.189 [-0.022, 0.400] |
| ST_PREM | .009 | 11.139 | 0.362 [0.096, 0.627] |
| CC_RG | .416 | 4.209 | 0.171 [-0.275, 0.617] |
| CC_body | .064 | 9.837 | 0.364 [-0.025, 0.752] |
| CC_IS | .041 | 17.791 | 0.386 [0.018, 0.753] |

**Table S9.**
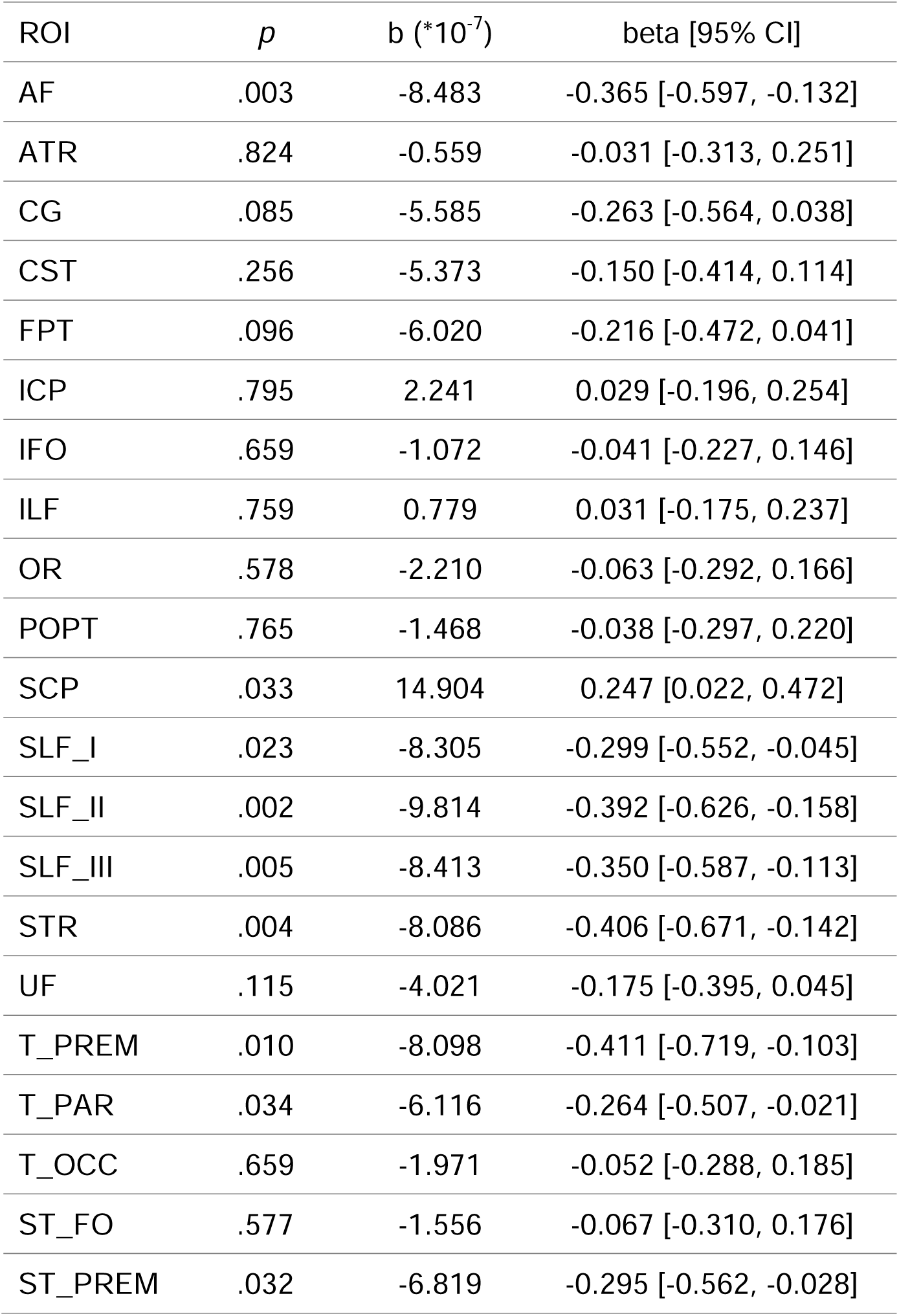
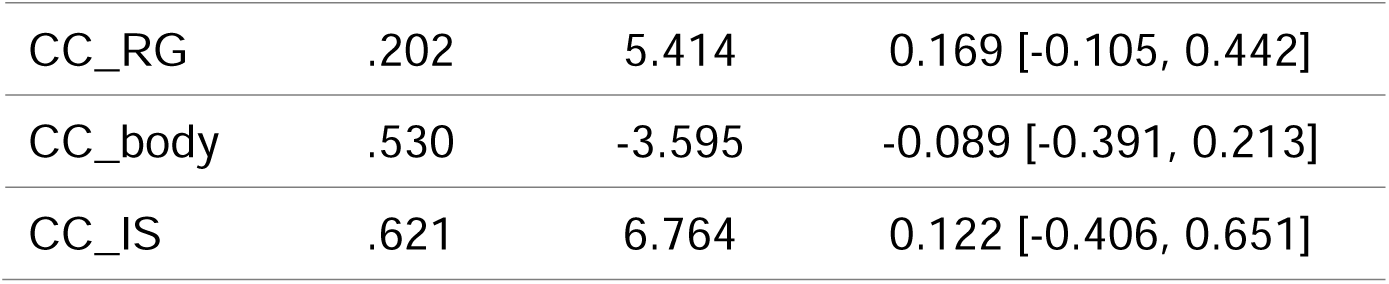
Statistical results of the linear mixed effects models for mean diffusivity. The models were utilized to assess changes in mean diffusivity in relation to gestation week throughout the duration of pregnancy. AF: Arcuate Fascicle, ATR: Anterior Thalamic Radiation, CG: Cingulum, CST: Corticospinal Tract, FPT: Fronto-Pontine Tract, ICP: Inferior Cerebellar Peduncle, IFO: Inferior Occipito-Frontal Fascicle, ILF: Inferior Longitudinal Fascicle, OR: Optic Radiation, POPT: Parieto-Occipital Pontine, SCP: Superior Cerebellar Peduncle, SLF_I: Superior Longitudinal Fascicle I; SLF_II: Superior Longitudinal Fascicle II, SLF_III: Superior Longitudinal Fascicle III, STR: Superior Thalamic Radiation, UF: Uncinate Fascicle, T_PREM: Thalamo-Premotor, T_PAR: Thalamo-Parietal, T_OCC: Thalamo-Occipital, ST_FO: Striato-Fronto-Orbital, ST_PREM: Striato-Premotor, CC_RG: Rostrum and Genu, CC_body: Corpus Callosum Body, CC_IS: Isthmus and Splenium. ROI: region of interest.

| ROI | *p* | *b* (*10<sup>-7</sup>) | beta [95% CI] |
| --- | --- | --- | --- |
| AF | .003 | -8.483 | -0.365 [-0.597, -0.132] |
| ATR | .824 | -0.559 | -0.031 [-0.313, 0.251] |
| CG | .085 | -5.585 | -0.263 [-0.564, 0.038] |
| CST | .256 | -5.373 | -0.150 [-0.414, 0.114] |
| FPT | .096 | -6.020 | -0.216 [-0.472, 0.041] |
| ICP | .795 | 2.241 | 0.029 [-0.196, 0.254] |
| IFO | .659 | -1.072 | -0.041 [-0.227, 0.146] |
| ILF | .759 | 0.779 | 0.031 [-0.175, 0.237] |
| OR | .578 | -2.210 | -0.063 [-0.292, 0.166] |
| POPT | .765 | -1.468 | -0.038 [-0.297, 0.220] |
| SCP | .033 | 14.904 | 0.247 [0.022, 0.472] |
| SLF_I | .023 | -8.305 | -0.299 [-0.552, -0.045] |
| SLF_II | .002 | -9.814 | -0.392 [-0.626, -0.158] |
| SLF_III | .005 | -8.413 | -0.350 [-0.587, -0.113] |
| STR | .004 | -8.086 | -0.406 [-0.671, -0.142] |
| UF | .115 | -4.021 | -0.175 [-0.395, 0.045] |
| T_PREM | .010 | -8.098 | -0.411 [-0.719, -0.103] |
| T_PAR | .034 | -6.116 | -0.264 [-0.507, -0.021] |
| T_OCC | .659 | -1.971 | -0.052 [-0.288, 0.185] |
| ST_FO | .577 | -1.556 | -0.067 [-0.310, 0.176] |
| ST_PREM | .032 | -6.819 | -0.295 [-0.562, -0.028] |
| CC_RG | .202 | 5.414 | 0.169 [-0.105, 0.442] |
| CC_body | .530 | -3.595 | -0.089 [-0.391, 0.213] |
| CC_IS | .621 | 6.764 | 0.122 [-0.406, 0.651] |

**Table S10.**
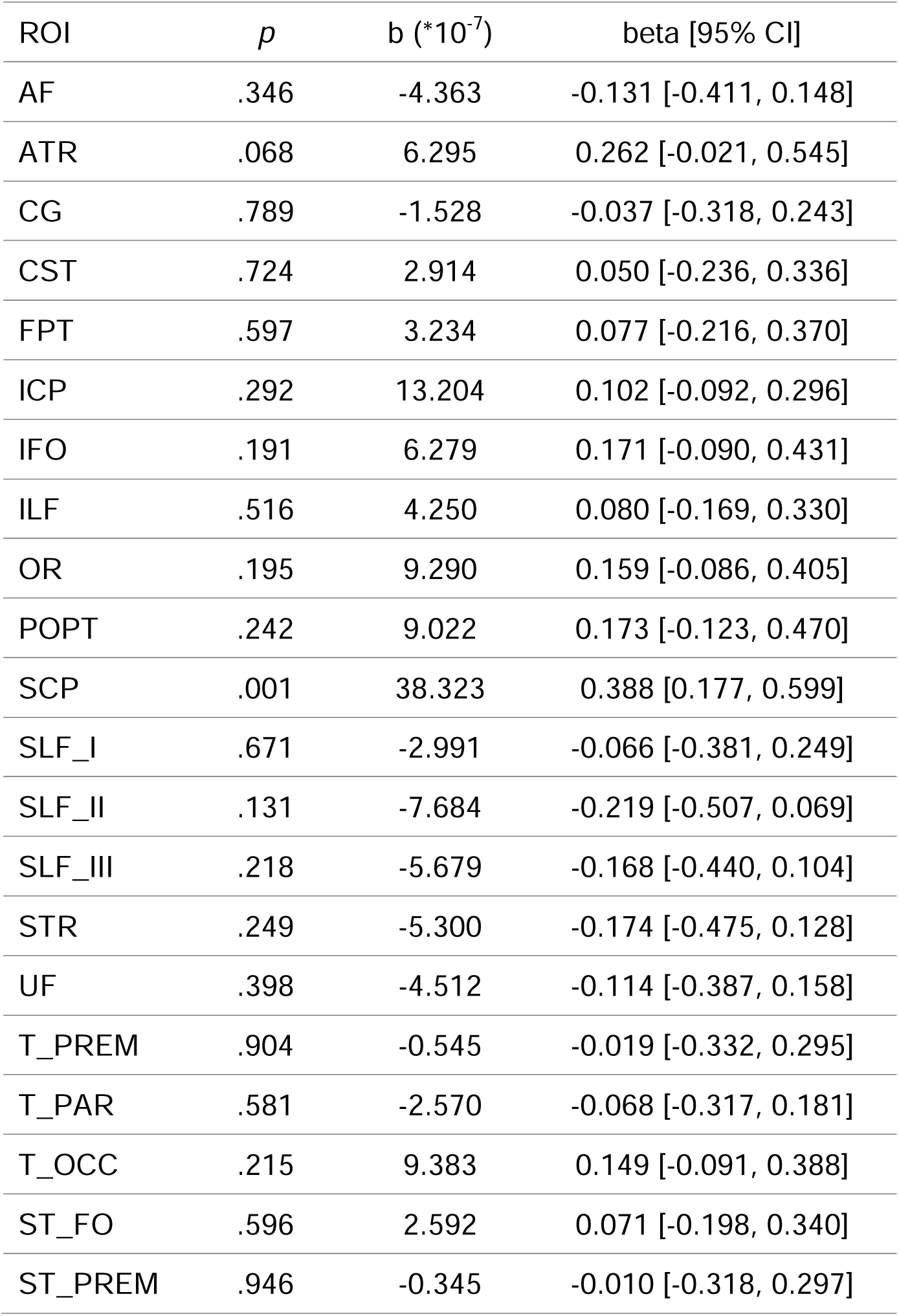
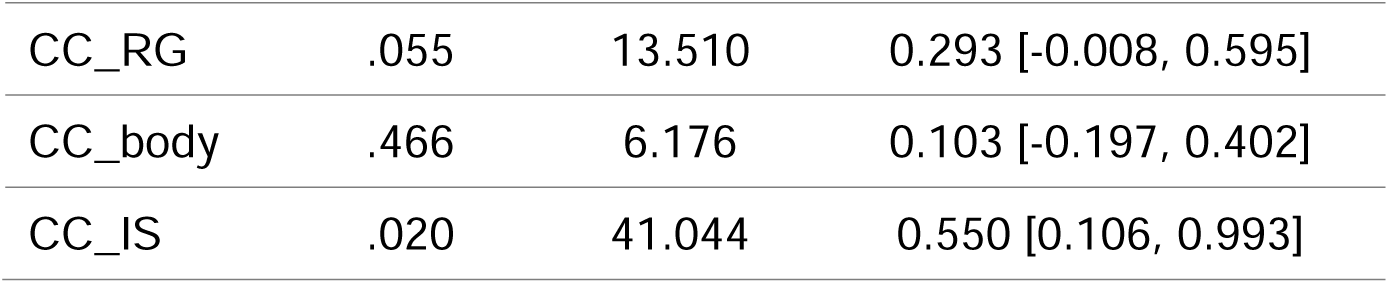
Statistical results of the linear mixed effects models for axial diffusivity. The models were utilized to assess changes in axial diffusivity in relation to gestation week throughout the duration of pregnancy. AF: Arcuate Fascicle, ATR: Anterior Thalamic Radiation, CG: Cingulum, CST: Corticospinal Tract, FPT: Fronto-Pontine Tract, ICP: Inferior Cerebellar Peduncle, IFO: Inferior Occipito-Frontal Fascicle, ILF: Inferior Longitudinal Fascicle, OR: Optic Radiation, POPT: Parieto-Occipital Pontine, SCP: Superior Cerebellar Peduncle, SLF_I: Superior Longitudinal Fascicle I; SLF_II: Superior Longitudinal Fascicle II, SLF_III: Superior Longitudinal Fascicle III, STR: Superior Thalamic Radiation, UF: Uncinate Fascicle, T_PREM: Thalamo-Premotor, T_PAR: Thalamo-Parietal, T_OCC: Thalamo-Occipital, ST_FO: Striato-Fronto-Orbital, ST_PREM: Striato-Premotor, CC_RG: Rostrum and Genu, CC_body: Corpus Callosum Body, CC_IS: Isthmus and Splenium. ROI: region of interest.

| ROI | *p* | b (*10<sup>-7</sup>) | beta [95% CI] |
| --- | --- | --- | --- |
| AF | .346 | -4.363 | -0.131 [-0.411, 0.148] |
| ATR | .068 | 6.295 | 0.262 [-0.021, 0.545] |
| CG | .789 | -1.528 | -0.037 [-0.318, 0.243] |
| CST | .724 | 2.914 | 0.050 [-0.236, 0.336] |
| FPT | .597 | 3.234 | 0.077 [-0.216, 0.370] |
| ICP | .292 | 13.204 | 0.102 [-0.092, 0.296] |
| IFO | .191 | 6.279 | 0.171 [-0.090, 0.431] |
| ILF | .516 | 4.250 | 0.080 [-0.169, 0.330] |
| OR | .195 | 9.290 | 0.159 [-0.086, 0.405] |
| POPT | .242 | 9.022 | 0.173 [-0.123, 0.470] |
| SCP | .001 | 38.323 | 0.388 [0.177, 0.599] |
| SLF_I | .671 | -2.991 | -0.066 [-0.381, 0.249] |
| SLF_II | .131 | -7.684 | -0.219 [-0.507, 0.069] |
| SLF_III | .218 | -5.679 | -0.168 [-0.440, 0.104] |
| STR | .249 | -5.300 | -0.174 [-0.475, 0.128] |
| UF | .398 | -4.512 | -0.114 [-0.387, 0.158] |
| T_PREM | .904 | -0.545 | -0.019 [-0.332, 0.295] |
| T_PAR | .581 | -2.570 | -0.068 [-0.317, 0.181] |
| T_OCC | .215 | 9.383 | 0.149 [-0.091, 0.388] |
| ST_FO | .596 | 2.592 | 0.071 [-0.198, 0.340] |
| ST_PREM | .946 | -0.345 | -0.010 [-0.318, 0.297] |
| CC_RG | .055 | 13.510 | 0.293 [-0.008, 0.595] |
| CC_body | .466 | 6.176 | 0.103 [-0.197, 0.402] |
| CC_IS | .020 | 41.044 | 0.550 [0.106, 0.993] |

**Table S11.**
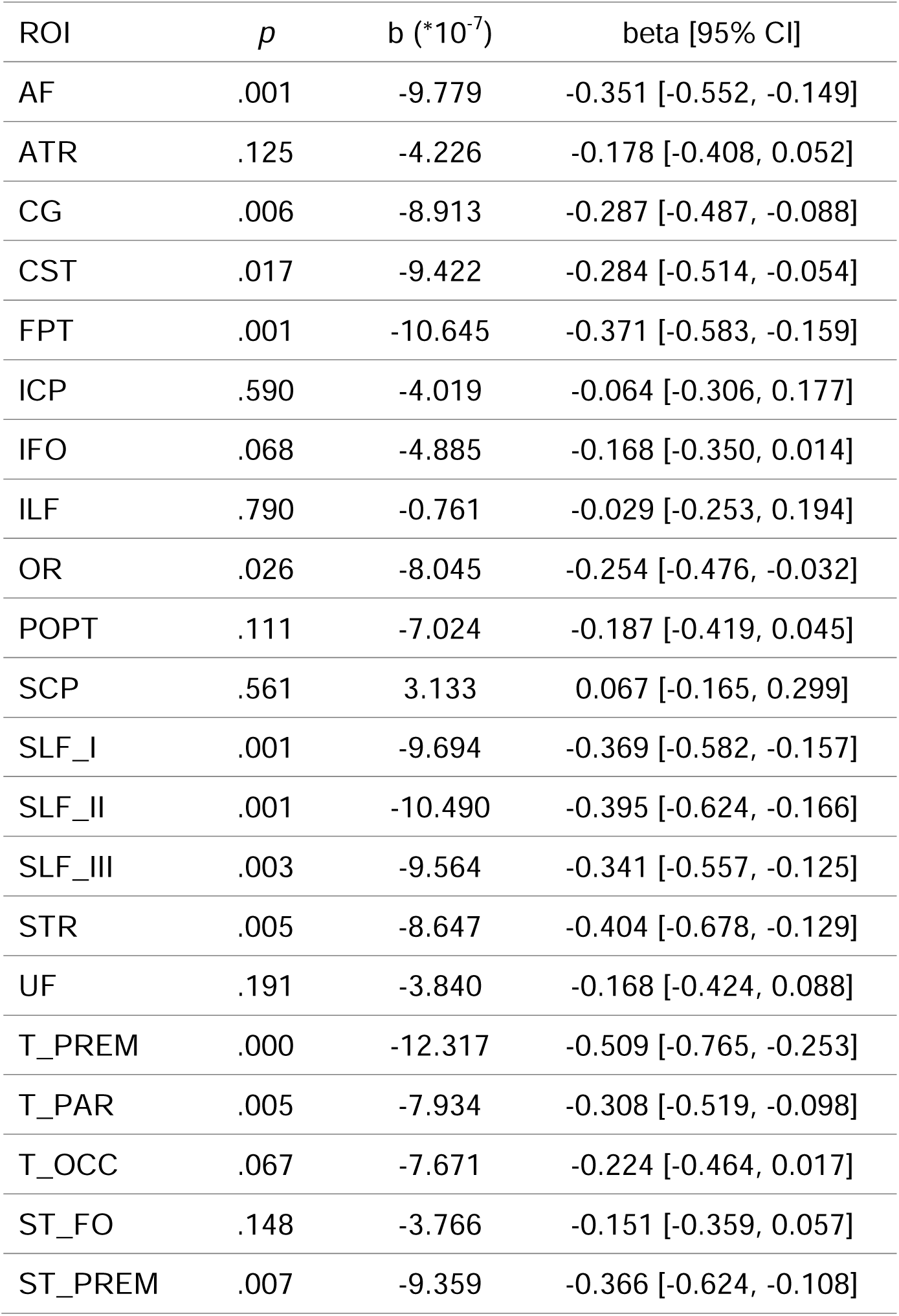
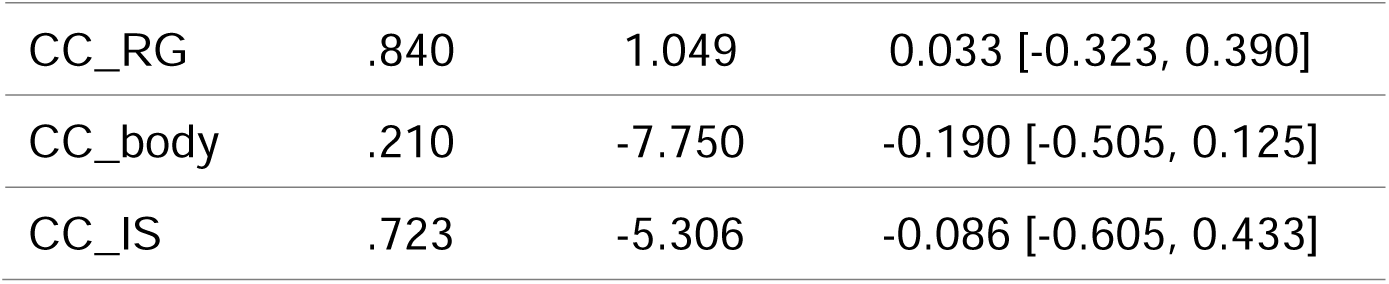
Statistical results of the linear mixed effects models for radial diffusivity. The models were utilized to assess changes in radial diffusivity in relation to gestation week throughout the duration of pregnancy. AF: Arcuate Fascicle, ATR: Anterior Thalamic Radiation, CG: Cingulum, CST: Corticospinal Tract, FPT: Fronto-Pontine Tract, ICP: Inferior Cerebellar Peduncle, IFO: Inferior Occipito-Frontal Fascicle, ILF: Inferior Longitudinal Fascicle, OR: Optic Radiation, POPT: Parieto-Occipital Pontine, SCP: Superior Cerebellar Peduncle, SLF_I: Superior Longitudinal Fascicle I; SLF_II: Superior Longitudinal Fascicle II, SLF_III: Superior Longitudinal Fascicle III, STR: Superior Thalamic Radiation, UF: Uncinate Fascicle, T_PREM: Thalamo-Premotor, T_PAR: Thalamo-Parietal, T_OCC: Thalamo-Occipital, ST_FO: Striato-Fronto-Orbital, ST_PREM: Striato-Premotor, CC_RG: Rostrum and Genu, CC_body: Corpus Callosum Body, CC_IS: Isthmus and Splenium. ROI: region of interest.

| ROI | *p* | *b* (*10<sup>-7</sup>) | beta [95% CI] |
| --- | --- | --- | --- |
| AF | .001 | -9.779 | -0.351 [-0.552, -0.149] |
| ATR | .125 | -4.226 | -0.178 [-0.408, 0.052] |
| CG | .006 | -8.913 | -0.287 [-0.487, -0.088] |
| CST | .017 | -9.422 | -0.284 [-0.514, -0.054] |
| FPT | .001 | -10.645 | -0.371 [-0.583, -0.159] |
| ICP | .590 | -4.019 | -0.064 [-0.306, 0.177] |
| IFO | .068 | -4.885 | -0.168 [-0.350, 0.014] |
| ILF | .790 | -0.761 | -0.029 [-0.253, 0.194] |
| OR | .026 | -8.045 | -0.254 [-0.476, -0.032] |
| POPT | .111 | -7.024 | -0.187 [-0.419, 0.045] |
| SCP | .561 | 3.133 | 0.067 [-0.165, 0.299] |
| SLF_I | .001 | -9.694 | -0.369 [-0.582, -0.157] |
| SLF_II | .001 | -10.490 | -0.395 [-0.624, -0.166] |
| SLF_III | .003 | -9.564 | -0.341 [-0.557, -0.125] |
| STR | .005 | -8.647 | -0.404 [-0.678, -0.129] |
| UF | .191 | -3.840 | -0.168 [-0.424, 0.088] |
| T_PREM | .000 | -12.317 | -0.509 [-0.765, -0.253] |
| T_PAR | .005 | -7.934 | -0.308 [-0.519, -0.098] |
| T_OCC | .067 | -7.671 | -0.224 [-0.464, 0.017] |
| ST_FO | .148 | -3.766 | -0.151 [-0.359, 0.057] |
| ST_PREM | .007 | -9.359 | -0.366 [-0.624, -0.108] |
| CC_RG | .840 | 1.049 | 0.033 [-0.323, 0.390] |
| CC_body | .210 | -7.750 | -0.190 [-0.505, 0.125] |
| CC_IS | .723 | -5.306 | -0.086 [-0.605, 0.433] |

**Fig. S6.**
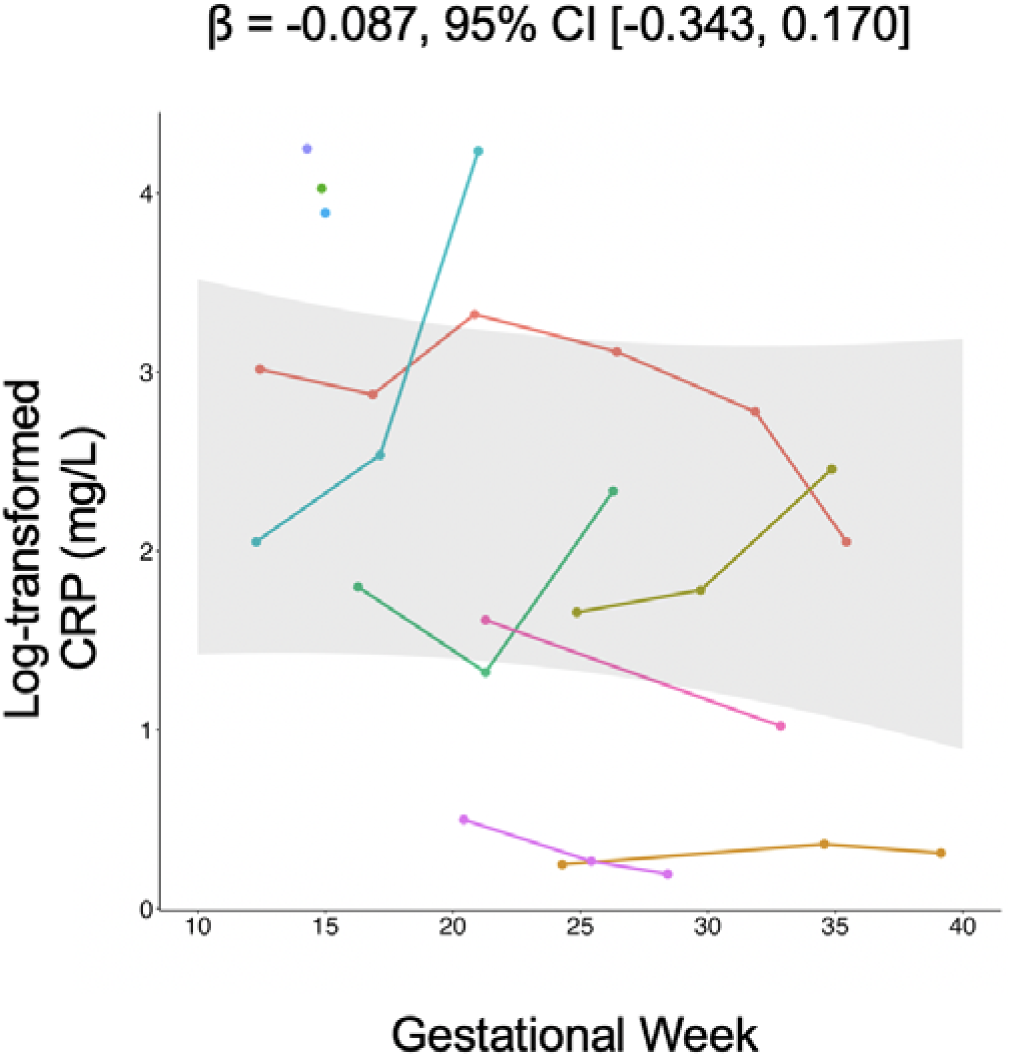
Changes in C-reactive protein levels throughout pregnancy (mg/L). CRP: C-reactive protein.

**Fig. S7.**
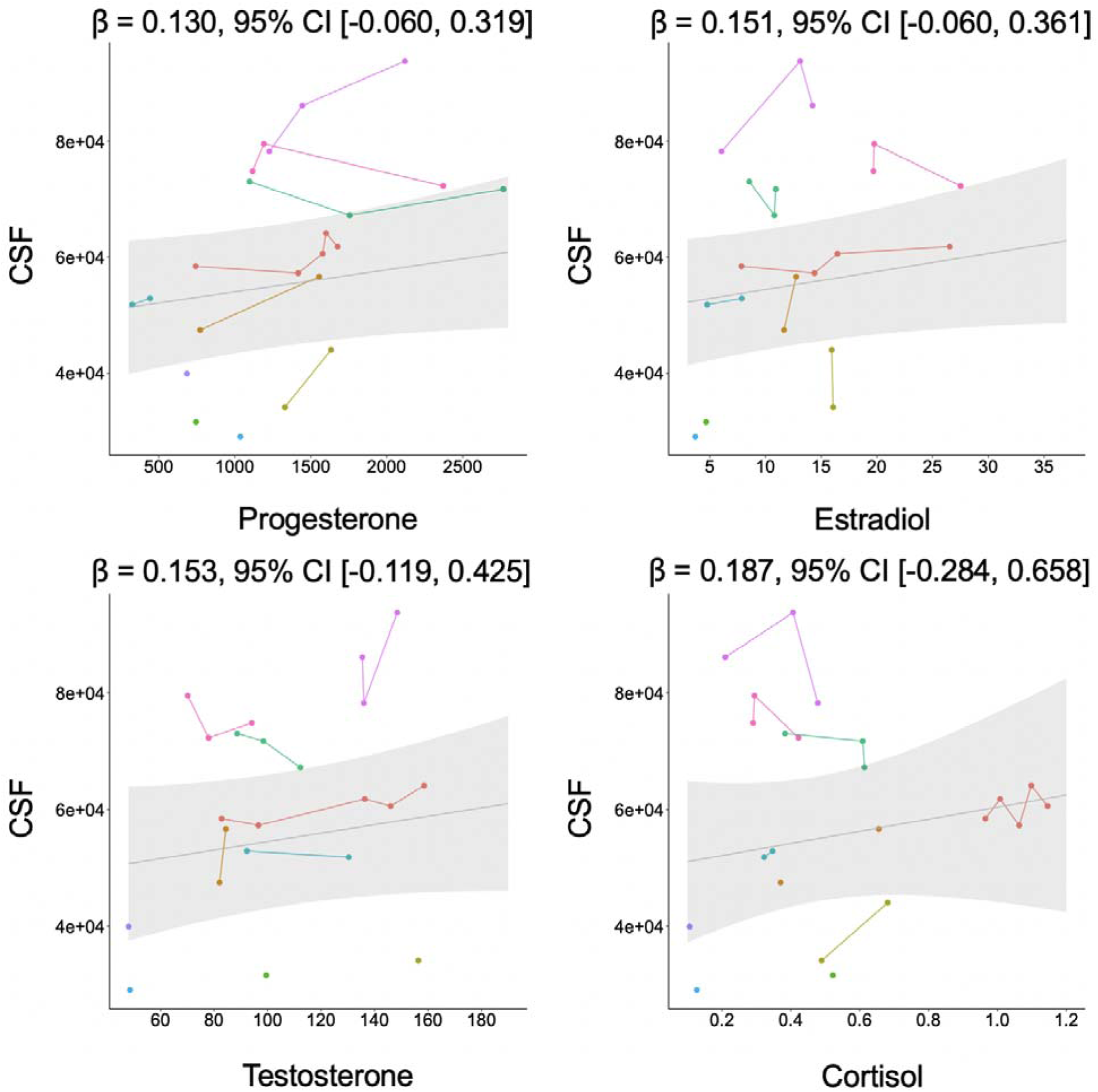
Illustration of the relationship between total intracranial cerebrospinal fluid and salivary hormone levels, specifically progesterone, estradiol, testosterone, and cortisol. CSF: cerebrospinal fluid.

**Fig. S8.**
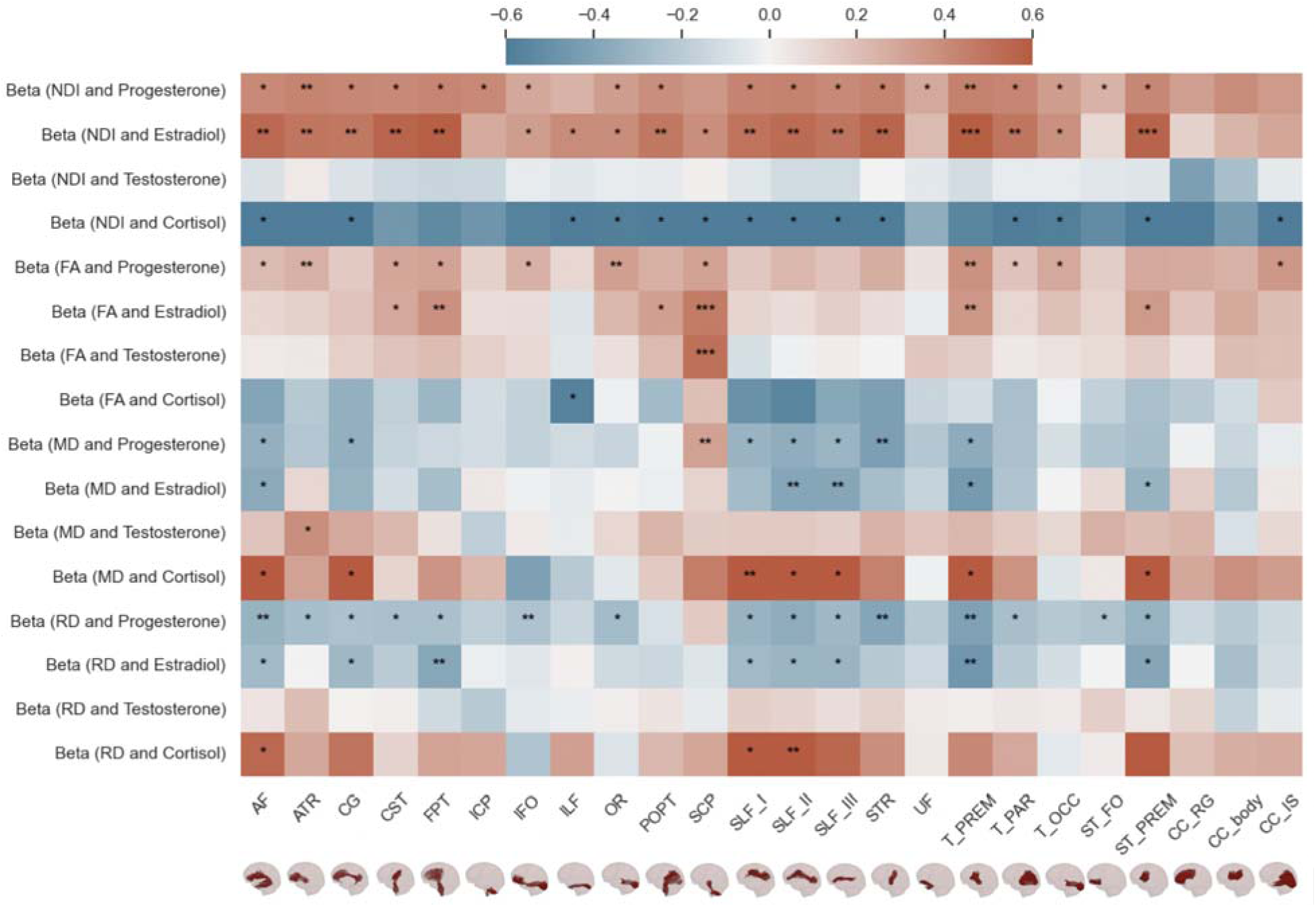
Illustration of the association between white matter metrics and salivary hormone levels. It depicts a heatmap of standardized regression coefficients obtained from linear mixed effects models that were fitted to assess the white matter metrics and salivary hormones (progesterone, estradiol, testosterone, and cortisol). FA: fractional anisotropy, MD: mean diffusivity, AD: axial diffusivity, RD: radial diffusivity, NDI: neurite density index, ODI: orientation dispersion index, AF: Arcuate Fascicle, ATR: Anterior Thalamic Radiation, CG: Cingulum, CST: Corticospinal Tract, FPT: Fronto-Pontine Tract, ICP: Inferior Cerebellar Peduncle, IFO: Inferior Occipito-Frontal Fascicle, ILF: Inferior Longitudinal Fascicle, OR: Optic Radiation, POPT: Parieto-Occipital Pontine, SCP: Superior Cerebellar Peduncle, SLF_I: Superior Longitudinal Fascicle I; SLF_II: Superior Longitudinal Fascicle II, SLF_III: Superior Longitudinal Fascicle III, STR: Superior Thalamic Radiation, UF: Uncinate Fascicle, T_PREM: Thalamo-Premotor, T_PAR: Thalamo-Parietal, T_OCC: Thalamo-Occipital, ST_FO: Striato-Fronto-Orbital, ST_PREM: Striato-Premotor, CC_RG: Rostrum and Genu, CC_body: Corpus Callosum Body, CC_IS: Isthmus and Splenium.

**Table S12.**
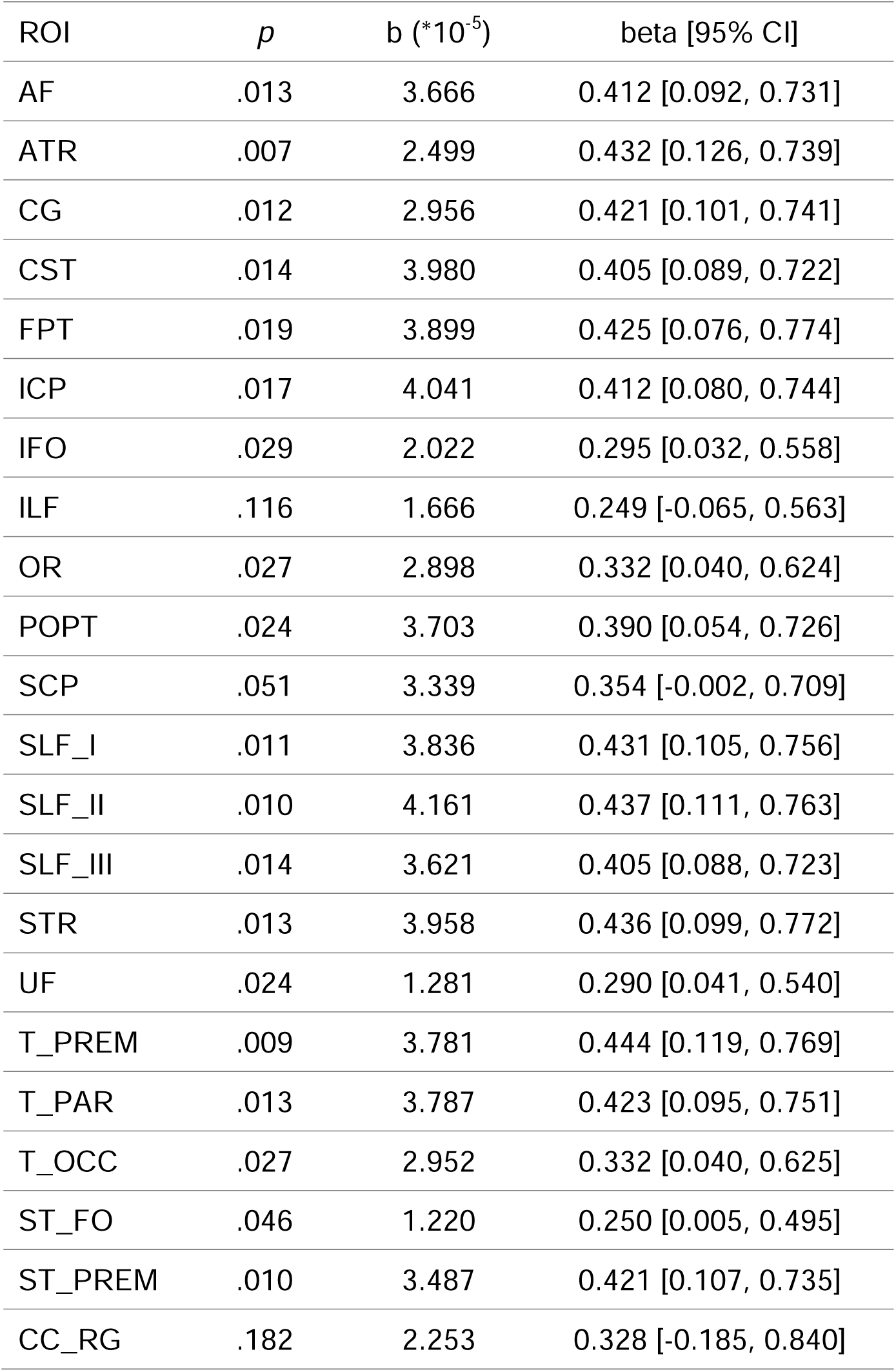
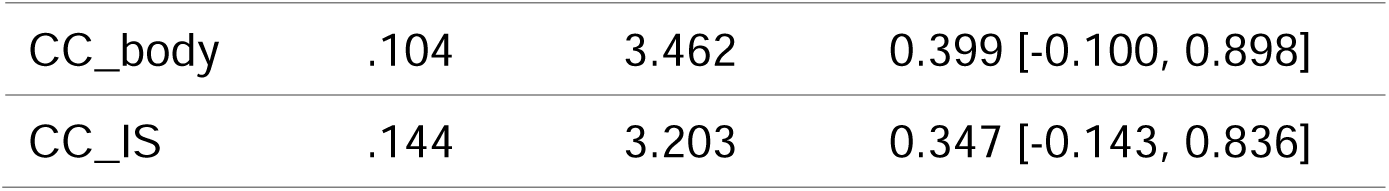
Statistical outcomes of the linear mixed effects models of the neurite density index in relation to salivary progesterone. AF: Arcuate Fascicle, ATR: Anterior Thalamic Radiation, CG: Cingulum, CST: Corticospinal Tract, FPT: Fronto-Pontine Tract, ICP: Inferior Cerebellar Peduncle, IFO: Inferior Occipito-Frontal Fascicle, ILF: Inferior Longitudinal Fascicle, OR: Optic Radiation, POPT: Parieto-Occipital Pontine, SCP: Superior Cerebellar Peduncle, SLF_I: Superior Longitudinal Fascicle I; SLF_II: Superior Longitudinal Fascicle II, SLF_III: Superior Longitudinal Fascicle III, STR: Superior Thalamic Radiation, UF: Uncinate Fascicle, T_PREM: Thalamo-Premotor, T_PAR: Thalamo-Parietal, T_OCC: Thalamo-Occipital, ST_FO: Striato-Fronto-Orbital, ST_PREM: Striato-Premotor, CC_RG: Rostrum and Genu, CC_body: Corpus Callosum Body, CC_IS: Isthmus and Splenium. ROI: region of interest.

| ROI | *p* | b (*10<sup>-5</sup>) | beta [95% CI] |
| --- | --- | --- | --- |
| AF | .013 | 3.666 | 0.412 [0.092, 0.731] |
| ATR | .007 | 2.499 | 0.432 [0.126, 0.739] |
| CG | .012 | 2.956 | 0.421 [0.101, 0.741] |
| CST | .014 | 3.980 | 0.405 [0.089, 0.722] |
| FPT | .019 | 3.899 | 0.425 [0.076, 0.774] |
| ICP | .017 | 4.041 | 0.412 [0.080, 0.744] |
| IFO | .029 | 2.022 | 0.295 [0.032, 0.558] |
| ILF | .116 | 1.666 | 0.249 [-0.065, 0.563] |
| OR | .027 | 2.898 | 0.332 [0.040, 0.624] |
| POPT | .024 | 3.703 | 0.390 [0.054, 0.726] |
| SCP | .051 | 3.339 | 0.354 [-0.002, 0.709] |
| SLF_I | .011 | 3.836 | 0.431 [0.105, 0.756] |
| SLF_II | .010 | 4.161 | 0.437 [0.111, 0.763] |
| SLF_III | .014 | 3.621 | 0.405 [0.088, 0.723] |
| STR | .013 | 3.958 | 0.436 [0.099, 0.772] |
| UF | .024 | 1.281 | 0.290 [0.041, 0.540] |
| T_PREM | .009 | 3.781 | 0.444 [0.119, 0.769] |
| T_PAR | .013 | 3.787 | 0.423 [0.095, 0.751] |
| T_OCC | .027 | 2.952 | 0.332 [0.040, 0.625] |
| ST_FO | .046 | 1.220 | 0.250 [0.005, 0.495] |
| ST_PREM | .010 | 3.487 | 0.421 [0.107, 0.735] |
| CC_RG | .182 | 2.253 | 0.328 [-0.185, 0.840] |
| CC_body | .104 | 3.462 | 0.399 [-0.100, 0.898] |
| CC_IS | .144 | 3.203 | 0.347 [-0.143, 0.836] |

**Table S13.**
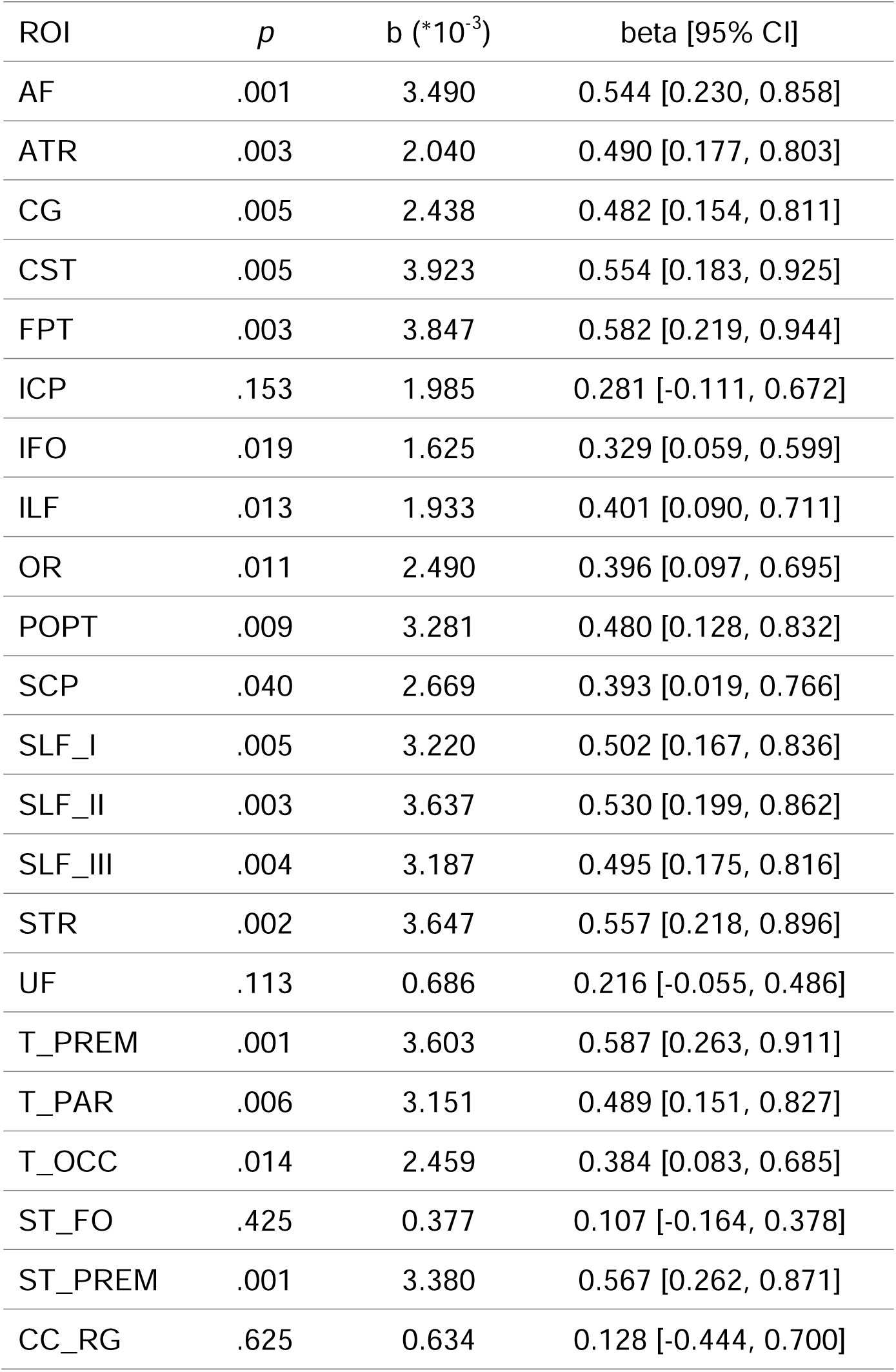
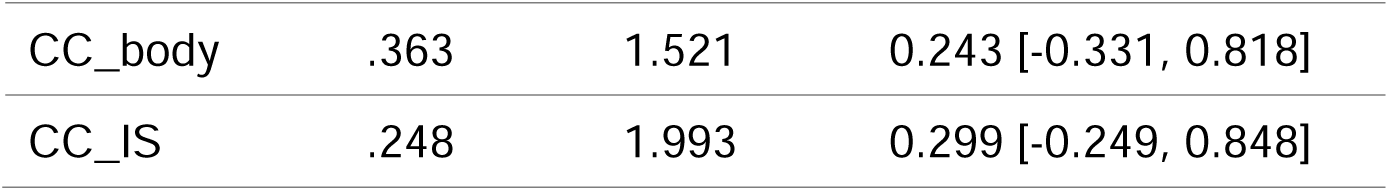
Statistical outcomes of the linear mixed effects models of the neurite density index in relation to salivary estradiol. AF: Arcuate Fascicle, ATR: Anterior Thalamic Radiation, CG: Cingulum, CST: Corticospinal Tract, FPT: Fronto-Pontine Tract, ICP: Inferior Cerebellar Peduncle, IFO: Inferior Occipito-Frontal Fascicle, ILF: Inferior Longitudinal Fascicle, OR: Optic Radiation, POPT: Parieto-Occipital Pontine, SCP: Superior Cerebellar Peduncle, SLF_I: Superior Longitudinal Fascicle I; SLF_II: Superior Longitudinal Fascicle II, SLF_III: Superior Longitudinal Fascicle III, STR: Superior Thalamic Radiation, UF: Uncinate Fascicle, T_PREM: Thalamo-Premotor, T_PAR: Thalamo-Parietal, T_OCC: Thalamo-Occipital, ST_FO: Striato-Fronto-Orbital, ST_PREM: Striato-Premotor, CC_RG: Rostrum and Genu, CC_body: Corpus Callosum Body, CC_IS: Isthmus and Splenium. ROI: region of interest.

| ROI | *p* | b (*10<sup>-3</sup>) | beta [95% CI] |
| --- | --- | --- | --- |
| AF | .001 | 3.490 | 0.544 [0.230, 0.858] |
| ATR | .003 | 2.040 | 0.490 [0.177, 0.803] |
| CG | .005 | 2.438 | 0.482 [0.154, 0.811] |
| CST | .005 | 3.923 | 0.554 [0.183, 0.925] |
| FPT | .003 | 3.847 | 0.582 [0.219, 0.944] |
| ICP | .153 | 1.985 | 0.281 [-0.111, 0.672] |
| IFO | .019 | 1.625 | 0.329 [0.059, 0.599] |
| ILF | .013 | 1.933 | 0.401 [0.090, 0.711] |
| OR | .011 | 2.490 | 0.396 [0.097, 0.695] |
| POPT | .009 | 3.281 | 0.480 [0.128, 0.832] |
| SCP | .040 | 2.669 | 0.393 [0.019, 0.766] |
| SLF_I | .005 | 3.220 | 0.502 [0.167, 0.836] |
| SLF_II | .003 | 3.637 | 0.530 [0.199, 0.862] |
| SLF_III | .004 | 3.187 | 0.495 [0.175, 0.816] |
| STR | .002 | 3.647 | 0.557 [0.218, 0.896] |
| UF | .113 | 0.686 | 0.216 [-0.055, 0.486] |
| T_PREM | .001 | 3.603 | 0.587 [0.263, 0.911] |
| T_PAR | .006 | 3.151 | 0.489 [0.151, 0.827] |
| T_OCC | .014 | 2.459 | 0.384 [0.083, 0.685] |
| ST_FO | .425 | 0.377 | 0.107 [-0.164, 0.378] |
| ST_PREM | .001 | 3.380 | 0.567 [0.262, 0.871] |
| CC_RG | .625 | 0.634 | 0.128 [-0.444, 0.700] |
| CC_body | .363 | 1.521 | 0.243 [-0.331, 0.818] |
| CC_IS | .248 | 1.993 | 0.299 [-0.249, 0.848] |

**Table S14.**
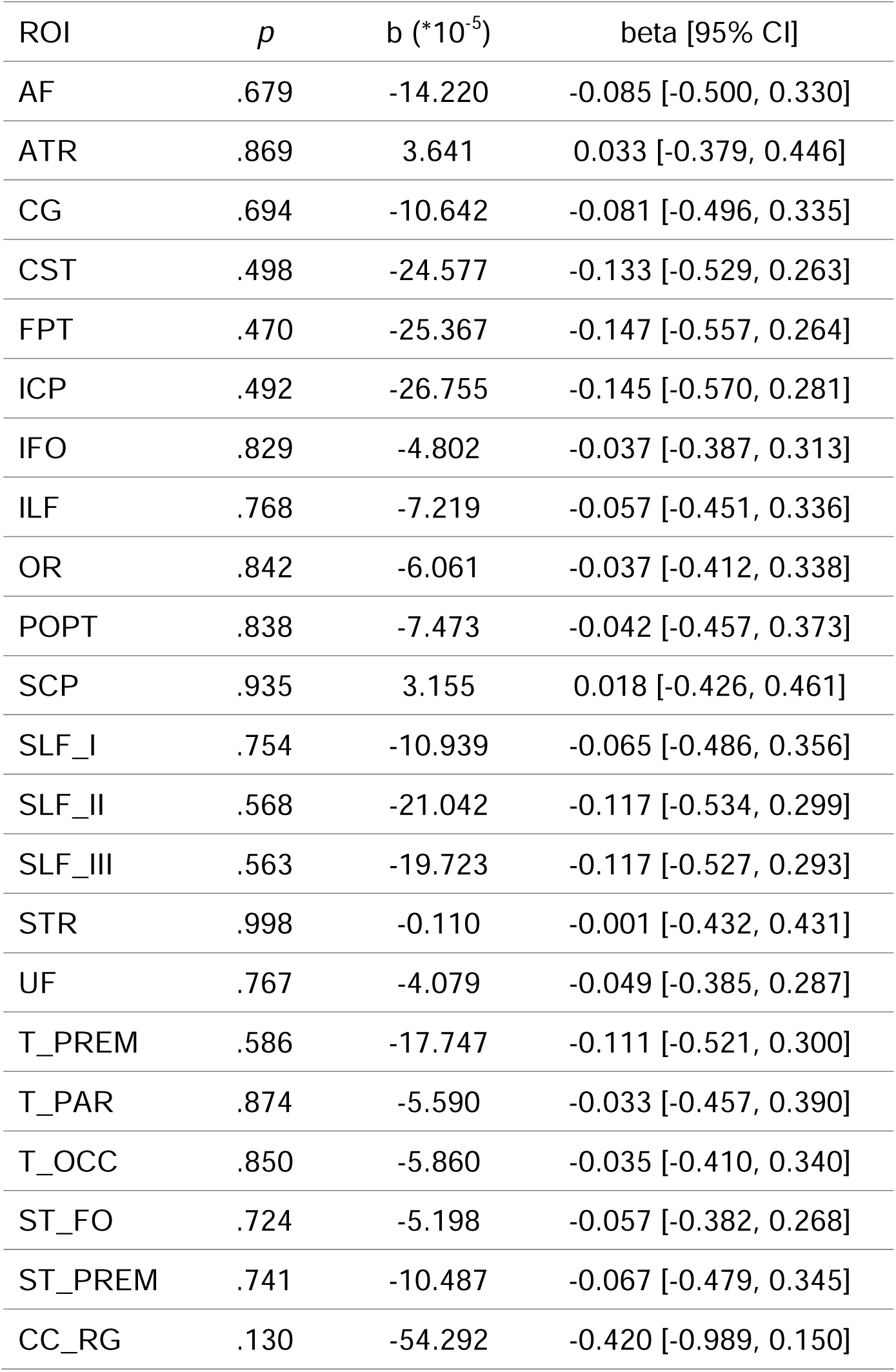
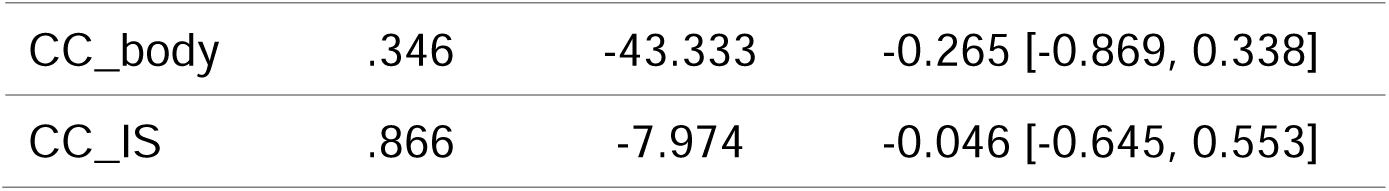
Statistical outcomes of the linear mixed effects models of the neurite density index in relation to salivary testosterone. AF: Arcuate Fascicle, ATR: Anterior Thalamic Radiation, CG: Cingulum, CST: Corticospinal Tract, FPT: Fronto-Pontine Tract, ICP: Inferior Cerebellar Peduncle, IFO: Inferior Occipito-Frontal Fascicle, ILF: Inferior Longitudinal Fascicle, OR: Optic Radiation, POPT: Parieto-Occipital Pontine, SCP: Superior Cerebellar Peduncle, SLF_I: Superior Longitudinal Fascicle I; SLF_II: Superior Longitudinal Fascicle II, SLF_III: Superior Longitudinal Fascicle III, STR: Superior Thalamic Radiation, UF: Uncinate Fascicle, T_PREM: Thalamo-Premotor, T_PAR: Thalamo-Parietal, T_OCC: Thalamo-Occipital, ST_FO: Striato-Fronto-Orbital, ST_PREM: Striato-Premotor, CC_RG: Rostrum and Genu, CC_body: Corpus Callosum Body, CC_IS: Isthmus and Splenium. ROI: region of interest.

| ROI | *p* | b (*10<sup>-5</sup>) | beta [95% CI] |
| --- | --- | --- | --- |
| AF | .679 | -14.220 | -0.085 [-0.500, 0.330] |
| ATR | .869 | 3.641 | 0.033 [-0.379, 0.446] |
| CG | .694 | -10.642 | -0.081 [-0.496, 0.335] |
| CST | .498 | -24.577 | -0.133 [-0.529, 0.263] |
| FPT | .470 | -25.367 | -0.147 [-0.557, 0.264] |
| ICP | .492 | -26.755 | -0.145 [-0.570, 0.281] |
| IFO | .829 | -4.802 | -0.037 [-0.387, 0.313] |
| ILF | .768 | -7.219 | -0.057 [-0.451, 0.336] |
| OR | .842 | -6.061 | -0.037 [-0.412, 0.338] |
| POPT | .838 | -7.473 | -0.042 [-0.457, 0.373] |
| SCP | .935 | 3.155 | 0.018 [-0.426, 0.461] |
| SLF_I | .754 | -10.939 | -0.065 [-0.486, 0.356] |
| SLF_II | .568 | -21.042 | -0.117 [-0.534, 0.299] |
| SLF_III | .563 | -19.723 | -0.117 [-0.527, 0.293] |
| STR | .998 | -0.110 | -0.001 [-0.432, 0.431] |
| UF | .767 | -4.079 | -0.049 [-0.385, 0.287] |
| T_PREM | .586 | -17.747 | -0.111 [-0.521, 0.300] |
| T_PAR | .874 | -5.590 | -0.033 [-0.457, 0.390] |
| T_OCC | .850 | -5.860 | -0.035 [-0.410, 0.340] |
| ST_FO | .724 | -5.198 | -0.057 [-0.382, 0.268] |
| ST_PREM | .741 | -10.487 | -0.067 [-0.479, 0.345] |
| CC_RG | .130 | -54.292 | -0.420 [-0.989, 0.150] |
| CC_body | .346 | -43.333 | -0.265 [-0.869, 0.338] |
| CC_IS | .866 | -7.974 | -0.046 [-0.645, 0.553] |

**Table S15.**
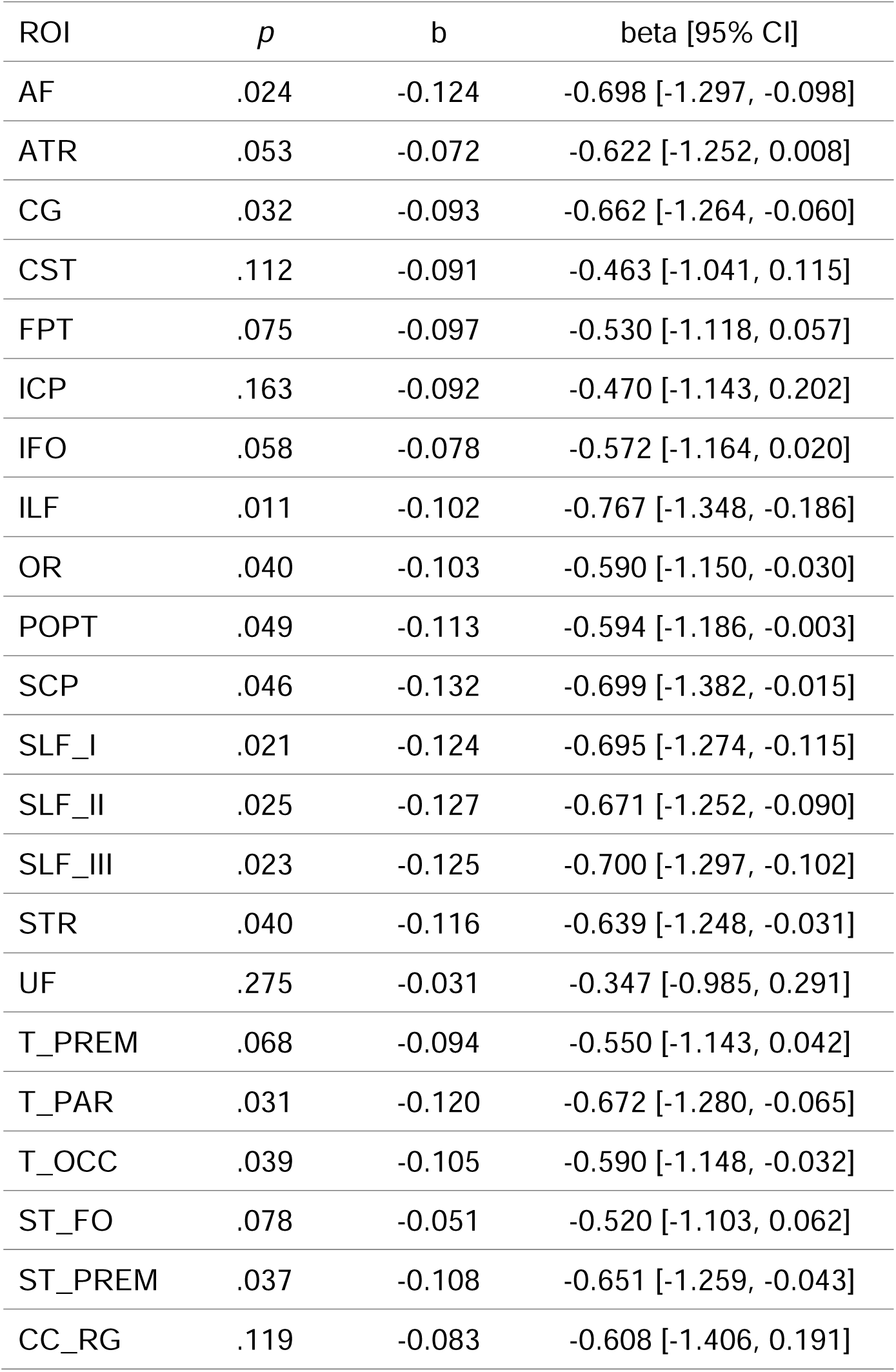
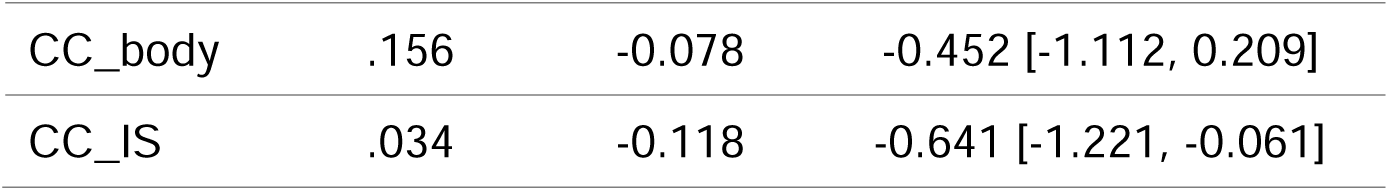
Statistical outcomes of the linear mixed effects models of the neurite density index in relation to salivary cortisol. AF: Arcuate Fascicle, ATR: Anterior Thalamic Radiation, CG: Cingulum, CST: Corticospinal Tract, FPT: Fronto-Pontine Tract, ICP: Inferior Cerebellar Peduncle, IFO: Inferior Occipito-Frontal Fascicle, ILF: Inferior Longitudinal Fascicle, OR: Optic Radiation, POPT: Parieto-Occipital Pontine, SCP: Superior Cerebellar Peduncle, SLF_I: Superior Longitudinal Fascicle I; SLF_II: Superior Longitudinal Fascicle II, SLF_III: Superior Longitudinal Fascicle III, STR: Superior Thalamic Radiation, UF: Uncinate Fascicle, T_PREM: Thalamo-Premotor, T_PAR: Thalamo-Parietal, T_OCC: Thalamo-Occipital, ST_FO: Striato-Fronto-Orbital, ST_PREM: Striato-Premotor, CC_RG: Rostrum and Genu, CC_body: Corpus Callosum Body, CC_IS: Isthmus and Splenium. ROI: region of interest.

| ROI | *p* | b | beta [95% CI] |
| --- | --- | --- | --- |
| AF | .024 | -0.124 | -0.698 [-1.297, -0.098] |
| ATR | .053 | -0.072 | -0.622 [-1.252, 0.008] |
| CG | .032 | -0.093 | -0.662 [-1.264, -0.060] |
| CST | .112 | -0.091 | -0.463 [-1.041, 0.115] |
| FPT | .075 | -0.097 | -0.530 [-1.118, 0.057] |
| ICP | .163 | -0.092 | -0.470 [-1.143, 0.202] |
| IFO | .058 | -0.078 | -0.572 [-1.164, 0.020] |
| ILF | .011 | -0.102 | -0.767 [-1.348, -0.186] |
| OR | .040 | -0.103 | -0.590 [-1.150, -0.030] |
| POPT | .049 | -0.113 | -0.594 [-1.186, -0.003] |
| SCP | .046 | -0.132 | -0.699 [-1.382, -0.015] |
| SLF_I | .021 | -0.124 | -0.695 [-1.274, -0.115] |
| SLF_II | .025 | -0.127 | -0.671 [-1.252, -0.090] |
| SLF_III | .023 | -0.125 | -0.700 [-1.297, -0.102] |
| STR | .040 | -0.116 | -0.639 [-1.248, -0.031] |
| UF | .275 | -0.031 | -0.347 [-0.985, 0.291] |
| T_PREM | .068 | -0.094 | -0.550 [-1.143, 0.042] |
| T_PAR | .031 | -0.120 | -0.672 [-1.280, -0.065] |
| T_OCC | .039 | -0.105 | -0.590 [-1.148, -0.032] |
| ST_FO | .078 | -0.051 | -0.520 [-1.103, 0.062] |
| ST_PREM | .037 | -0.108 | -0.651 [-1.259, -0.043] |
| CC_RG | .119 | -0.083 | -0.608 [-1.406, 0.191] |
| CC_body | .156 | -0.078 | -0.452 [-1.112, 0.209] |
| CC_IS | .034 | -0.118 | -0.641 [-1.221, -0.061] |

**Table S16.**
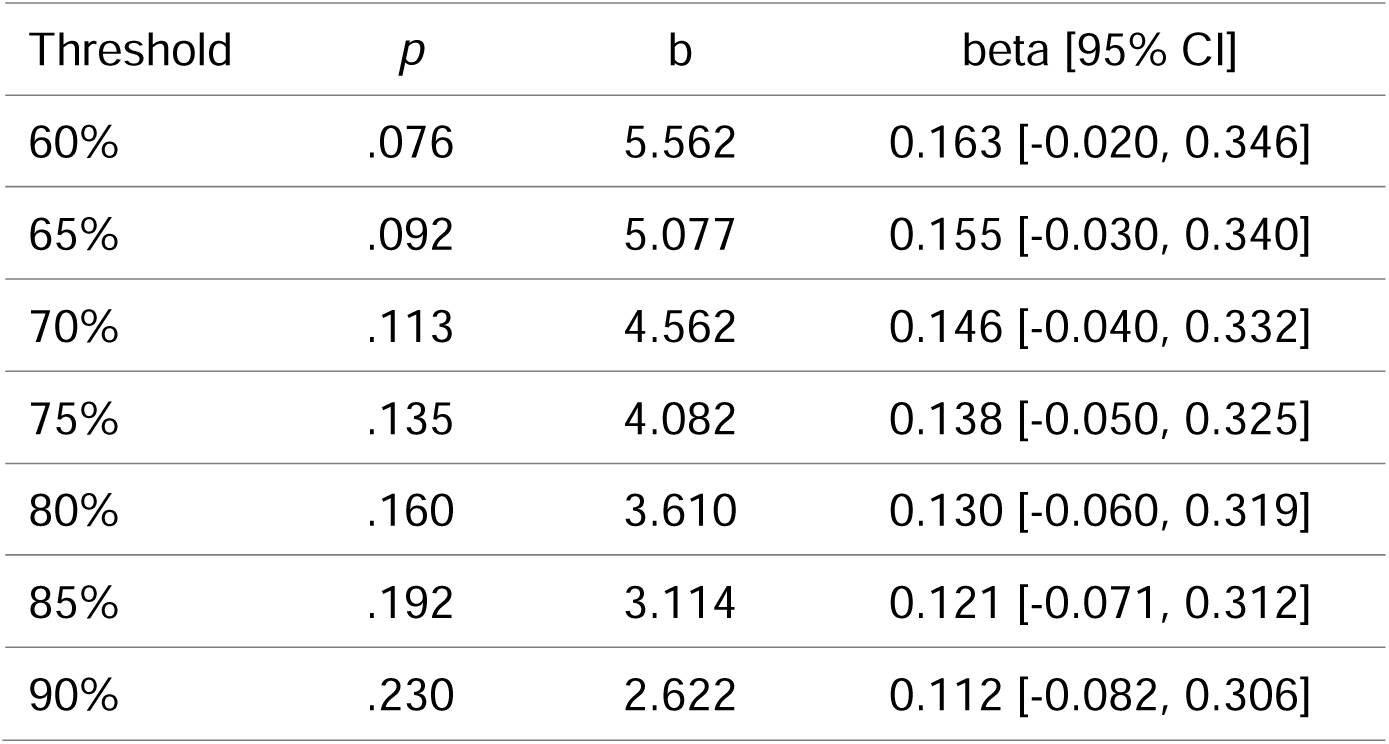
Statistical results of the linear mixed effects models for the relations between cerebrospinal fluid and salivary progesterone over the course of pregnancy with different voxel-level thresholds.

| Threshold | *p* | b | beta [95% CI] |
| --- | --- | --- | --- |
| 60% | .076 | 5.562 | 0.163 [-0.020, 0.346] |
| 65% | .092 | 5.077 | 0.155 [-0.030, 0.340] |
| 70% | .113 | 4.562 | 0.146 [-0.040, 0.332] |
| 75% | .135 | 4.082 | 0.138 [-0.050, 0.325] |
| 80% | .160 | 3.610 | 0.130 [-0.060, 0.319] |
| 85% | .192 | 3.114 | 0.121 [-0.071, 0.312] |
| 90% | .230 | 2.622 | 0.112 [-0.082, 0.306] |

**Table S17.** Statistical results of the linear mixed effects models for the relations between cerebrospinal fluid and salivary estradiol over the course of pregnancy with different voxel-level thresholds.

| Threshold | *p* | b | beta [95% CI] |
| --- | --- | --- | --- |
| 60% | .113 | 395.667 | 0.165 [-0.046, 0.375] |
| 65% | .114 | 378.352 | 0.164 [-0.046, 0.374] |
| 70% | .122 | 351.023 | 0.159 [-0.050, 0.369] |
| 75% | .130 | 325.897 | 0.156 [-0.054, 0.366] |
| 80% | .143 | 296.155 | 0.151 [-0.060, 0.361] |
| 85% | .153 | 267.516 | 0.147 [-0.064, 0.359] |
| 90% | .174 | 232.953 | 0.142 [-0.073, 0.356] |

**Table S18.**
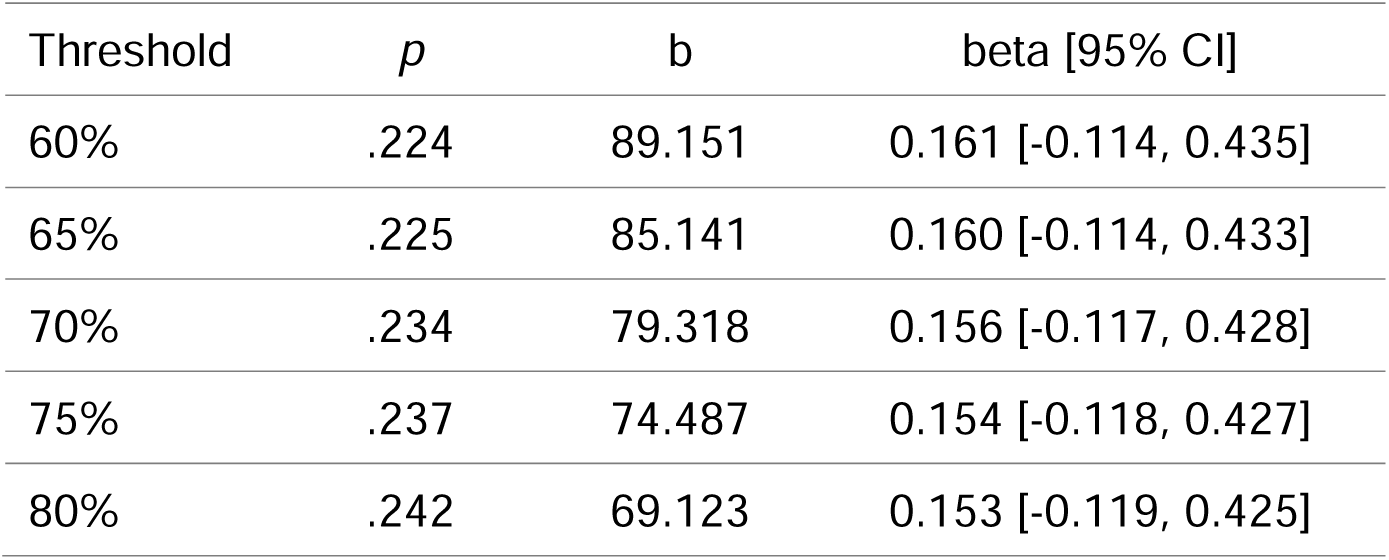
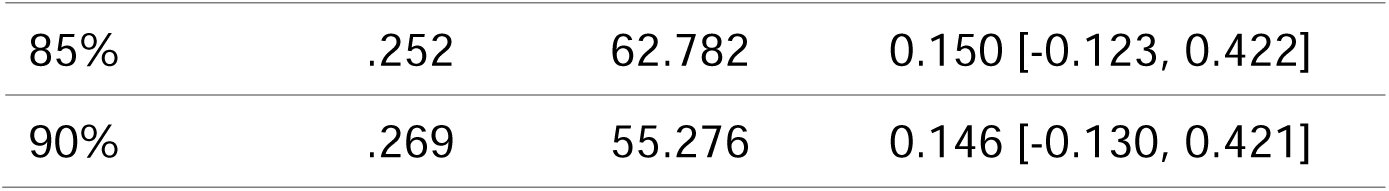
Statistical results of the linear mixed effects models for the relations between cerebrospinal fluid and salivary testosterone over the course of pregnancy with different voxel-level thresholds.

| Threshold | *p* | b | beta [95% CI] |
| --- | --- | --- | --- |
| 60% | .224 | 89.151 | 0.161 [-0.114, 0.435] |
| 65% | .225 | 85.141 | 0.160 [-0.114, 0.433] |
| 70% | .234 | 79.318 | 0.156 [-0.117, 0.428] |
| 75% | .237 | 74.487 | 0.154 [-0.118, 0.427] |
| 80% | .242 | 69.123 | 0.153 [-0.119, 0.425] |
| 85% | .252 | 62.782 | 0.150 [-0.123, 0.422] |
| 90% | .269 | 55.276 | 0.146 [-0.130, 0.421] |

**Table S19.** Statistical results of the linear mixed effects models for the relations between cerebrospinal fluid and salivary cortisol over the course of pregnancy with different voxel-level thresholds.

| Threshold | $p$ | b | beta [95% CI] |
| --- | --- | --- | --- |
| 60% | .285 | 15794.831 | 0.241 [-0.231, 0.713] |
| 65% | .319 | 14109.914 | 0.224 [-0.248, 0.697] |
| 70% | .346 | 12644.758 | 0.210 [-0.260, 0.681] |
| 75% | .376 | 11245.365 | 0.197 [-0.273, 0.668] |
| 80% | .401 | 9985.900 | 0.187 [-0.284, 0.658] |
| 85% | .423 | 8828.631 | 0.178 [-0.293, 0.650] |
| 90% | .441 | 7734.040 | 0.172 [-0.302, 0.647] |

**Table S20.**
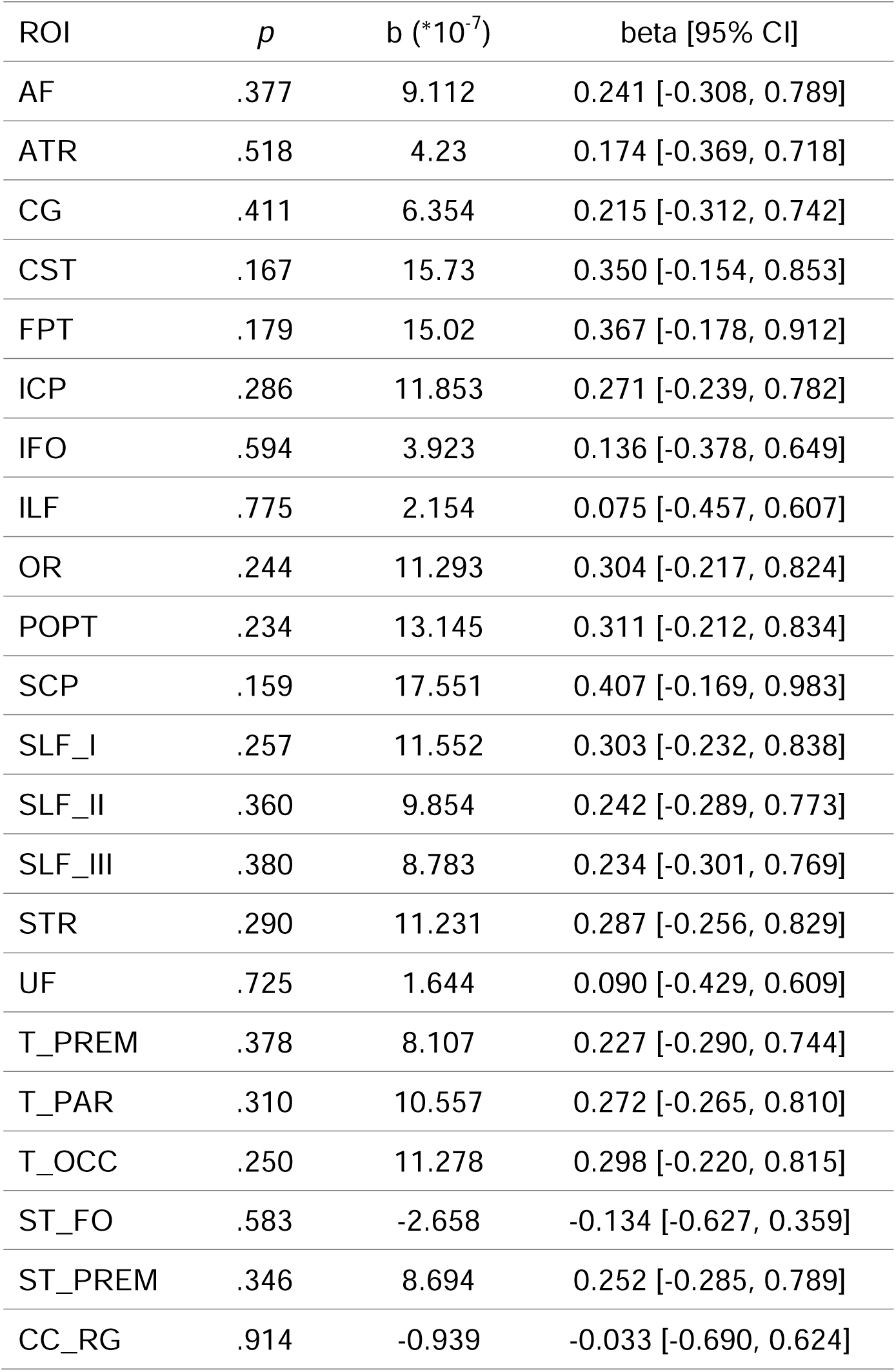
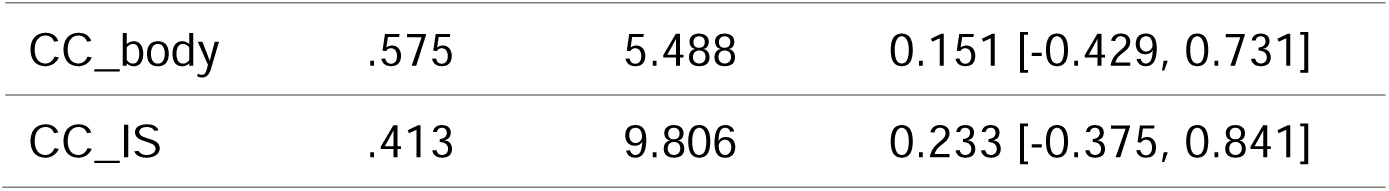
Statistical outcomes of the linear mixed effects models of the neurite density index in relation to total cerebrospinal fluid volume. AF: Arcuate Fascicle, ATR: Anterior Thalamic Radiation, CG: Cingulum, CST: Corticospinal Tract, FPT: Fronto-Pontine Tract, ICP: Inferior Cerebellar Peduncle, IFO: Inferior Occipito-Frontal Fascicle, ILF: Inferior Longitudinal Fascicle, OR: Optic Radiation, POPT: Parieto-Occipital Pontine, SCP: Superior Cerebellar Peduncle, SLF_I: Superior Longitudinal Fascicle I; SLF_II: Superior Longitudinal Fascicle II, SLF_III: Superior Longitudinal Fascicle III, STR: Superior Thalamic Radiation, UF: Uncinate Fascicle, T_PREM: Thalamo-Premotor, T_PAR: Thalamo-Parietal, T_OCC: Thalamo-Occipital, ST_FO: Striato-Fronto-Orbital, ST_PREM: Striato-Premotor, CC_RG: Rostrum and Genu, CC_body: Corpus Callosum Body, CC_IS: Isthmus and Splenium. ROI: region of interest.

| ROI | *p* | b (*10<sup>-7</sup>) | beta [95% CI] |
| --- | --- | --- | --- |
| AF | .377 | 9.112 | 0.241 [-0.308, 0.789] |
| ATR | .518 | 4.23 | 0.174 [-0.369, 0.718] |
| CG | .411 | 6.354 | 0.215 [-0.312, 0.742] |
| CST | .167 | 15.73 | 0.350 [-0.154, 0.853] |
| FPT | .179 | 15.02 | 0.367 [-0.178, 0.912] |
| ICP | .286 | 11.853 | 0.271 [-0.239, 0.782] |
| IFO | .594 | 3.923 | 0.136 [-0.378, 0.649] |
| ILF | .775 | 2.154 | 0.075 [-0.457, 0.607] |
| OR | .244 | 11.293 | 0.304 [-0.217, 0.824] |
| POPT | .234 | 13.145 | 0.311 [-0.212, 0.834] |
| SCP | .159 | 17.551 | 0.407 [-0.169, 0.983] |
| SLF_I | .257 | 11.552 | 0.303 [-0.232, 0.838] |
| SLF_II | .360 | 9.854 | 0.242 [-0.289, 0.773] |
| SLF_III | .380 | 8.783 | 0.234 [-0.301, 0.769] |
| STR | .290 | 11.231 | 0.287 [-0.256, 0.829] |
| UF | .725 | 1.644 | 0.090 [-0.429, 0.609] |
| T_PREM | .378 | 8.107 | 0.227 [-0.290, 0.744] |
| T_PAR | .310 | 10.557 | 0.272 [-0.265, 0.810] |
| T_OCC | .250 | 11.278 | 0.298 [-0.220, 0.815] |
| ST_FO | .583 | -2.658 | -0.134 [-0.627, 0.359] |
| ST_PREM | .346 | 8.694 | 0.252 [-0.285, 0.789] |
| CC_RG | .914 | -0.939 | -0.033 [-0.690, 0.624] |
| CC_body | .575 | 5.488 | 0.151 [-0.429, 0.731] |
| CC_IS | .413 | 9.806 | 0.233 [-0.375, 0.841] |

**Table S21.**
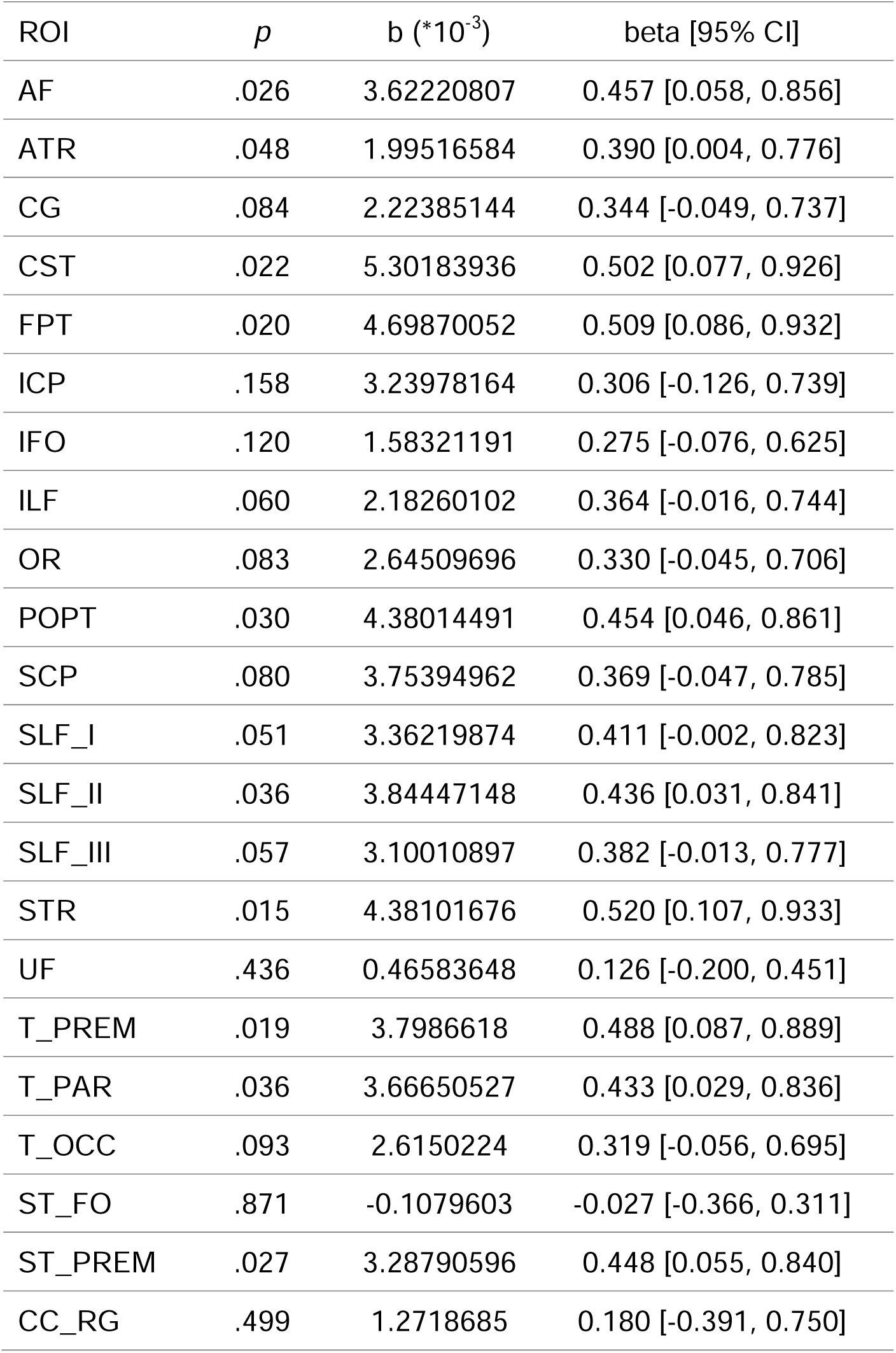
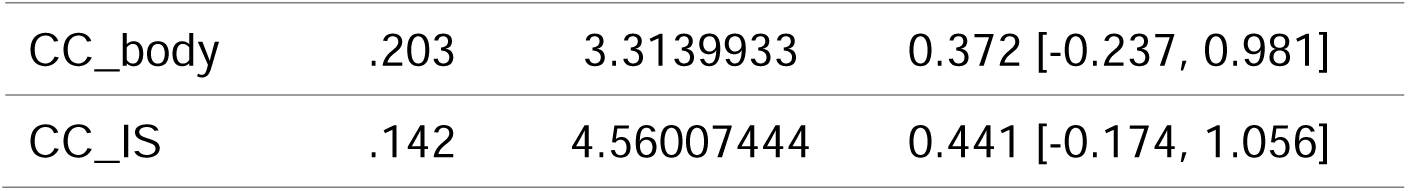
Statistical outcomes of the linear mixed effects models of the neurite density index in relation to hair cortisol. AF: Arcuate Fascicle, ATR: Anterior Thalamic Radiation, CG: Cingulum, CST: Corticospinal Tract, FPT: Fronto-Pontine Tract, ICP: Inferior Cerebellar Peduncle, IFO: Inferior Occipito-Frontal Fascicle, ILF: Inferior Longitudinal Fascicle, OR: Optic Radiation, POPT: Parieto-Occipital Pontine, SCP: Superior Cerebellar Peduncle, SLF_I: Superior Longitudinal Fascicle I; SLF_II: Superior Longitudinal Fascicle II, SLF_III: Superior Longitudinal Fascicle III, STR: Superior Thalamic Radiation, UF: Uncinate Fascicle, T_PREM: Thalamo-Premotor, T_PAR: Thalamo-Parietal, T_OCC: Thalamo-Occipital, ST_FO: Striato-Fronto-Orbital, ST_PREM: Striato-Premotor, CC_RG: Rostrum and Genu, CC_body: Corpus Callosum Body, CC_IS: Isthmus and Splenium. ROI: region of interest.

| ROI | *p* | *b* (*10<sup>-3</sup>) | beta [95% CI] |
| --- | --- | --- | --- |
| AF | .026 | 3.62220807 | 0.457 [0.058, 0.856] |
| ATR | .048 | 1.99516584 | 0.390 [0.004, 0.776] |
| CG | .084 | 2.22385144 | 0.344 [-0.049, 0.737] |
| CST | .022 | 5.30183936 | 0.502 [0.077, 0.926] |
| FPT | .020 | 4.69870052 | 0.509 [0.086, 0.932] |
| ICP | .158 | 3.23978164 | 0.306 [-0.126, 0.739] |
| IFO | .120 | 1.58321191 | 0.275 [-0.076, 0.625] |
| ILF | .060 | 2.18260102 | 0.364 [-0.016, 0.744] |
| OR | .083 | 2.64509696 | 0.330 [-0.045, 0.706] |
| POPT | .030 | 4.38014491 | 0.454 [0.046, 0.861] |
| SCP | .080 | 3.75394962 | 0.369 [-0.047, 0.785] |
| SLF_I | .051 | 3.36219874 | 0.411 [-0.002, 0.823] |
| SLF_II | .036 | 3.84447148 | 0.436 [0.031, 0.841] |
| SLF_III | .057 | 3.10010897 | 0.382 [-0.013, 0.777] |
| STR | .015 | 4.38101676 | 0.520 [0.107, 0.933] |
| UF | .436 | 0.46583648 | 0.126 [-0.200, 0.451] |
| T_PREM | .019 | 3.7986618 | 0.488 [0.087, 0.889] |
| T_PAR | .036 | 3.66650527 | 0.433 [0.029, 0.836] |
| T_OCC | .093 | 2.6150224 | 0.319 [-0.056, 0.695] |
| ST_FO | .871 | -0.1079603 | -0.027 [-0.366, 0.311] |
| ST_PREM | .027 | 3.28790596 | 0.448 [0.055, 0.840] |
| CC_RG | .499 | 1.2718685 | 0.180 [-0.391, 0.750] |
| CC_body | .203 | 3.3139933 | 0.372 [-0.237, 0.981] |
| CC_IS | .142 | 4.56007444 | 0.441 [-0.174, 1.056] |

**Table S22.**
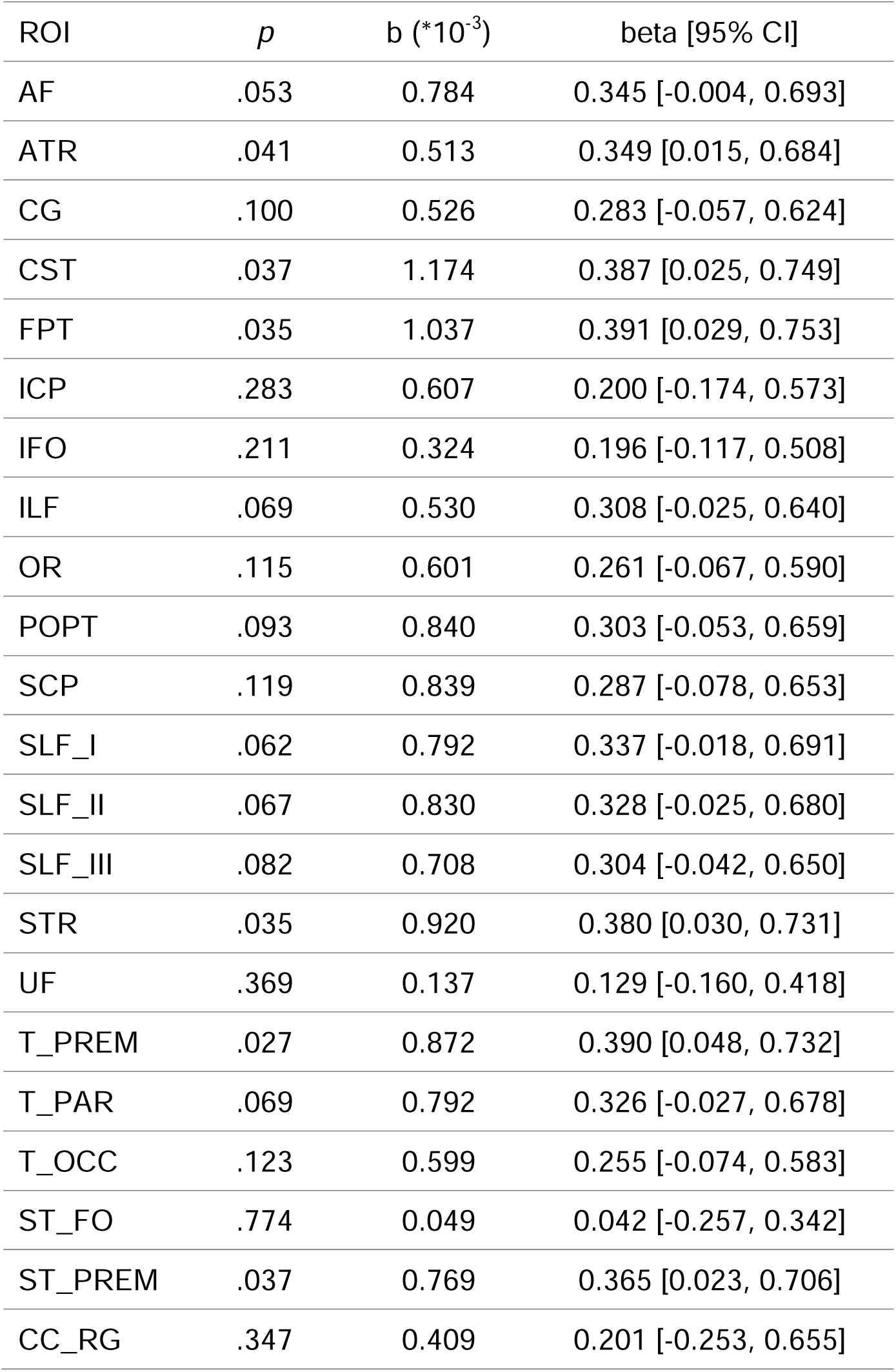
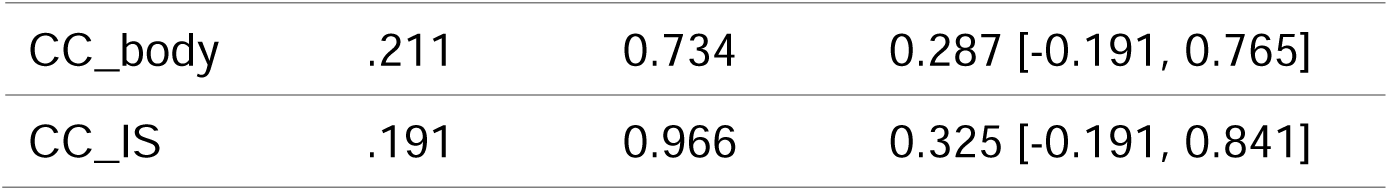
Statistical outcomes of the linear mixed effects models of the neurite density index in relation to hair cortisone. AF: Arcuate Fascicle, ATR: Anterior Thalamic Radiation, CG: Cingulum, CST: Corticospinal Tract, FPT: Fronto-Pontine Tract, ICP: Inferior Cerebellar Peduncle, IFO: Inferior Occipito-Frontal Fascicle, ILF: Inferior Longitudinal Fascicle, OR: Optic Radiation, POPT: Parieto-Occipital Pontine, SCP: Superior Cerebellar Peduncle, SLF_I: Superior Longitudinal Fascicle I; SLF_II: Superior Longitudinal Fascicle II, SLF_III: Superior Longitudinal Fascicle III, STR: Superior Thalamic Radiation, UF: Uncinate Fascicle, T_PREM: Thalamo-Premotor, T_PAR: Thalamo-Parietal, T_OCC: Thalamo-Occipital, ST_FO: Striato-Fronto-Orbital, ST_PREM: Striato-Premotor, CC_RG: Rostrum and Genu, CC_body: Corpus Callosum Body, CC_IS: Isthmus and Splenium. ROI: region of interest.

| ROI | *p* | b (*10<sup>-3</sup>) | beta [95% CI] |
| --- | --- | --- | --- |
| AF | .053 | 0.784 | 0.345 [-0.004, 0.693] |
| ATR | .041 | 0.513 | 0.349 [0.015, 0.684] |
| CG | .100 | 0.526 | 0.283 [-0.057, 0.624] |
| CST | .037 | 1.174 | 0.387 [0.025, 0.749] |
| FPT | .035 | 1.037 | 0.391 [0.029, 0.753] |
| ICP | .283 | 0.607 | 0.200 [-0.174, 0.573] |
| IFO | .211 | 0.324 | 0.196 [-0.117, 0.508] |
| ILF | .069 | 0.530 | 0.308 [-0.025, 0.640] |
| OR | .115 | 0.601 | 0.261 [-0.067, 0.590] |
| POPT | .093 | 0.840 | 0.303 [-0.053, 0.659] |
| SCP | .119 | 0.839 | 0.287 [-0.078, 0.653] |
| SLF_I | .062 | 0.792 | 0.337 [-0.018, 0.691] |
| SLF_II | .067 | 0.830 | 0.328 [-0.025, 0.680] |
| SLF_III | .082 | 0.708 | 0.304 [-0.042, 0.650] |
| STR | .035 | 0.920 | 0.380 [0.030, 0.731] |
| UF | .369 | 0.137 | 0.129 [-0.160, 0.418] |
| T_PREM | .027 | 0.872 | 0.390 [0.048, 0.732] |
| T_PAR | .069 | 0.792 | 0.326 [-0.027, 0.678] |
| T_OCC | .123 | 0.599 | 0.255 [-0.074, 0.583] |
| ST_FO | .774 | 0.049 | 0.042 [-0.257, 0.342] |
| ST_PREM | .037 | 0.769 | 0.365 [0.023, 0.706] |
| CC_RG | .347 | 0.409 | 0.201 [-0.253, 0.655] |
| CC_body | .211 | 0.734 | 0.287 [-0.191, 0.765] |
| CC_IS | .191 | 0.966 | 0.325 [-0.191, 0.841] |

